# Post-translational epigenetics: PRMT7 regulates RNA-binding capacity and protein stability to control *Leishmania* parasite virulence

**DOI:** 10.1101/736736

**Authors:** Tiago R. Ferreira, Adam A. Dowle, Ewan Parry, Eliza V. C. Alves-Ferreira, Foteini Kolokousi, Tony R. Larson, Michael J. Plevin, Angela K. Cruz, Pegine B. Walrad

**Affiliations:** York Biomedical Research Institute, Department of Biology, University of York, York, UK; Metabolomics and Proteomics Laboratory, Bioscience Technology Facility, Department of Biology, University of York, UK; Cell and Molecular Biology Department, Ribeirão Preto Medical School, University of São Paulo, Ribeirão Preto, Brazil

**Keywords:** *Leishmania*, arginine methylation, methyltransferase, PRMT, RBP, RNA binding

## Abstract

RNA binding proteins (RBPs) are the primary gene regulators in kinetoplastids as transcriptional control is nearly absent, making *Leishmania* an exceptional model for investigating methylation of non-histone substrates. Arginine methylation is an evolutionarily conserved protein modification catalyzed by Protein aRginine MethylTransferases (PRMTs). The chromatin modifier PRMT7 is the only Type III PRMT found in higher eukaryotes and a restricted number of unicellular eukaryotes. In *Leishmania major*, PRMT7 is a cytoplasmic protein implicit in pathogenesis with unknown substrates. Using comparative methyl-SILAC proteomics for the first time in protozoa, we identified 40 putative targets, including 17 RBPs hypomethylated upon PRMT7 knockout. PRMT7 can modify Alba3 and RBP16 *trans*-regulators (mammalian RPP25 and YBX2 homologs, respectively) as direct substrates *in vitro*. The absence of PRMT7 levels *in vivo* selectively reduces Alba3 mRNA-binding capacity to specific target transcripts and can impact the relative stability of RBP16 in the cytoplasm. RNA immunoprecipitation analyses demonstrate PRMT7-dependent methylation promotes Alba3 association with select target transcripts and stability of *δ-amastin* surface antigen. These results highlight a novel role for PRMT7-mediated arginine methylation upon RBP substrates, suggesting a regulatory pathway controlling gene expression and virulence in *Leishmania*. This work introduces *Leishmania* PRMTs as epigenetic regulators of mRNA metabolism with mechanistic insight into the functional manipulation of RBPs by methylation.

## INTRODUCTION

Protein arginine methyltransferases (PRMTs) are widely distributed across eukaryotes in 11 different classes of enzymes (PRMT1-11). They catalyze arginine methylation in multiple cellular processes including histone modification, transcription control, RNA processing, protein localization and cell signaling (1) (2–4). Histones and RNA-binding proteins (RBPs) have been characterized as PRMT substrates in different organisms (1).

PRMTs are classified as Type I, II or III according to the targeted nitrogen and number of methyl groups transferred (Figure 1B). PRMT7 is a unique enzyme as it is the sole PRMT known to catalyze only monomethyl arginine (MMA; Type III PRMT), which may indicate a regulatory step that primes substrates prior to dimethylation (5–7). PRMT7 has been well described in mammalian cells as a histone methyltransferase which modulates chromatin to repress gene promoters. Remarkably, the only unicellular eukaryotes known to carry a PRMT7 homolog are choanoflagellates and trypanosomatids, including *Leishmania*, where gene expression control is primarily post-transcriptional (8). The heightened regulatory role of RBPs in *Leishmania* can lend clear functional insight not obscured by complex networks of transcriptional regulation (9).

**Figure 1.**
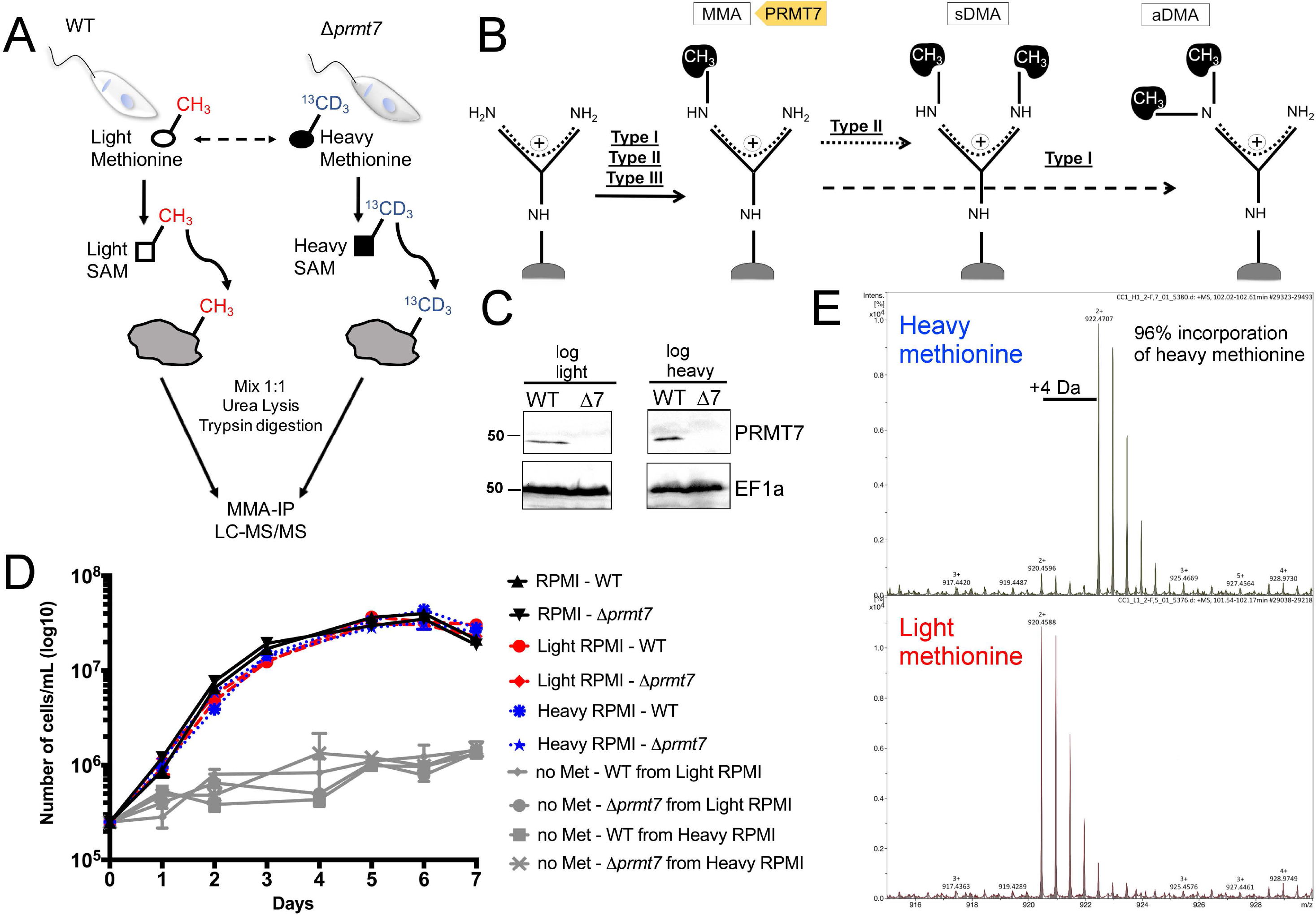
Heavy methyl SILAC is a feasible approach in *Leishmania major* promastigotes and does not significantly affect promastigote culture growth. (A) *L. major* wild-type (WT) and PRMT7 knockout (Δ*prmt7*) parasites were kept in SILAC media in the presence of light or heavy methionine (L-Methionine-methyl-13C,d3) with label-swap replications. Tryptic peptides were submitted to anti-monomethyl arginine immunoprecipitation and LC-MS/MS for identification and quantification of methyl peptides. (B) PRMT7 is the single Type III methyltransferase identified to date and thus catalyzes only MMA. Type II and Type I PRMTs catalyze, respectively, symmetric dimethylarginine (sDMA) and asymmetric dimethylarginine (aDMA), in addition to MMA in a first step. (C) Expression of PRMT7 in logarithmic growth phase promastigotes was evaluated by immunoblotting for comparison between light and heavy SILAC media, which showed no significant difference. EF1a levels are shown for protein loading control. Blots are representative of a biological duplicate. (D) WT and Δ*prmt7* promastigotes growth curves were not significantly altered in different SILAC media (light or heavy RPMI). If methionine is not added to the culture media (no Met) parasite growth is not significant, which confirms the successful removal of methionine from the dialyzed fetal bovine serum. Data are plotted as mean ± standard error from three biological replicates. (E) MS spectra of a representative WT-derived peptide showing 96% incorporation of heavy methionine.

*Leishmania* spp. parasites are the causative agent of the leishmaniases, infectious diseases that contribute the ninth largest global disease burden, with 1 million new cases diagnosed annually (10). We have previously shown that *Leishmania major* PRMT7 expression is stage-regulated during development, regulates host pathology and localizes to the cytoplasm unlike the nuclear-specific mammalian PRMT7 (11). Remarkably, the *Leishmania* PRMT7-directed regulatory pathway epigenetically controls parasite gene expression, and resultant pathology outcomes 4 cellular differentiation events and 4 months after PRMT7 protein expression.

The impact of PRMT7 on the overall MMA-modified proteome is still unknown. To date, histones are the only mammalian PRMT7 targets validated *in vivo* (6,12). In contrast, *L. major* PRMT7 displays distinct localization, low sequence similarity and is nonessential. Combined with the negligible transcriptional control in *Leishmania* spp., all evidence suggests a divergent function between human and *Leishmania* PRMT7 enzyme (11). A recent large-scale study compared the distribution of arginine residues across eukaryotes and found that unicellular organisms that carry a PRMT7 homolog (including *Monosiga brevicolis* and *Leishmania infantum*) have a higher ratio of predicted methylarginine/unmodified arginine than others (*Plasmodium falciparum* and *Dictyostelium discoideum*) (4). Indeed both *Leishmania major* and *Trypanosoma brucei* PRMT7s are distinct enough from mammalian orthologs that the regulatory pathways they drive may present anti-parasitic drug targets or an opportunity to repurpose existing drugs (5,11,12). We previously demonstrated that PRMT7 levels are inversely proportional to *L. major* parasite virulence, functionally linking this enzymatic pathway to leishmaniasis pathogenesis (11).

Here, we present the first comprehensive investigation of the molecular impact of modulating PRMT7 levels and the first quantified identification of a *Leishmania* arginine monomethyl proteome. Using methyl-SILAC proteomics, we identified 247 arginine monomethylated proteins, including 62 RNA-binding proteins (RBPs); 17 of which are hypomethylated upon PRMT7 knockout. RBPs are the primary genetic regulators in *Leishmania* species as genes are constitutively transcribed (9). We further validate PRMT7 methylation targets both *in vitro* and *in vivo* and reveal arginine methylation as an important post-translational regulator of RBP function with differential impact upon individual *trans*-regulators. PRMT7-dependent monomethylation promotes cytoplasmic RBP16 protein stability as well as controls the highly selective mRNA binding capacity of Alba3; stabilizing *δ-amastin* surface antigen mRNA levels but not impacting association or stability of *nmt* transcript target. The near-absence of transcriptional control in *Leishmania* amplifies the regulatory role of *trans*-regulators and renders it an excellent system to investigate the impact of PTMs on RBP function (8). This study introduces *Leishmania* PRMTs as epigenetic regulators of downstream parasite virulence via modified RBP protein expression, selective RNA binding capacity and subsequent mRNA metabolism.

## MATERIAL AND METHODS

### *Leishmania major* cultures and transfection

*Leishmania major* promastigotes strain CC1 (MHOM/IR/83/LT252) were cultured at 26°C in M199 medium supplemented with 40 mM HEPES, 10% FBS, 100 U penicillin/mL, 100 μg/mL, 100 μM adenine, 0.0005% Hemin. In the phlebotomine vector, *L. major* parasites differentiate from the proliferative non-infective procyclic promastigotes to the non-dividing human-infective metacyclic promastigotes, a metacyclogenesis process that is mimicked in culture (13). Procyclics and metacyclics are enriched in axenic culture, respectively, at log and stationary growth phases. Inside mammalian phagocytic cells (primarily macrophages) metacyclic promastigotes differentiate into non-motile amastigotes.

*PRMT7* null parasites (Δ*prmt7*) were previously generated (11). Plasmid and PCR oligos used in this study are shown in Supplementary Figure S2 and Table S3, respectively. The coding sequence (CDS) and 500bp 5’ flanking region (FLR) of RNA-binding proteins Torus (LmjF.36.0740), DDX3 (LmjF.32.0400), RBP16 (LmjF.28.0825) and Alba3 (LmjF.34.2580) were PCR amplified. The cloning strategy involved using four different SfiI enzyme sites to allow a single-step four-way ligation: SfiA (GGCCACCTAGGCC), SfiB reverse (GGCCACGCAGGCC), SfiC (GGCCGCTGGGGCC) and SfiD reverse (GGCCTGACTGGCC)(14). PCR products were purified using NucleoSpin Gel and PCR Clean-up (Macherey-Nagel), cloned into pGEM^®^Teasy vector (Promega), SfiI digested and ligated into plasmid pFLAG_HA-RBP (Supplementary Figure S2A). Positive clones were confirmed by diagnostic restriction and sequencing, PmeI-PacI digested overnight and purified DNA was transfected into *L. major* WT and Δ*prmt7* using Amaxa Nucleofector IIb (Lonza). Transfectants were selected on 1% agar M199 plates with 10 μg/mL G418 and PCR-validated using 5’FLR and 3’FLR primers (Supplementary Figures S2B and C).

Alba3 hypomethylated mutants were generated by replacing arginine residues (RG/RGG motifs) in the C-terminus to either tryptophan (WGG) or lysine (R192K or KGG) and inserting each construct into pFLAG_HA-Alba3. For the latter, *alba3* sequence was modified by Q5^®^ Site-Directed Mutagenesis (New England Biolabs), according to manufacturer instructions, using oligos shown in Table S3 to either replace a single arginine detected in the MMA methyl-SILAC analysis (Alba3^R192K^) or all the eleven arginine residues in RGG motifs present in its C-terminal tail (Alba3^KGG^). The Alba3^WGG^ mutant sequence was obtained as a custom synthetic gene (GenScript), in which each of the eleven arginine residues were mutated. Final constructs were transfected into both WT and Δ*prmt7 L. major* cell lines.

### Parasite labeling by SILAC

Heavy methyl-SILAC has been shown to provide high confidence methyl peptide matches, and it is considered the optimal method for identifying protein methylation in large-scale proteomics (15). Parasites were cultured for 10 passages in SILAC-RPMI media (RPMI lacking Methionine (Dundee Cell Products), 10% 3.5 kDa-dialyzed serum, 20 mM HEPES pH 7.4, 100 μM adenine, 20 mM L-glutamine, 100 U/mL penicillin, 100 μg/mL streptomycin, 1 μM biopterin, 0.0005% hemin) supplemented with either L-methionine or L-methionine-methyl-^13^CD_3_ (Sigma-Aldrich), for 10 passages corresponding to approximately 100 parasite divisions. L-methionine is converted into S-adenosylmethionine (SAM), the main biological methyl donor for transmethylation reactions, which allows direct labelling of methylated substrates (3). Next, J774.2 macrophages kept in SILAC-RPMI media for 4 passages were infected with late-stationary phase promastigotes at a 10:1 ratio to obtain recently differentiated promastigotes, as prolonged axenic promastigote culturing can alter virulence-related features (16). Three hours post-infection, plates were washed in incomplete RPMI and kept for 48 h under the same conditions. Amastigotes were purified and differentiated into procyclic promastigotes in SILAC-RPMI at 26°C and cultured for 3 passages.

### Parasite protein extraction and enrichment of monomethylated arginine peptides

Logarithmic procyclic promastigotes (10^9^ parasites) were pelleted at 2,000 g for 10 min, washed in PBS and lysed (20 mM HEPES pH 8.0, 9 M urea, 1 mM sodium orthovanadate, 2.5 mM sodium pyrophosphate, 1 mM β-glycerophosphate) via three rounds of sonication at 15 W for 15 s at room temperature (RT) to avoid urea precipitation with 1 min intervals on ice. Lysates were cleared at 20,000 g, 20 min, 15°C and supernatants were collected.

Sample preparation and enrichment of monomethyl arginine peptides were conducted using PTMScan[mme-RG] kit (Cell Signaling Technology) according to manufacturer’s protocols with the following modifications: proteins were reduced in 4.5 μM DTT at 55°C for 30 min, cooled on ice and alkylated in 2 mg/mL iodoacetamide at RT for 15 min. Lysates were diluted 3-fold in 20 mM HEPES pH 8.0 and digested overnight in 10 μg/mL trypsin-TPCK (Sigma-Aldrich) and 10 μM HCl, at RT under agitation. Digestion was confirmed by SDS-PAGE and Coomassie Blue R staining. Digests were acidified with trifluoroacetic acid (17) to a final concentration of 0.1% and a pH strip was used to confirm a pH < 3. Precipitate was formed at RT for 15 min. Peptides were purified using a Sep-Pak C_18_ column (Waters) connected to a 10 cc reservoir, pre-wet in 5 mL 100% acetonitrile and washed sequentially in 1, 3 and 6 mL 0.1% TFA. Acidified peptide solution was cleared at 1,800 g for 15min at RT and cleared supernatant was loaded onto the C_18_ column using a 10 cc plunger. The column was washed sequentially in 1, 5 and 6 mL 0.1% TFA and then in 2 mL 5% acetonitrile, 0.1% TFA. Peptides were collected by 3 elutions in 2 mL 40% acetronitrile, 0.1% TFA and dried in a Speed-Vac concentrator for approximately 3 h.

Dry peptide pellets were dissolved in 1.4 mL Immunoaffinity Purification (IAP) buffer (CST) and pH ∼7.0 was confirmed on a pH strip. All subsequent steps were performed on ice or at 4°C. Peptide solution was cleared at 10,000 g for 5 min and supernatant was transferred to a fresh tube. Anti-MMA agarose beads from the PTMScan kit were washed 4 times at 2,000 g for 1 min, in 1 mL PBS (8 mM Na_2_HPO_4_; 1.5 mM NaH_2_PO_4_; 2.7 mM KCl; 137 mM NaCl; pH 7.0) and resuspended in 40 μL PBS. Peptide solution was mixed with the anti-MMA beads and incubated for 2 h under rotation. Peptide-beads slurry was centrifuged at 2,000 g for 1 min and unbound peptides were collected and kept at −80°C. Beads were washed twice in 1 mL IAP buffer and three times in HPLC-grade water; inverting the tube 5 times and centrifuged at 2,000 g for 1 min. MMA peptides were collected by 2 elutions in 55 μL of 0.15% TFA at RT for 10 min, mixing every 2 min, and centrifuged at 2,000 g for 1 min.

Eluted monomethyl peptides were concentrated and purified using a Ziptip (Merck-Millipore). The Ziptip was first equilibrated in 50 μL of 50% acetronitrile, 0.1% TFA and twice in 50 μL of 0.1% TFA. IP eluent was loaded by passing through the Ziptip 10 times. Tips were washed twice in 0.1% TFA and peptides were eluted in 10 μL 40% acetonitrile, 0.1% TFA at least 10 times. Ziptip eluent was precipitated in a Speed-Vac and MMA peptides were resuspended in 10 μL of 0.1% TFA.

### Mass spectrometry data acquisition and analysis

Samples were loaded onto an UltiMate 3000 RSLCnano HPLC system (Thermo) equipped with a PepMap 100 Å C_18_, 5 μm trap column (300 μm × 5 mm, Thermo) and a PepMap, 2 μm, 100 Å, C_18_ EasyNano nanocapillary column (75 μm × 150 mm, Thermo). The trap wash solvent was 0.05% (v:v) aqueous trifluoroacetic acid and the trapping flow rate was 15 μL/min. The trap was washed for 3 min before switching flow to the capillary column. Separation used gradient elution of two solvents: solvent A, aqueous 1% (v:v) formic acid; solvent B, aqueous 80% (v:v) acetonitrile containing 1% (v:v) formic acid. The flow rate for the capillary column was 300 nL/min and the column temperature was 40°C. The linear multi-step gradient profile was: 3-10% B over 8 min, 10-35% B over 125 min, 35-65% B over 50 min, 65-99% B over 7 min and then proceeded to wash with 99% solvent B for 4 min. The column was returned to initial conditions and re-equilibrated for 15 min before subsequent injections.

The nanoLC system was interfaced with an Orbitrap Fusion hybrid mass spectrometer (Thermo) with an EasyNano ionisation source (Thermo). Positive ESI-MS and MS^2^ spectra were acquired using Xcalibur software (version 4.0, Thermo). Instrument source settings were: ion spray voltage, 1,900 V; sweep gas, 0 Arb; ion transfer tube temperature; 275°C. MS^1^ spectra were acquired in the Orbitrap with: 120,000 resolution, scan range: *m/z* 375-1,500; AGC target, 4e^5^; max fill time, 100 ms. Data dependant acquisition was performed in top speed mode using a 1 s cycle, selecting the most intense precursors with charge states 2-5. Dynamic exclusion was performed for 50s post precursor selection and a minimum threshold for fragmentation was set at 3e^4^. MS^2^ spectra were acquired in the linear ion trap with: scan rate, rapid; quadrupole isolation, 1.6 *m/z*; activation type, HCD; activation energy: 32%; AGC target, 5e^3^; first mass, 110*m/z*; max fill time, 100 ms. Acquisitions were arranged by Xcalibur to inject ions for all available parallelizable time.

### Peptide Identification

Protein identification was performed using Sequest HT and Mascot .Thermo .raw files were submitted to Sequest HT database searching using Proteome Discoverer (version 2.1, Thermo). Peak lists were extracted following .raw to .mgf format conversion using MSconvert (version 3.0.9967, ProteoWizard) before submitting via Mascot Daemon (version 2.5.1, Matrix Science Ltd.) to a local-running copy of the Mascot program (version 2.5.1, Matrix Science Ltd.). Database searching was performed against the TriTrypDB (version 8.1)(18) *Leishmania major* Friedlin genome (8,400 sequences). Search criteria specified: Enzyme, trypsin; Fixed modifications, Carbamidomethyl (C); Variable modifications, Oxidation (M), Phospho (ST), Methyl (R), Methyl (K), Methyl:2H(7)13C(1) (R), Label:13C(1)2H(7) (M), Label:13C(1)2H(7) Oxidation (M); Peptide tolerance, 5 ppm; MS/MS tolerance, 0.5 Da; Maximum missed cleavages, 3; Instrument, ESI-TRAP. Scaffold PTM (version 3.0.0, Proteome Software) was used to calculate methylation site localization probabilities for Mascot derived peptide identifications using the algorithm as described (19). Briefly, MS^2^ spectra identified as originating from methylated peptides were re-analyzed to calculate Ascore values and site localization probabilities to assess the level of confidence in each PTM positional assignment. Scaffold PTM then combined localization probabilities for all peptides containing each identified PTM site to obtain the best estimate probability that a PTM is present at that particular site.

### Relative Quantification of Methylated Peptides

A list of unique, methylated peptides was compiled from Sequest and Mascot identifications across all samples post-filtering to require posterior error probabilities <0.05. The list was used to calculate theoretical *m/z* values for the light and heavy analogues of all identified methylated peptides. Observed retention times from search results were associated with theoretical *m/z* pairs and appended to the mass list to create an extracted ion chromatogram (XIC) target list. Where methylated peptides were identified multiple times the median retention time was used. Thermo .raw files were converted to .mzML using Bioconductor (version 3.5) in R (version 3.3.1) before processing using the xcms package (20). XICs were extracted for all theoretical light and heavy *m/z* pairs across retention time aligned samples within 60 s windows, and peak areas extracted using the centWave algorithm (21). Light and heavy peak areas were normalized to relative percentages (light/heavy) and the reciprocal percentage (heavy/light) calculated for replicates originating from switched light to heavy metabolic labelling. Significance testing for quantitative differences between normalized light and heavy peak areas was performed using Student’s t-test (two-tailed, heteroscedastic). Calculated p-values were multiple-test corrected using the Hochberg and Benjamini false discovery rate estimation. Accepted differences were required to have corrected p-values <0.05, be derived from XIC peak areas in >2 samples and be observed consistently in at least one replicate upon light to heavy metabolic label switching.

### Bioinformatic analyses of PRMT7 MMA RNA binding protein targets

Amino acid sequences of 18 PRMT7 targets were downloaded from TriTrypDB.org (18) and subjected to a set of bioinformatic tools to predict the type and location of constituent domains. Initial Pfam (22) domain annotations were obtained using BLASTP (23) and RefSeq and PDB sequence databases. Predictions of secondary structure and intrinsic disorder were performed using PSIPRED (24) and DISOPRED (25), respectively. Additional structural predictions were conducted using Phyre2 in standard mode (26). The results of these analyses were combined to inform manual predictions of likely domain boundaries and to annotate domain type and function. Extended regions of predicted secondary structure and low disorder that lacked a clear functional annotation were assigned as domains of unknown function (DUF).

### *In vitro* methylation assay by PRMT7

Recombinant proteins were expressed in *Escherichia coli* BL21(DE3)pLys using pET28a+ (Novagen) and purified using Ni-Sepharose affinity medium (GE Healthcare) as previously described (11). Purified and dialyzed proteins were analyzed on SDS-PAGE and had their identity verified by mass spectrometry. PRMT7 wild-type and mutants were incubated with putative substrates in PBS with 2 μCi of S-adenosyl-[methyl-3H]methionine (^[3H]^AdoMet) (55-85 Ci/mmol, PerkinElmer) for 18 h at 26°C. Reactions were then halted by the addition of 2X SDS sample buffer (125 mM Tris-Cl pH 6.8, 20% glycerol, 2% SDS, 0.7 M β-Mercaptoethanol and 0.01% bromophenol blue) at 95°C for 5 min. Reactions were loaded and run on 12.5% SDS-PAGE and gels were dried on Whatman paper in a vacuum (Gel Dryer 583, Bio-Rad) for 2 h at 80°C on a gradient drying cycle. The dried gel was exposed to Hyperfilm (GE Healthcare) for 3-14 days at −70°C. Gels were also stained with Coomassie R for loading control. Gels and films were scanned on an ImageScanner III (GE Healthcare) for posterior analysis using GIMP (GNU Image Manipulation Program).

### Immunoblotting

Promastigotes at log or stationary growth phase (10^7^ cells per lane) were sedimented at 2,000 g for 10 min, washed once in PBS under the same conditions, lysed in 2X SDS sample buffer and boiled 5 min before loading on 12.5% SDS-PAGE gels. Proteins were transferred to PVDF membranes at 20 V for 1 h using a Novex Semi-Dry blotter (Thermo). Blots were blocked in 5% milk TBS-T (20 mM Tris, 150 mM NaCl, pH 7.4, 0.05% Tween) for 1 h at RT, or 5% BSA TBS-T for anti-PRMT7 blots. Primary antibodies were diluted in 1% milk or BSA TBS-T accordingly: mouse anti-HA [1:20,000] (Thermo); rabbit anti-MMA [1:3,000] (MultiMab, CST); chicken anti-PRMT7 [1:1,000] (11), rabbit anti-*Tb*RBP16 [1:1,000] (27) and mouse anti-EF1a [1:200,000] (clone CBP-KK1, Merck-Millipore) for 18 h at 4°C. All subsequent steps were performed at RT. Blots were briefly washed twice, then washed three times in TBS-T for 10 min and incubated in the corresponding diluted secondary antibodies for 1 h at RT. Membranes were washed and incubated in ECL Prime (GE Healthcare) detection solutions for 5min prior to ECL Hyperfilm exposure (GE Healthcare). Films were developed, scanned and images were processed using GIMP.

### Immunofluorescence

Promastigotes were incubated with 200 nM of MitoTracker™ Green FM (Molecular Probe) for 30 minutes at 26°C. Cells were harvested at 2,000 g 10 min, washed in 1X PBS and fixed in 2% paraformaldehyde for 20 min. Fixed cells were centrifuged, suspended in 1 M glycine solution and attached to poly-lysine-coated coverslips. Permeabilization was performed using 0.2% Triton X-100 for 3 min, blocking was carried out in 2% BSA-PBS for 30 min, parasites were incubated in monoclonal anti-HA mouse [1:500] (Sigma) or rabbit anti-TbRBP16 [1:500] in 2% BSA-PBS for 1 h, washed 3 times in PBS, incubated for 30 min in anti-mouse or anti-rabbit Alexa Fluor 594 [1:1,000] (Invitrogen) in 2% BSA-PBS supplemented with 60 μM DAPI, washed in PBS 3 times and mounted with Fluoroshield™ (Sigma). Images were acquired on an Axio Observer microscope (Zeiss) and images were processed using ZEN 2.3 SP1 (black, Carl Zeiss).

### RNA immunoprecipitation and qRT-PCR

HA-Alba3 was immunoprecipitated from *in vivo* UV-crosslinked log promastigotes with anti-HA magnetic beads (Thermo) as described (9,28). Crosslinking was performed with UVC using the LT40 “Minitron” system (UV03 Ltd; (Granneman, 2010 #817)) for 120 s at 1.6mJ/cm^2^, which is less stressful to parasites and damaging to mRNA (9). Total RNA was extracted from WT and Δ*prmt7* input samples using Direct-zol RNA Miniprep (Zymo Research). cDNA was synthesized from 2 μg of total RNA or 100-400 ng of RIP RNA using the SuperScript IV Reverse Transcriptase (Invitrogen). Absolute quantification curves were performed for each oligonucleotide pair using serial dilutions of cDNA. Relative quantification (−ΔΔCt) was performed using the Fast SYBR Green Master Mix and Quantstudio 3 PCR System (Thermo Fisher). RNA levels were normalized to the 18S rRNA and glycosomal glyceraldehyde 3-phosphate dehydrogenase (GAPDHg, LmjF.30.2980). GeneIDs of the mRNAs are: *NIMA-related kinase* (LmjF.35.5190), *nmt* (LmjF.32.0080), *δ-amastin* (LmjF.34.0500) and *p1/s1 nuclease* (LmjF.30.1510). Oligonucleotides used are listed in Supplementary Table S3.

### Protein and mRNA decay

For protein stability evaluation, promastigotes at day 2 (log phase; non human-infective promastigote) or day 6 (stationary phase; human-infective promastigote) post-inoculum were incubated in 200 μg/mL cycloheximide at 26°C up to 24 h. Cells were harvested at specific time-points and resuspended in 2X Sample buffer (as above). A volume equivalent to 10^7^ cells/lane was loaded on 12.5% SDS-PAGE and assessed by immunoblot as described above. Membranes were stained post-transfer in Ponceau for loading controls. Densitometry analysis was performed using ImageJ software (29)

For transcript stability analysis, log phase promastigotes were incubated up to 4 h in 2.5 μg/mL sinefungin (Cayman Chemical) and 5 μg/mL actinomycin D (Sigma) to inhibit *trans*-splicing and *de novo* transcription (30). Sinefungin was added 15 min before actinomycin D. Cells (2×10^7^) were harvested at specific time points and lysed in Trizol reagent (Invitrogen). Total RNA was extracted using Direct-zol RNA Miniprep (Zymo Research). RNA levels were quantified by qRT-PCR as explained above and normalized to 18S rRNA.

## RESULTS

### RNA-binding proteins are the primary targets of arginine monomethylation in *L. major*

To evaluate the impact of PRMT7 loss upon the arginine monomethyl proteome of *Leishmania major*, wild-type (WT) and Δ*prmt7* parasites were labeled with either L-methionine (31) or L-methionine-methyl-^13^CD_3_ (heavy) and submitted for methyl-SILAC proteomic analysis (Figure 1A). Expression of PRMT7 was similar between light and heavy methyl-labeled parasites (Figure 1C) and no significant growth difference was observed between WT and Δ*prmt7* cells, light or heavy (Figure 1D). Heavy methionine incorporation of >95% was confirmed by LC-MS (Figure 1E), which is sufficient and essential for SILAC analyses (32).

Comprehensive peptide quantification by methyl-SILAC revealed 40 proteins hypomethylated in Δ*prmt7* cells (WT/Δ*prmt7* methyl peptide ratio >1.5) from the 247 MMA-containing proteins identified (≥ 2 peptides and detection in label-swap repetition; Figures 2A and 2B; Supplementary Table S1). In total, these proteins are represented by 387 unique MMA peptides identified. Mass spectrometry analysis of unfractionated peptides (input) did not show significant differential expression of total methylated proteins between wild-type and Δ*prmt7* samples.

**Figure 2.**
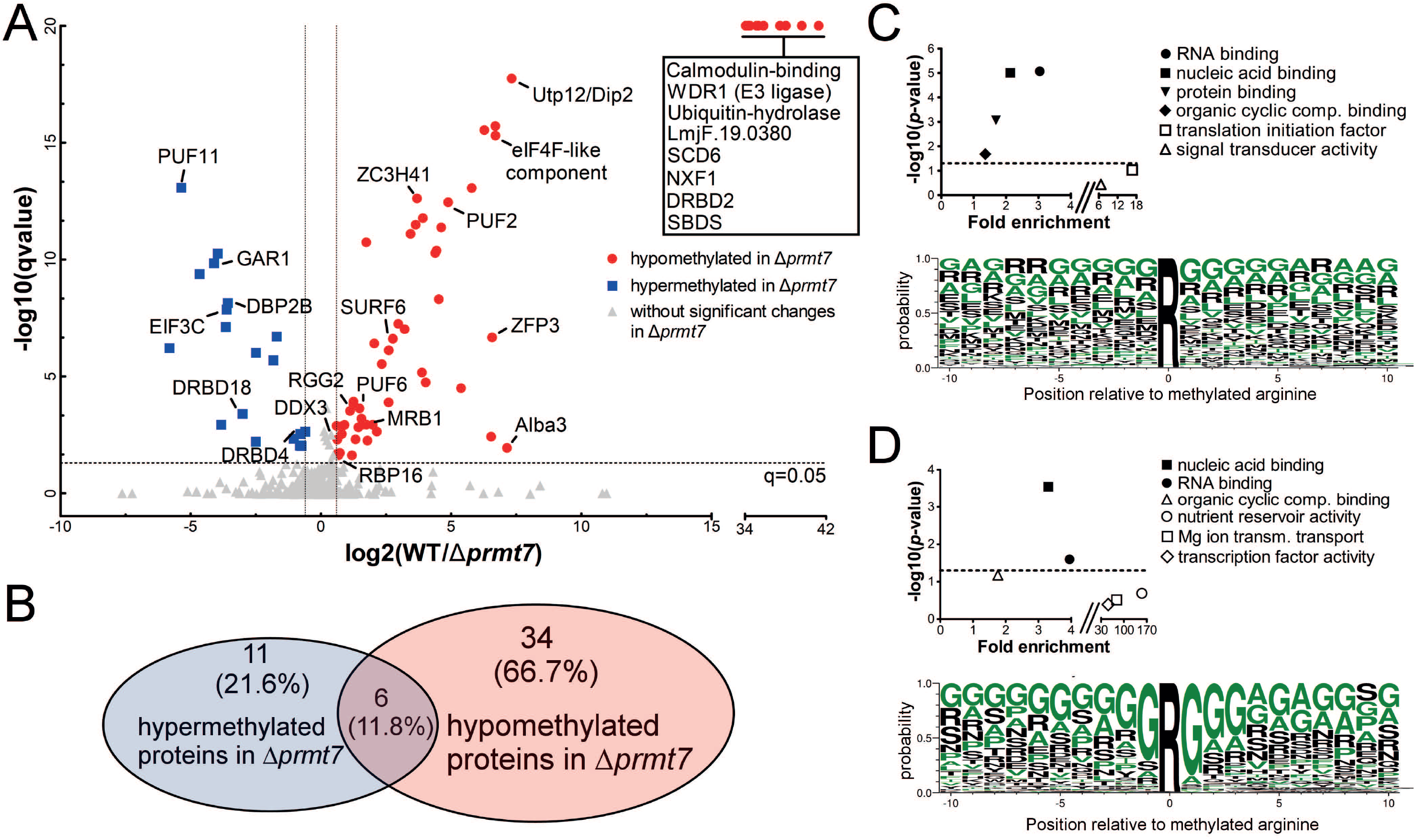
RNA-binding proteins are putative substrates for PRMT7 arginine monomethylation. (A) Methyl-SILAC peptide pairs were analyzed according to their methylation status in WT versus Δ*prmt7 Leishmania major*. Mass spectra quantification of each light and heavy pair ratios detected peptides with increased (hypermethylated, in blue) or decreased (hypomethylated, in red) methylation in Δ*prmt7* as compared to WT. Methylated RNA-binding proteins (RBPs) are shown as well as proteins that had no methyl peptides detected in Δ*prmt7* (boxed RBPs). Of 247 identified proteins with isolated MMA peptides, 62 are RBPs; 17 of which are hypomethylated in Δ*prmt7* (Hochberg and Benjamini test *q*<0.05.) (B) 51 proteins are differentially methylated between WT and Δ*prmt7*; 6 of which are both hypomethylated and hypermethylated at different residues. (C,D) Significantly enriched Molecular Function Gene Ontology (GO) Terms (upper panel) and selected amino acid residues (lower panel) in the whole MMA proteome (C) versus Δ*prmt7* hypomethylated proteins (D). For the global data, RNA binding was highlighted (upper panel), yet there is no clear consensus MMA site sequence (lower panel). RNA binding remains characteristic of Δ*prmt7* hypomethylated proteins, and an enrichment for the “RGG” motif is evident (lower panel). Dotted lines in upper panels depict Bonferroni p-value=0.05.

Molecular Function Gene Ontology (GO) Term analysis reveals that nucleic acid binding and RNA-binding functions are significantly enriched in the global MMA proteome and more notably in the Δ*prmt7* hypomethylated proteins (Figures 2C and D). Among 62 RNA-binding proteins (Supplementary Table S2), we found 24 differentially methylated in Δ*prmt7* cells, of which 17 are hypomethylated and represent putative substrates of PRMT7 activity (Supplementary Table S2). Analysis of the peptide sequences surrounding the methylated arginines using WebLogo 3 (33) demonstrate differential residue selection between the hypomethylated arginines versus total MMA proteome in Δ*prmt7* cells. The global MMA data do not present a significant motif consensus, with 10-20% probability for glycines to surround the methylarginine (Figure 2C). However, there is a strong selection evident for RG/RGG (or GR/GGR) motifs in the hypomethylated PRMT7 substrate peptides (Figure 2D), revealing a candidate target motif for the PRMT7 enzyme. Of interest, MMA sites in isolated hypomethylated RBPs reside outside conserved Interpro/Pfam domains (Figure 3), suggesting RBP-RNA interactions might not be directly obstructed.

**Figure 3.**
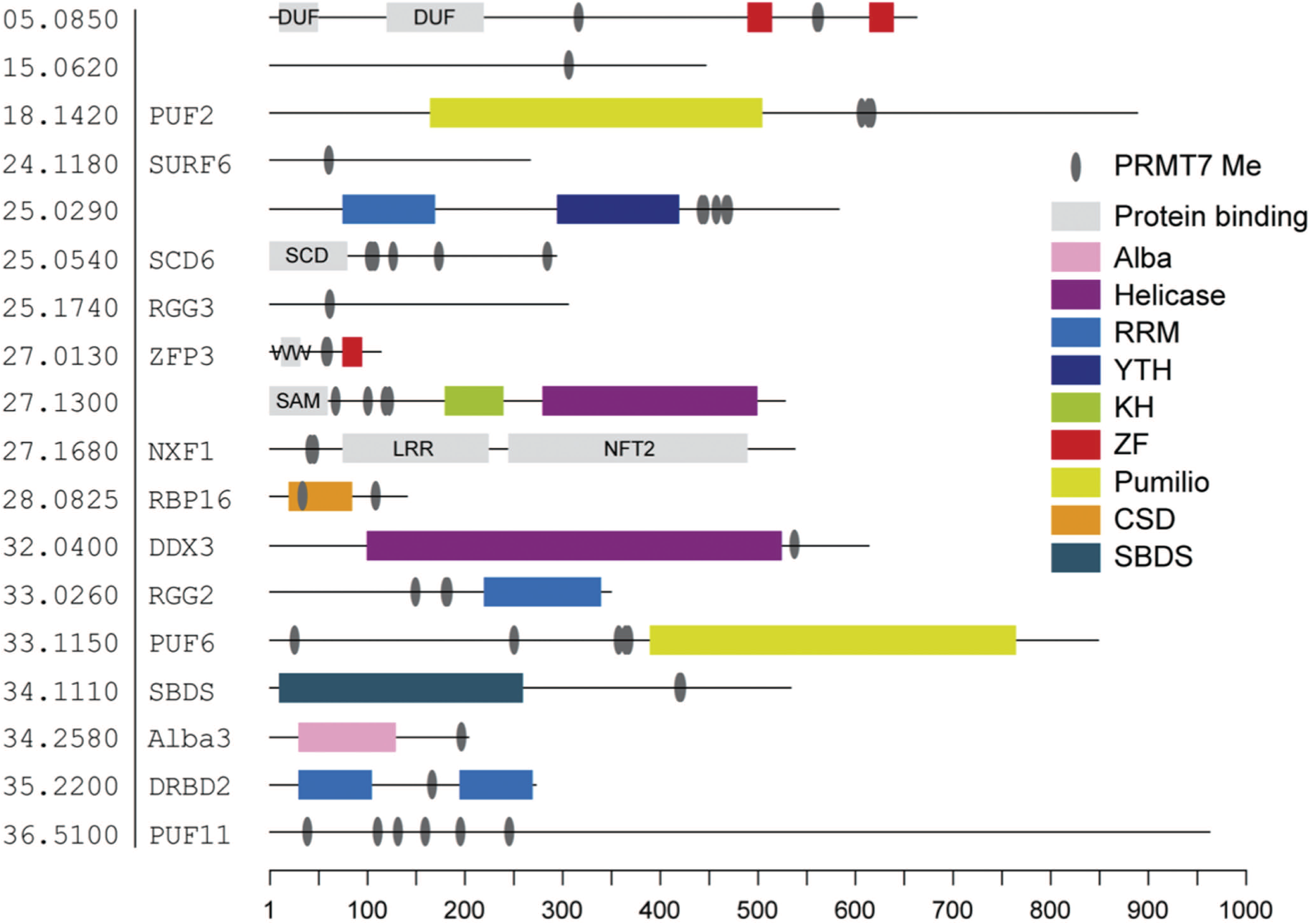
Monomethyl arginines are found outside RNA-binding domains. Bioinformatic analysis of 18 PRMT7 targets showing sites where MMA was detected and the predicted locations of different classes of RNA- or protein-binding domains. RNA-binding domains are colour coded (see legend) while protein-binding domains are annotated and shown in grey. Regions lacking an identifiable domain or with predicted disorder are shown by a solid line. Domain annotations: DUF, domain of unknown function; KH, K homology; LRR, Leucine-rich repeat; LSm, like Smith antigen; NFT2, nuclear transport factor 2; SAM, sterile alpha motif; SBDS, Shwachman-Bodian-Diamond syndrome protein; YTH, YT521-B homology. Positions of domain boundaries and methylated arginines are relative to the scale bar. Entries ordered by TriTrypDB gene identifier. Protein names provided where assigned.

### Validating RBP substrates of PRMT7 methylation

From the list of proteins hypomethylated in Δ*prmt7*, we selected Alba3 (Alba20; (Ferreira, 2014 #628)(34,35)) and RBP16 for further investigation (Figure 2A). Alba3 is a RPP25-like protein (ribonuclease P subunit p25-like) involved in the stabilization of the *δ-amastin* virulence factor mRNA upregulated in amastigotes (35). RBP16 is a Y-box binding protein (YBX2-like) important for trypanosomatid mitochondrial mRNA editing and stabilization (36). Our previous investigation identified these two RBPs as candidate targets of PRMT7 methylation (11). To determine whether these RBPs are directly targeted by PRMT7 methylation, purified recombinant 6xHis-tagged PRMT7 (His-PRMT7) and putative substrates His-Alba3 and His-RBP16 were tested for activity *in vitro*. We first confirmed His-PRMT7 activity is detectable *in vitro* using a commercially available human Histone H4, a canonical target of mammalian PRMTs (Figure 4A). In the absence of substrate, it appeared His-PRMT7 automethylates (Figure 4A), however we found this to be artifactual and disappeared when replaced with an alternative His sequence (data not shown).

**Figure 4.**
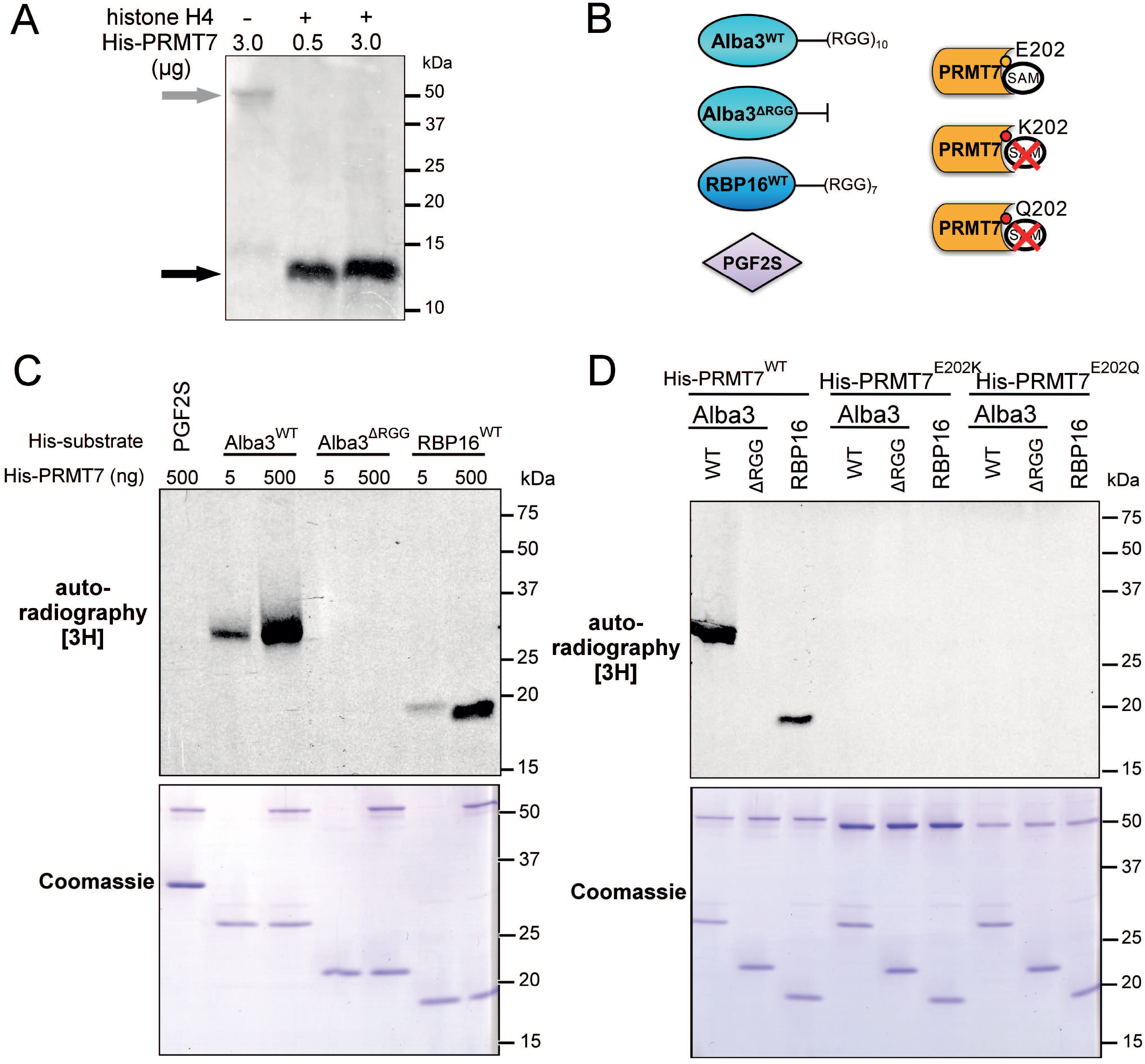
*In vitro* methylation of Alba3 and RBP16 by PRMT7. (A) Recombinant His-PRMT7 was tested for methyltransferase activity *in vitro* in the presence of radioactive methyl donor S-adenosyl methionine (SAM) and human histone H4 as a canonical RGG-containing PRMT substrate (black arrow). Artifactual automethylation of HIS-PRMT7 is detected in the absence of substrate due to HIS tag sequence (grey arrow). (B) Purified His-tagged PRMT7 putative substrates Alba3^WT^, RBP16^WT^ and RGG-deficient Alba3^ΔRGG^ were tested, as well as LbrPGF2S^WT^ (negative control). PRMT7 E202 residue in the double E loop motif was mutated to K or Q to generate catalytically-inactive mutants. (C) RNA-binding proteins Alba3^WT^ and RBP16 are arginine methylated by PRMT7 *in vitro*, while Alba3^ΔRGG^ and LbrPGF2S^WT^ are not. (D) *In vitro* methylation of target RBPs by PRMT7 is disrupted by mutating residue E202 (PRMT7^E202K^ and PRMT7^E202Q^).

Next, we examined His-PRMT7 methylation of recombinant His-RBP16 and His-Alba3, as well as a mutant protein devoid of the RGG-rich C-terminal tail (His-Alba3^ΔRGG^, Figure 4B). His-PRMT7 methylates His-Alba^WT^ and His-RBP16^WT^ *in vitro* in a concentration-dependent manner but not His-Alba3^ΔRGG^ (Figure 4C). This indicates that recombinant PRMT7 is sufficient to monomethylate both RBP targets directly and that PRMT7-catalyzed methylation requires the C-terminal RGG motifs of Alba3, despite 18 additional non-RGG arginine residues still present in His-Alba3^ΔRGG^. As a negative control, we examined the potential for PRMT7 to modify a *Leishmania* protein containing 12 arginine residues but lacking a defined RGG motif, His-PGF2S (LbrM.31.2410, Figure 4B). PGF2S is not methylated by PRMT7 *in vitro* (Figure 4C).

Building upon these observations, we generated two catalytically-inactive PRMT7 mutants to confirm conservation of the catalytic site in *Leishmania* and the specificity of the observed methylation signal. In other eukaryotes, key glutamate (E) residues in the double E loop of the SAM-binding domain have been replaced to generate inactive PRMT mutants (37,38). Analysis of the double E loop sequence of PRMTs in *L. major* showed conservation of residues E202 and E211 (Supplementary Figure S1). The first glutamate residue in this domain (E202) was mutated to either a glutamine or lysine accordingly. In contrast to His-PRMT7^WT^, recombinant His-PRMT7^E202Q^ and His-PRMT7^E202K^ do not methylate His-Alba3 or His-RBP16 (Figure 4D) *in vitro*. These results confirm the catalytic domain and the double E loop are essential for PRMT7-dependent monomethylation of RBP targets *in vitro*.

### Loss of PRMT7 impacts RBP expression and function *in vivo*

To investigate a possible role of MMA arginine in mRNA metabolism in *Leishmania* parasites, the N-termini of target RBPs were endogenously HA-tagged, leaving *3’UTRs* intact as transcript stability and translational control are overwhelmingly *3’UTR*-driven (39). Wild-type and Δ*prmt7* parasites were engineered to express HA-tagged Alba3, RBP16, Torus and DDX3 RBPs (Supplementary Figure S2). Torus is a CCCH-type zinc finger protein with a conserved Torus domain that, like DDX3 (HEL67; (Padmanabhan, 2016 #780)) co-immunoprecipitates with PRMT7 (11). Examination of candidate target HA-RBP levels in WT and Δ*prmt7* promastigotes at log (proliferative) and stationary (non-proliferative) growth stages show that three RBPs are differentially expressed (Figure 5A). Both HA-Torus and HA-RBP16 have reduced expression in Δ*prmt7* parasites, the former at both log and stationary phase, the latter only in stationary phase promastigotes. In contrast, HA-DDX3 is slightly upregulated in Δ*prmt7* at stationary phase with potential degradative products evident. The detection of a smaller HA-DDX3 band of ∼30 kDa at the stationary phase suggests enzymatic cleavage, a possible sign of protein instability and tightly coordinated control of expression. The observation that HA-RBP16 protein levels are reduced in Δ*prmt7* relative to wildtype human-infective stationary promastigote cells (Figure 5A) is particularly interesting as PRMT7 expression is tightly downregulated prior to this lifecycle stage (11). The stage-specific eradication of RBP16 levels in the Δ*prmt7* mutant cells was therefore examined in two different but complementary experimental contexts.

To determine whether reduced HA-RBP16 levels in Δ*prmt7* cells are due to expedited protein decay rates or reduced translation of *rbp16* transcript, the protein stability of both RBP16 and Alba3 was assessed over a time course in translationally-inhibited WT and Δ*prmt7* cells (Figure 5B and Supplementary Figures S3B, S3C). Of note, HA-RBP16 degradation is significantly faster in the absence of PRMT7 levels than in WT cells with the half-life of HA-RBP16 reduced specifically in stationary phase Δ*prmt7* cells relative to WT background or log phase (Figure 5B). In contrast, stability of HA-Alba3 protein was unchanged between samples (Supplementary Figures S3B, S3C). Densitometry analysis revealed HA-RBP16 protein half-life is >6 h in WT and 0.5 h in Δ*prmt7* cells at stationary phase (Figure 5B and Supplementary Figure S3C). This suggests PRMT7-dependent methylation can post-translationally control select target protein degradation rates in a stage-specific manner.

**Figure 5.**
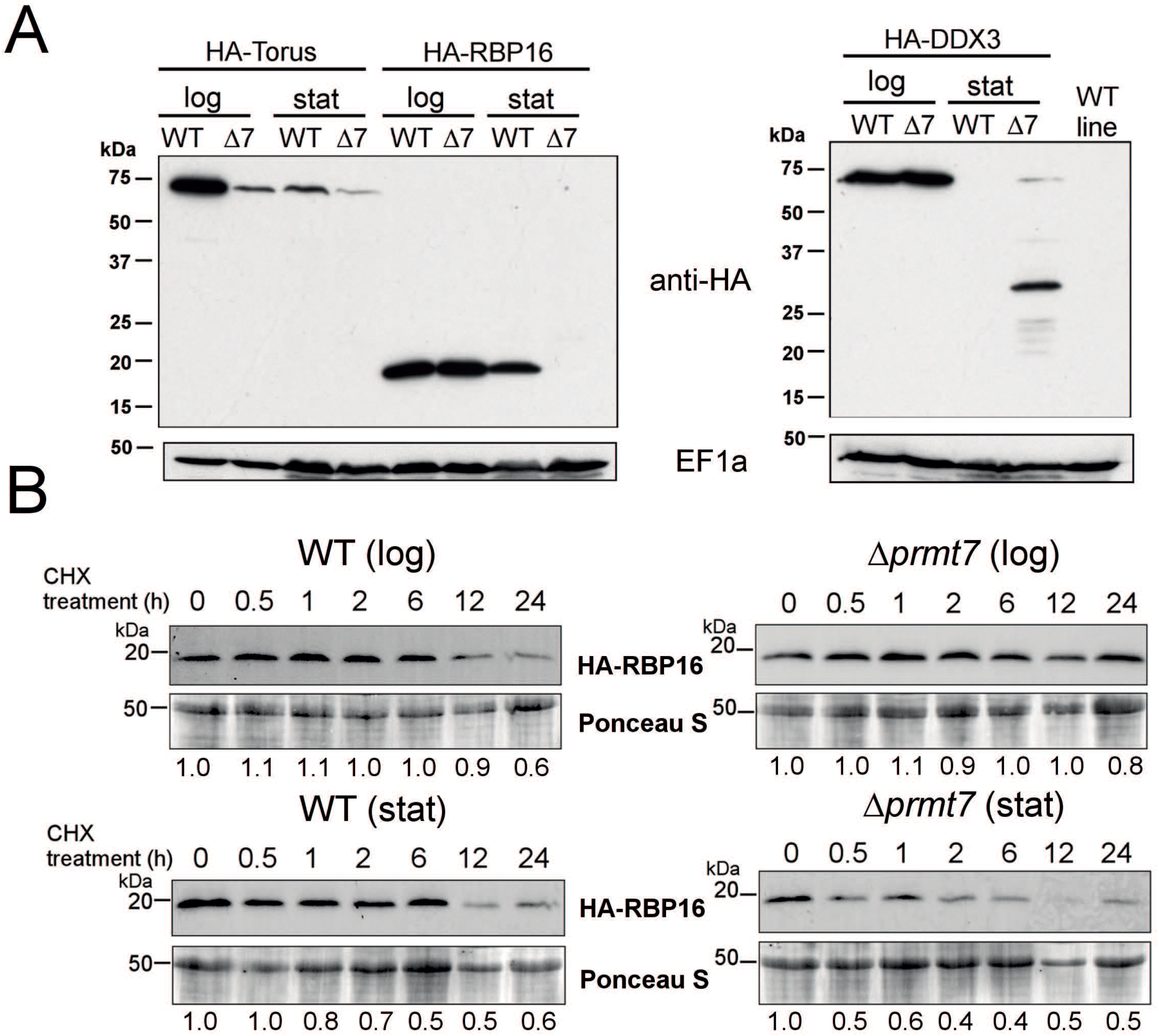
PRMT7 knockout affects target RBP expression *in vivo*. (A) Expression levels of 3 endogenously-tagged RBPs, Torus, RBP16 and DDX3, are altered in Δ*prmt7* (Δ7) parasites. Lanes are labeled for logarithmic (log) versus stationary (stat) phase promastigotes. EF1a levels are shown as a loading control. (B) HA-RBP16 protein stability was analyzed in log and stationary promastigotes after cycloheximide (CHX) addition. Ponceau staining is shown as loading control. HA-RBP16/Ponceau ratio values are shown below each lane.

To determine whether the half-life of HA-RBP16 was impacted by subcellular localization, N-terminally tagged HA-RBP16 was examined using immunofluorescence. Similar to immunoblot results, the HA-RBP16 protein signal specifically decreases in Δ*prmt7* promastigotes at stationary phase (Supplementary Figure S4C). Unlike the close RBP16 orthologue in *T. brucei*, TbRBP16 (36,40) N-terminally tagged HA-RBP16 localises in the cytoplasm and does not traffick to the mitochondria (Supplementary Figure S4C). Cytoplasmic localization of HA-Alba3 was unaltered between promastigotes samples (Supplementary Figure S4C). We therefore examined the subcellular localisation and relative stability of endogenous LmjRBP16 using anti-TbRBP16 (kind gift of L.Read) and found that while the HA-RBP16 protein is cytoplasmic, untagged endogenous LmjRBP16 is indeed mitochondrial (Supplementary Figure S4C). We tested the subcellular localisation of C-terminally-tagged endogenous RBP16-HA and found it is also mitochondrial (data not shown). This suggests N-terminal HA tagging of LmjRBP16 is sufficient to disrupt proper mitochondrial localisation and correlates with the absence of bound cytoplasmic RNA targets (data not shown). Of note, neither the mitochondrial-localised endogenous LmjRBP16 or endogenously tagged RBP16-HA are destabilized in the stationary cell stage in the absence of PRMT7 levels. As *rbp16* transcript is nuclear-derived and cytoplasmically translated, there is biological relevance to the relative stability of this potential metabolic regulator in the cytoplasm. This provides two useful mechanistic insights, firstly that PRMT7 methylation can shield a cytoplasmic RBP from protein degradation in an *in vivo* context and secondly that the protein degradative machinery context differs significantly between both cytoplasmic and mitochondrial cellular compartments and the distinct lifecycle stages.

### PRMT7-mediated methylation modulates post-transcriptional gene control

Similarly, endogenous levels and subcellular localisation of HA-Alba3 remain unaltered by removal of PRMT7 expression (Supplementary Figure S4D). We generated hypomethylated Alba3^WGG^ heterozygous mutants with RGG methyl sites replaced by WGG in the protein C-terminus to evaluate possible effects of an increased hydrophobicity in this region. In contrast to other RBPs investigated, there were no changes in either endogenously tagged HA-Alba3^WT^ or HA-Alba3^WGG^ protein levels between transfectants at the two promastigote stages analyzed (Figure 6A). Therefore, we investigated the potential influence of arginine methylation on Alba3 mRNA-binding function. Immunoprecipitation of HA-Alba3 and probing with anti-MMA revealed that monomethylation is only detected in WT cells and absent in HA-Alba3^WGG^ mutant protein expressed in both WT and Δ*prmt7* cell lines (Figure 6B), validating the C-terminal RGG motifs are necessary for *in vivo* Alba3 methylation by PRMT7.

**Figure 6.**
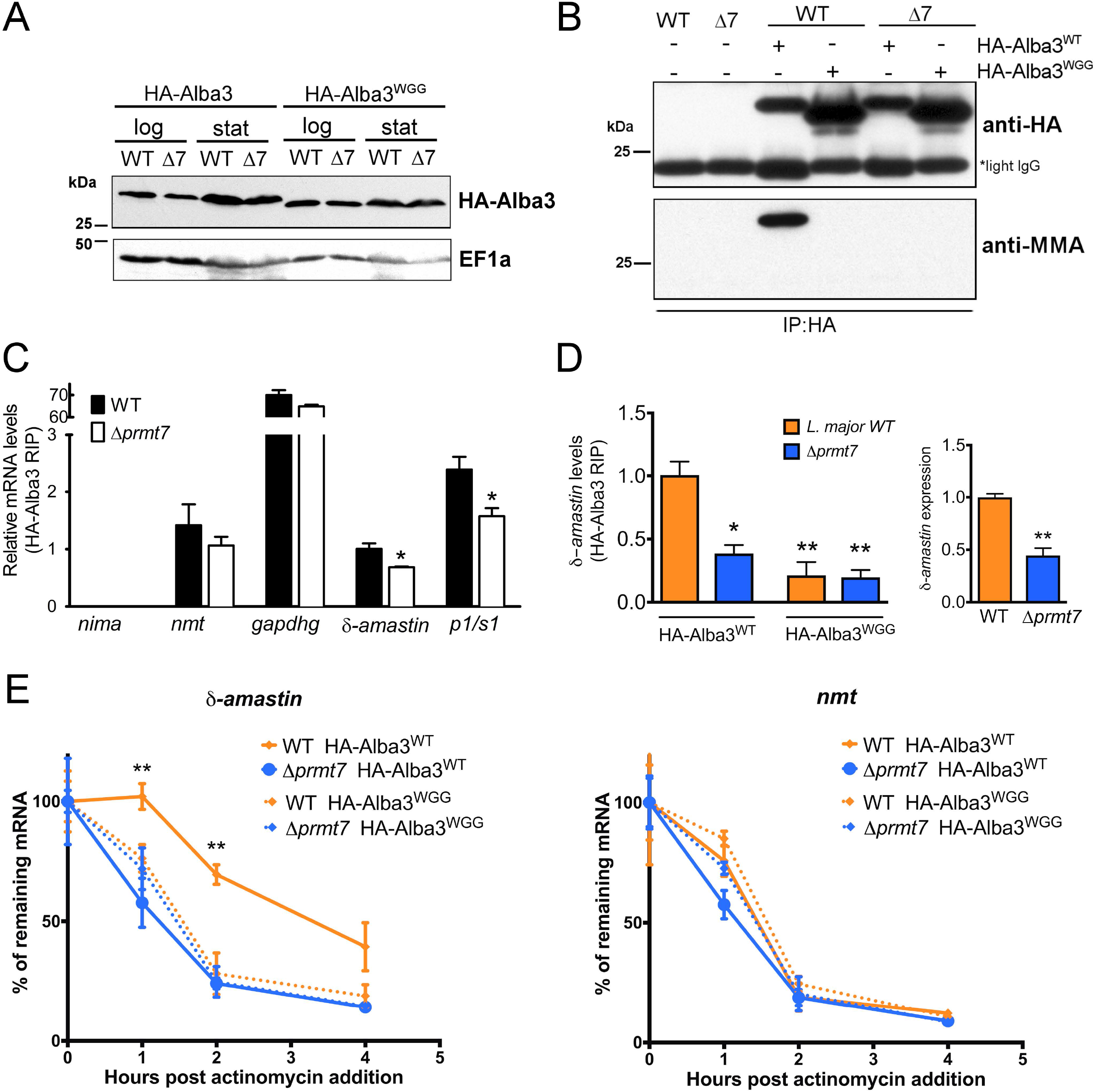
Arginine methylation of Alba3 contributes to mRNA binding and *δ-amastin* stabilization. (A) Endogenously tagged Alba3^WT^ and RGG->WGG hypomethylated mutant (Alba3^WGG^) showed no differential expression levels between WT and Δ*prmt7* cells. The migration of HA-Alba3^WGG^ mutant protein is faster on SDS-PAGE, independent of PRMT7 levels. (B) Immunoprecipitation (IP) of HA-Alba3 shows loss of arginine monomethylation in Δ*prmt7* parasites and when the C-terminal RGG motifs are mutated into WGG. (C) Alba3 complexes were purified by RNA co-immunoprecipitation (1) from *in vivo* UV-crosslinked cell lysates. Eluted RNA are analyzed by qRT-PCR and suggest that *δ-amastin* and *p1/s1* nuclease binding to Alba3 decreases in the absence of PRMT7. Transcript levels are relative to 18S rRNA. *p<0.05, Bonferroni two-way ANOVA test. (D) qRT-PCR analysis of *δ-amastin* mRNA association to HA-Alba3 and HA-Alba3^WGG^ mutant. Quantification of HA-Alba3 RIP eluted transcripts (left panel) and total input RNA (right panel) is shown. Constitutive *nmt* control mRNA levels are shown in Supplementary Figure S3A. *p<0.05, **p<0.01, Tukey one-way ANOVA test (left panel) and unpaired two-tailed t-test (right panel). (E) Stability of *δ-amastin* (left panel) and control *nmt* (right panel) transcripts were evaluated in wild-type (WT) and Δ*prmt7* after sinefungin and actinomycin treatment. RNA samples from cells expressing wild-type (Alba3^WT^) or hypomethylated (Alba3^WGG^) Alba3 were used in qRT-PCR. *δ-amastin* mRNA half-life was >3.5h in WT and ∼1h in Δ*prmt7* cells expressing HA-Alba3^WT^. Transcript levels were normalized against 18S rRNA and are plotted as mean ± standard error of two biological replicates. \*\**p*<0.01, Bonferroni two-way ANOVA test.

Next, we addressed the RNA-binding properties of hypomethylated Alba3 protein. Previously, *Li*Alba3 has been shown to bind to *δ-amastin* (LinJ.34.1010) and *p1/s1 nuclease* (LinJ.30.1520) transcripts in *Leishmania infantum* cells (28). Therefore, we tested whether Alba3 RNA-binding is conserved and impacted by PRMT7-dependent methylation. Promastigotes cells were UV-crosslinked *in vivo* followed by anti-HA RNA co-immunoprecipitation to isolate HA-Alba3 mRNP complexes and specific transcript targets were identified via RNA MiSeq and quantified by qRT-PCR. Remarkably, endogenous HA-Alba3 showed differential RNA-binding affinity in the absence of PRMT7-dependent methylation. The *in vivo* interaction of Alba3 with stage-regulated mRNAs *δ-amastin* (LmjF.34.0500) and *p1/s1 nuclease* (LmjF.30.1510) is dependent on methylation by PRMT7 (Figure 6C). In contrast, Alba3 binding to the constitutively expressed *nmt* (N-myristoyltransferase) and *gapdh* mRNA targets is not altered by PRMT7 knockout. While total *δ-amastin* expression levels are reduced in Δ*prmt7, p1/s1 nuclease* expression is unchanged, suggesting divergent regulatory mechanisms between the two transcripts.

To further analyze Alba3 binding to *δ-amastin*, HA-Alba3^WT^, HA-Alba3^WGG^, HA-Alba3^R192K^ and HA-Alba3^KGG^ mRNPs were purified from WT and Δ*prmt7* samples and quantified by qRT-PCR. Interaction between Alba3 and *δ-amastin* mRNA was again inhibited in Δ*prmt7* cells *in vivo* (Figures 6C, 6D and 7C). This interaction was disrupted in HA-Alba3^WGG^ and HA-Alba3^KGG^ (Figures 6D and 7C), corroborating the role of C-terminal tail methylation upon Alba3 RNA-binding capacity. As a control, *nmt* mRNA associated with Alba3 at similar levels in WT and Δ*prmt7* cells and *nmt* expression was unchanged (Figures 6E and 7C, Supplementary Figure S3A). Interestingly, mutation of a verified PRMT7 arginine target amino acid to a lysine (R192K) did not disrupt Alba3 association with *δ-amastin* or *nmt*, but was sufficient to disrupt Alba3 association with *p1s1* target transcript (Figure 7C). Indeed, the trypsin digest of proteins prior to isolation of monomethylated peptides likely disrupted the proper identification of multiple RGG motif-containing peptides from the Alba3 C-terminus. This suggests Alba3 association with *δ-amastin* mRNA requires PRMT7-dependent monomethylation of C-terminal RGG motifs in addition to R192.

**Figure 7.**
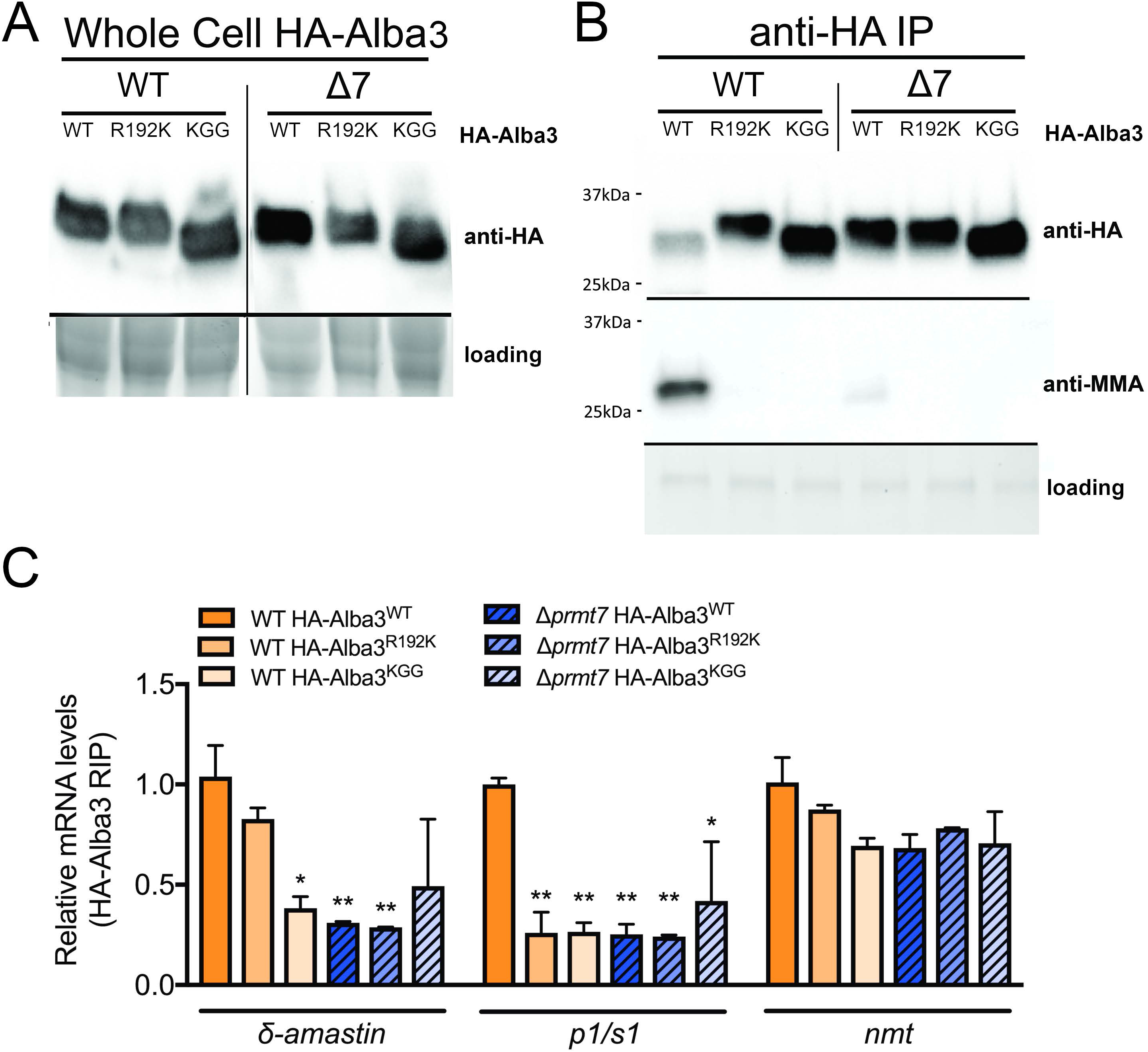
Arginine methylation of Alba3 contributes to binding of *δ-amastin* and *p1s1* mRNAs. (A) Endogenously tagged Alba3^WT^, RGG->KGG single point mutant (Alba3^R192K^) and hypomethylated mutant (Alba3^KGG^) showed no differential expression levels between WT and Δ*prmt7* cells. The migration of HA-Alba3^KGG^ mutant protein is faster on SDS-PAGE, independent of PRMT7 levels. (B) Immunoprecipitation (IP) of HA-Alba3 shows arginine monomethylation is ablated in the RGG->KGG single point mutant (Alba3^R192K^), when all C-terminal RGG motifs are mutated into KGG (HA-Alba3^KGG^) and in Δ*prmt7* parasites. (C) Alba3 complexes were purified by RNA co-immunoprecipitation from *in vivo* UV-crosslinked cell lysates. Eluted RNA were analyzed by qRT-PCR and confirm that Alba3 binding to *δ-amastin* and *p1/s1* nuclease decreases in the absence of PRMT7, but Alba3:*nmt* association remains constant independent of R mutation or PRMT7 levels. Interestingly, mutation of R192K disrupts Alba3:*p1s1* association, but only mildly impacts Alba3:*δ-amastin* binding. Transcript levels are relative to 18S rRNA. *p<0.05, **p<0.01, Bonferroni twoway ANOVA test.

Of interest, quantification of total *δ-amastin* levels revealed reduced expression in Δ*prmt7* cells, suggesting regulation via MMA-Alba3 binding (Supplementary Figure S3A and Figure 6E). It has been reported that Alba3 participates in the stabilization of *δ-amastin* mRNAs in *L. infantum* (34). We thus investigated if PRMT7-mediated methylation could affect *δ-amastin* transcript stability.

Samples from cells treated with sinefungin and actinomycin, to block RNA *trans*-splicing and transcription, respectively, revealed that *δ-amastin* is only stabilized when Alba3 is monomethylated (Figures 6E and 7C). Either PRMT7 knockout or expression of hypomethylated Alba3 (HA-Alba3^WGG^) resulted in a significant decrease of *δ-amastin* mRNA half-life (from > 3.5 h in WT, to 1 h in Δ*prmt7*). In contrast, no significant changes were observed in the stability of the constitutively expressed *nmt* target transcript between samples (Figures 6E and 7C). Importantly, this indicates PRMT7-dependent MMA methylation of Alba3 impacts RNA binding in a target-specific manner. Collectively our data demonstrate methylation by PRMT7 can regulate the relative protein stability and selective RNA affinity of distinct RBP targets.

## DISCUSSION

To date, a limited number of arginine methylated proteins have been identified in *Leishmania* spp. (11,41). In this study, we report that 3% (∼247) of the entire *L. major* predicted proteome carries at least one monomethylated arginine (42). This number of MMA-targeted proteins is relatively large considering arginines can also be dimethylated and since other post-translational modifications (PTMs) have been detected in *Leishmania* spp., including ∼600 phosphorylated proteins (7%) (43). We demonstrate that PRMT7-dependent methylation strongly impacts the MMA proteome of *L.major* parasites thus regulating RBP expression and/or function (Figure 8). Notably, PRMT7 levels regulate the expression levels of three isolated RBP targets in at least one developmental stage, suggesting that arginine methylation can affect protein turnover in *Leishmania* spp.

**Figure 8.**
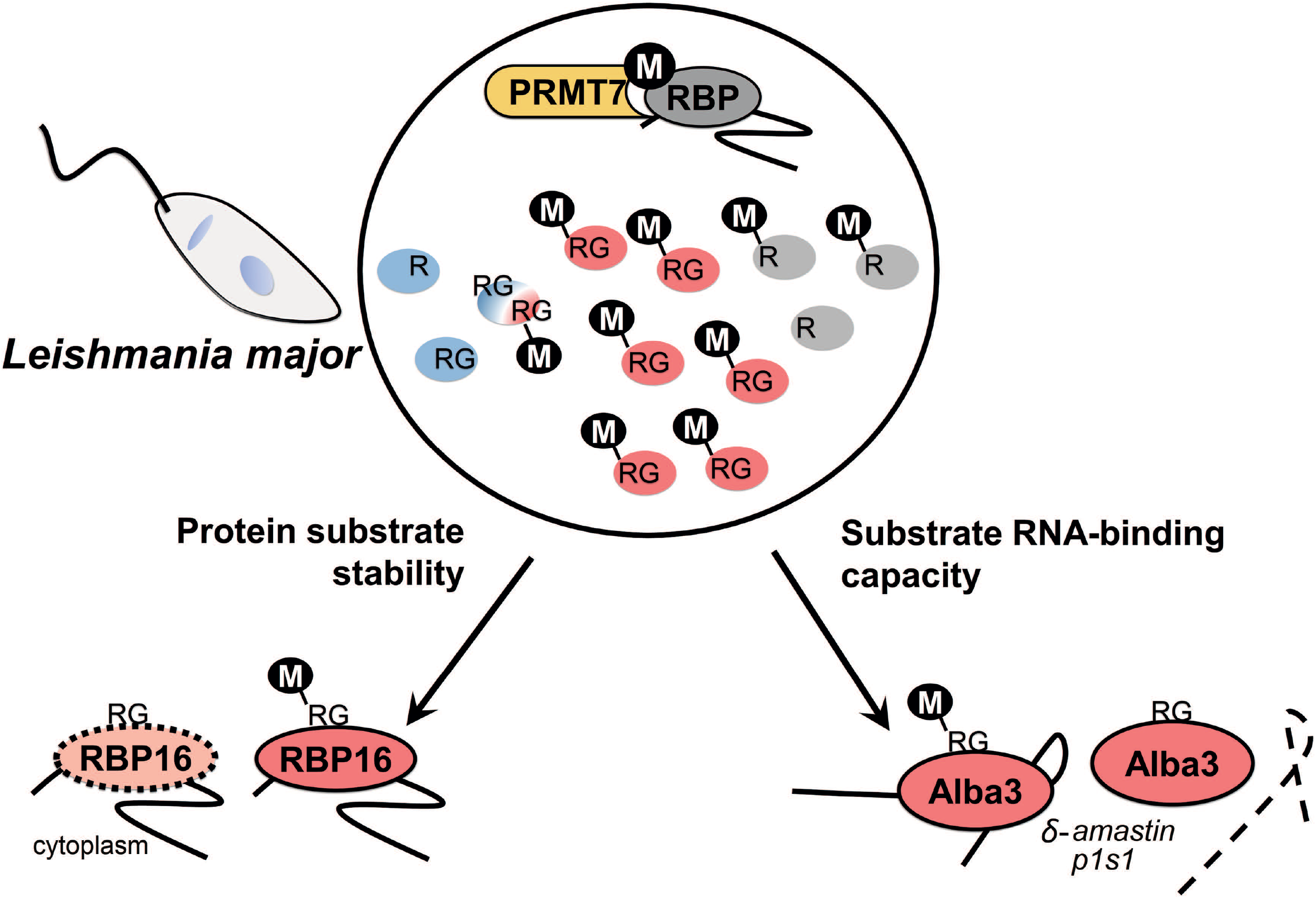
PRMT7 regulates RNA-binding protein function in *Leishmania*. Overview diagram describing the putative role of PRMT7-dependent methylation based on the effects of *L. major* PRMT7 knockout upon the arginine methylation of selected RBPs. Red circles represent RBPs hypomethylated in Δ*prmt7*, while blue are hypermethylated. PRMT7 contributes to the protein stability of cytoplasmic RBP16 in stationary promastigotes and to Alba3 RNA-binding affinity *to δ-amastin* and *p1/s1* nuclease mRNAs. Hypomethylation of Alba3 consequently results in decreased *δ-amastin* half-life.

Overall, experimental evidence indicates *Leishmania major* PRMT7 activity impacts protein target stability and function in a context-dependent manner, distinct from that of mammalian homologs. Despite not being an essential protein in *L. major*, PRMT7 levels regulate leishmaniasis disease pathology (11). Remarkably, epigenetic control by PRMT7-dependent monomethylation of the primary gene regulators in this system, RBPs, mediates virulence of the human-infective lifecycle stage at least two differentiation events downstream of PRMT7 expression. This undermines the traditional paradigm that influential *trans*-regulators are temporally synchronized with the cellular processes they promote and provides another layer of depth to the complexities of *Leishmania* genetics.

Interestingly, 17 proteins are hypermethylated in Δ*prmt7* parasites (Figures 2A and 2B). This may suggest modulation of alternative PRMT activity to functionally compensate for PRMT7 depletion. Six of the hypermethylated proteins also contain concurrent hypomethylated distal sites; indicative of a complex PRMT inter-regulatory system. Deletion of mammalian PRMT1, which catalyzes asymmetric dimethylarginine (ADMA), results in an increase of global MMA in mammals and *T.brucei* (44,45). Therefore, functional overlap between *Leishmania* PRMTs may involve a dynamic regulatory interplay as it does in *T.brucei* (46). A unique example of substrate competition by PRMTs in human cells is regulation of transcription elongation factor E2F-1-mediated apoptosis by arginine methylation (47). Here, PRMTs determine apoptotic outcomes by competing for the same substrate as SDMA methylation of E2F-1 by PRMT5 leads to apoptosis in DNA damaged cells, while ADMA methylation by PRMT1 favors proliferation. The detection of PRMT3 MMA peptides (LmjF.03.0600; Supplementary Table S1) further supports methyltransferase inter-regulation. While PRMT3 is catalytically inactive in *T. brucei* due to the absence of key residues in the SAM-binding domain (48), the two essential glutamate residues in the double-E loop are still present in *Leishmania*, suggesting a different regulatory role for *Lmj*PRMT3. This exciting distinction suggests PRMT3 may be a functional enzyme rather than a prozyme in *Leishmania*.

Arginine methylation sites vary considerably according to the PRMT enzyme or the organism investigated (3,38,46,49). RG and RXR sites represent the majority of mammalian PRMT target motifs but there are exceptions (1,38,46). The present methyl-SILAC analysis identified RG/RGG as the main target for PRMT7 methylation *in vivo*. Conserved protein domain analysis revealed that most of PRMT7-dependent methylation is not found directly within classical RNA-binding domains (Figure 3). Of interest, no histone MMA peptides were identified by our screen. This could be due to the cytoplasmic localisation of *Lmj*PRMT7, methodological limitations or a low overall abundance of MMA in *Leishmania* histone proteins.

Although arginine methylation has been mostly associated with changes in protein-protein interactions (50,51), there are previous examples of direct modulation of RNA-protein binding in mammalian cells (52–54). PRMTs have previously been shown to regulate the RNA affinity of target RBPs in other systems (55,56). In *T. brucei*, the RNA-binding proteins DRBD18 and PRMT1 have been shown to modulate the fate of mRNAs according to methylation state, impacting both protein and mRNA binding (45,50). Here we dissect the effect of MMA upon RNA affinity and isolate the specific arginines of Alba3 that are monomethylated by PRMT7; resulting in the discrete maintenance or loss of specific transcript targets. In line with the previous finding that *L. infantum* Alba3 stabilizes *δ-amastin* transcripts (34), we find that overall *δ-amastin* levels are significantly reduced in the Δ*prmt7* mutant *L. major* lines, most likely due to reduced association with the hypomethylated Alba3. Amastins are thought to function as membrane transporters and are largely regarded as important virulence factors in *Leishmania* spp.(57). Reduced levels of *δ-amastins* via gene knockdown limits pathogenesis of *Leishmania* infection in mice. Our results indicate arginine monomethylation of the Alba3 C-terminus increases protein affinity to *δ-amastin* and *p1s1* mRNAs, promoting transcript stabilization (Figures 6C, 7C and Supplementary Figure S3A). Notably, expression levels of Alba target transcript *p1/s1 nuclease* are not altered by PRMT7 levels, although arginine methylation of Alba3 is necessary for association. These results suggest Alba3 regulates mRNA fate in a highly bespoke, target-specific manner. There is a low correlation between protein expression and mRNA binding capacity of RBPs and mRNPs display context-dependent selection of components in *Leishmania* (9). Further investigation of the molecular requirements and functional impact of RBP modification is therefore crucial to understanding gene regulation not only in *Leishmania*, but all eukaryotes.

The impact upon RNA binding potential may be indirect. The CCCH zinc finger protein ZFP3 is known to associate with ZFP1 in *T.brucei* and this interaction is necessary for both RNA target specificity as well as polysomal association (58). Here, we identify ZFP3 as a target of PRMT7-dependent MMA in *Leishmania* (Figure 2A). It is possible a ZFP3:ZFP1 interaction could be impacted by the presence of methylarginine altering both the RBP spatial proximity and RNA target binding. *In silico* docking analyses suggest several residues in the Alba domain are buried inside the Alba3:Alba1 heterodimeric protein complex but the RGG motif in the C-terminal tail is exposed (35,59). Consistent with the absence of RGG motifs, the Alba1 protein was not detected in our methyl-SILAC analysis. It is not known whether methylation of the Alba3 RGG tail regulates the stress-induced localization of the Alba proteins (35). Proper characterization of MMA target RBP structural dynamics is necessary to determine the impact of methylation on mRNP complex assembly.

Unusually, PRMT7-dependent methylation promotes the stability of cytoplasmic RBP16 protein beyond the temporal expression of PRMT7 protein levels. The regulation of protein turnover observed represents a novel function for PRMT7-dependent methylation (Figures 5A, 5B). *T. brucei* RBP16 was the first Y-box protein to be found in mitochondria, where it associates with guide RNAs (gRNAs) and has a role in mitochondrial mRNA editing (27,36). The N-terminal cold-shock domain and RGG-rich C-terminus of TbRBP16 both contribute to RNA-binding capacity (60). Arginine residues in the *T. brucei* RBP16 C-terminus are targeted by PRMT1 methylation, impacting RNA editing complexes (36). We find that RBP16 in *Leishmania* is also mitochondrial and this localization is not affected by the absence of PRMT7 (Supplementary Figure S4B). However, the N-terminal tagging of endogenous RBP16 protein renders it cytoplasmic, where the RBP half-life is specifically reduced in Δ*prmt7* knockout cells in a human-infective stage (Supplementary Figure S4B). This reduced stability suggests protein degradation can be inhibited in wild-type cells via PRMT7-dependent methylation, which is particularly interesting at a lifecycle stage in which PRMT7 is not expressed. This lends insight into the distinct microenvironments influencing protein degradation in cytoplasm versus mitochondria which can determine RBP function and stability.

Potential mechanisms include inhibition of protein ubiquitination and proteasome degradation targeting. Interdependent ubiquitination and methylation has been detected previously in other systems (61,62). Cytoplasmic protein turnover rate is largely regulated by the ubiquitin-proteasome system and polyubiquitination is the classical marker for this pathway (63). In mammalian innate immune response, TNF receptor-associated 6 (TRAF6) ubiquitin ligase activity is inhibited by PRMT1 methylation (62). Activation of Toll-like receptor response leads to hypomethylated TRAF6 via demethylation by JMJD6 and downregulation of PRMT1 expression. This triggers TRAF6 ubiquitin ligase activity and NF-kB response. In *L. major*, a putative JMJD6 demethylase ortholog gene may enable dynamic arginine methylation in protozoa (Tritrypdb.org). To fully understand the function of RBP16 in these parasites we must examine other PTMs controling RBP16 function.

The Lsm domain-containing SCD6 is another protein with simultaneous identification of hypo- and hypermethylated sites in the data presented here. This RBP is involved in processing body (P-body) granule formation in *T. brucei*; specifically responsible for coordinating granule assembly (64). Sequential deletion of the TbSCD6 RGG motifs, orthologous to targets of PRMT7 methylation identified by our *Leishmania* screen, led to a proportional decrease in RNA granule numbers in *T.brucei*. This suggests arginine methylation may play a crucial role in controlling the fate of all mRNPs in the cell; impacting both mRNA metabolism and sequestration. At least four other methylated RBPs have been shown to co-purify with RNA granules in *T. brucei*: DDX3, PABP1, DRBD4 and Alba3 (65).

In conclusion, we introduce *Leishmania* as a model organism to specifically study PRMT activity and functional impact of methylation upon non-histone targets. We explore methyl-SILAC as a productive, verified approach to investigate protozoa PRMT function and target isolation. Our data indicate that PRMTs have a central role in mRNA metabolism control by modulating the stability and function of post-transcriptional regulators in *Leishmania*. PRMT7 activity thus has a demonstrated role in gene expression control, both directly at the post-translational level and indirectly at the post-transcriptional level. Taken together, our data indicate the *Leishmania* PRMT7-directed regulatory pathway epigenetically controls parasite gene expression, and resultant pathology long after PRMT7 expression.

## SUPPLEMENTARY DATA

Supplementary Data are available at NAR online.

## ACKNOWLEDGEMENT

We thank Profs. Deborah Smith and Jeremy Mottram for thoughtful comments on this manuscript. We thank Drs. James Brannigan, Jaspreet Grewal, Vincent Geoghegan, Juliana Diniz, Sarah Forrester and Luis de Pablos for helpful discussions on the *Leishmania* SILAC labeling and methyl-SILAC data analysis. Proteomic and RNA identification and analyses were conducted in the MAP and Genomics and Bioinformatics Labs, respectively, in the Bioscience Technology Facility at the University of York (https://www.york.ac.uk/biology/technology-facility/).

