## Supplemental for "Post-translational epigenetics: PRMT7 regulates RNA-binding capacity and protein stability to control *Leishmania* parasite virulence"

**PRMT7 methylation controls *trans*-regulator function in *Leishmania* protozoa**

#### Supplemental Information

##### Supplemental Figure Legends

**Supplemental Fig S1.** Multiple sequence alignment of all the annotated PRMTs in *L. major* (*Lmj*PRMTs) and *Homo sapiens* (*Hs*PRMTs). The double E loop is part of the catalytic domain of the PRMTs and is important for S-adenosyl-methionine (SAM) and protein substrate binding. The asterisk depicts the *Lmj*PRMT7 E202 residue mutated to generate catalytically inactive E202K and E202Q mutants *in vitro*.

**Supplemental Fig S2.** Endogenous FLAG-HA-tagging of *L. major* RBPs (**A**) Plasmid map of pFLAG_HA plasmid, synthesized by GenScript for this work, with Alba3 5'flank region (5'FLR) and Alba3$^{WGG}$ mutant sequence in the "RBP-CDS" region. *Sfi*I enzyme sites used (A,B,C,D) produce different overhangs allowing the ligation of four different synthesized DNA sequences at the same time in a specific order (Fulwiler *et al*. 2011). Homologous sequences required for gene replacement are the 5'FLR (500bp) and the protein coding sequence (RBP-CDS, variable). (**B**) Primers that bind to the 5'FLR region of the gene and 3'FLR immediately downstream of the RBP-CDS but absent in the plasmid sequence were used to verify correct replacement of at least one allele by the HA-tagged gene. Expected PCR band sizes are shown for each tagged gene. (**C**) Positive clones were analyzed by PCR amplification of the modified HA-RBP allele (upper bands) and the original unmodified RBP allele (lower bands). DNA marker used was Gene Ruler 1kb (Thermo Fisher).

**Supplemental Fig S3.** (**A**) qPCR showing relative concentrations of transcript targets immunoprecipitated with HA-Alba3 in the presence or absence of PRMT7 levels relative to endogenous transcript levels. Alba3 transcript target nmt levels remain unchanged in the presence or absence of PRMT7 levels or the mutation of Alba3 RGG motifs to WGG. (**B**) Alba3 protein levels and stability are unaltered in the presence or absence of PRMT7 levels in log- or stat- stage promastigote cells. (**C**) Cytoplasmic HA-RBP16 levels display a reduced half-life specifically in stat-stage human-infective promastigotes in the absence of PRMT7 expression.

**Supplemental Fig S4.** **(A)** Western blot examining endogenous levels of *Lmj*RBP16 using anti-*Tb*RBP16 (kind gift of L.Read). *Lmj*RBP16 protein levels are constant in the presence and absence of PRMT7 levels. (B-D) Immunofluorescence of Log and Stat stage promastigotes: DAPI (Blue), Mitotracker (Green), RBP (Red), line = 5$\mu$m. (**B**) Subcellular localisation of endogenous *Lmj*RBP16 using anti-*Tb*RBP16 shows mitochondrial localisation constant in promastigote lifecycle stages examined. (**C**) Subcellular localisation of endogenously-tagged HA-*Lmj*RBP16 using anti-HA shows cytoplasmic localisation that is destabilized specifically in the absence of PRMT7 levels in stationary (stat) stage promastigote cells. (**D**) Subcellular localisation of endogenously-tagged HA-*Lmj*Alba3 using anti-HA shows cytoplasmic localisation constant in promastigote lifecycle stages examined.

**Supplemental Table Legends**

**Supplemental Table S1.** Global monomethyl arginine peptides identified and quantified by heavy methyl SILAC analysis in WT and Δ*prmt7 Leishmania major*.

**Supplemental Table S2.** Methylpeptides from RNA-binding proteins (RBPs) that are differentially methylated between WT and Δ*prmt7 Leishmania major*. Proteins were considered RBPs if they present an RNA-binding domain or if they are orthologs of a validated *Trypanosoma brucei* RBP.

**Supplemental Table S3.** List of oligonucleotides used in this study for PCR or qRT-PCR.

Figure S1

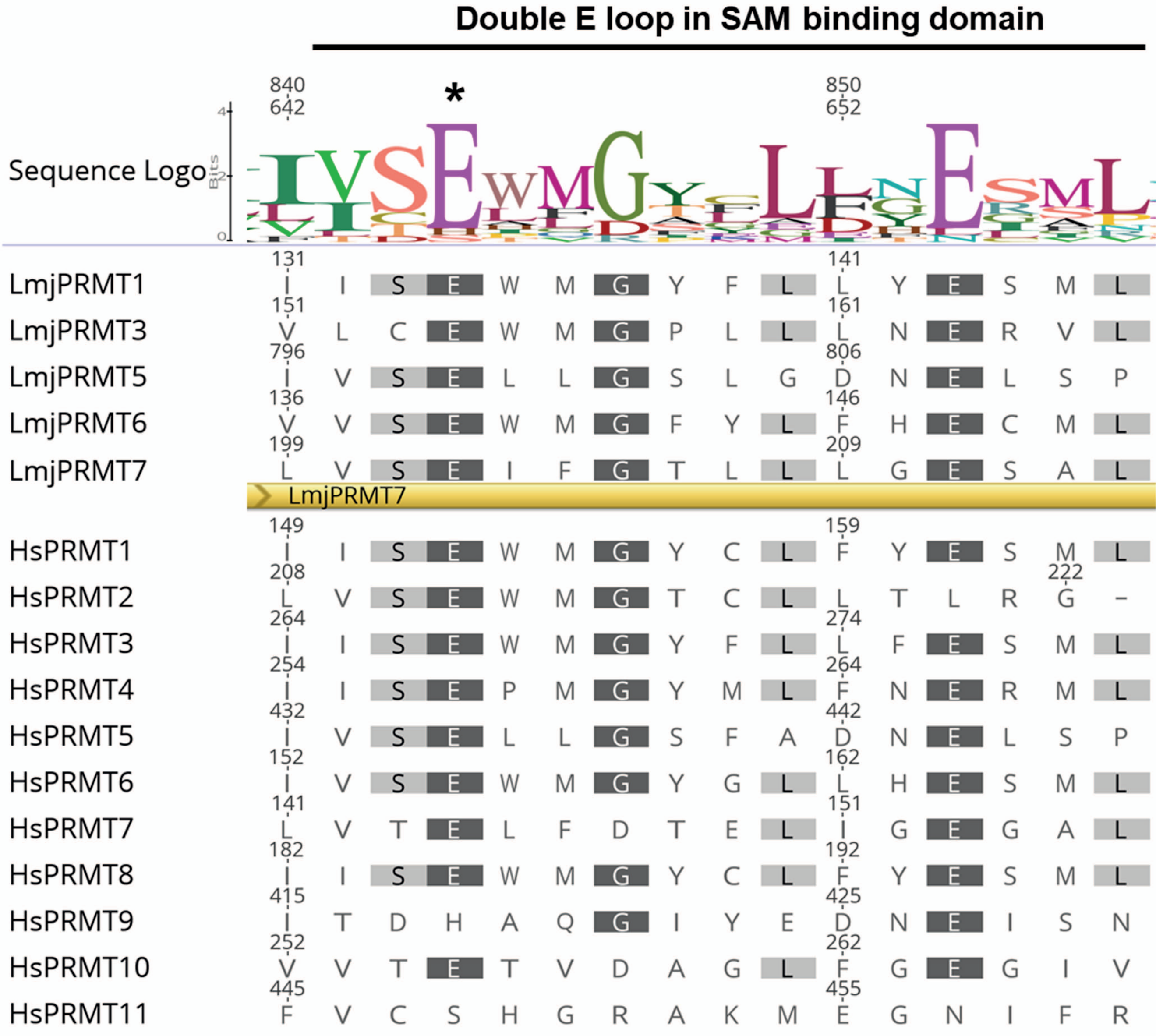
Double E loop in SAM binding domain
840 642
850 652
*
Sequence Logo
Bits
LmjPRMT1 131 I I S E W M G Y F L 141 L Y E S M L
LmjPRMT3 151 V L C E W M G P L L 161 L N E R V L
LmjPRMT5 796 I V S E L L G S L G 806 D N E L S P
LmjPRMT6 136 V V S E W M G F Y L 146 F H E C M L
LmjPRMT7 199 L V S E I F G T L L 209 L G E S A L
LmjPRMT7
HsPRMT1 149 I I S E W M G Y C L 159 F Y E S M L
HsPRMT2 208 L V S E W M G T C L L T L R 222 G –
HsPRMT3 264 I I S E W M G Y F L 274 L F E S M L
HsPRMT4 254 I I S E P M G Y M L 264 F N E R M L
HsPRMT5 432 I V S E L L G S F A 442 D N E L S P
HsPRMT6 152 I V S E W M G Y G L 162 L H E S M L
HsPRMT7 141 L V T E L F D T E L 151 I G E G A L
HsPRMT8 182 I I S E W M G Y C L 192 F Y E S M L
HsPRMT9 415 I T D H A Q G I Y E 425 D N E I S N
HsPRMT10 252 V V T E T V D A G L 262 F G E G I V
HsPRMT11 445 F V C S H G R A K M 455 E G N I F R

Figure S2

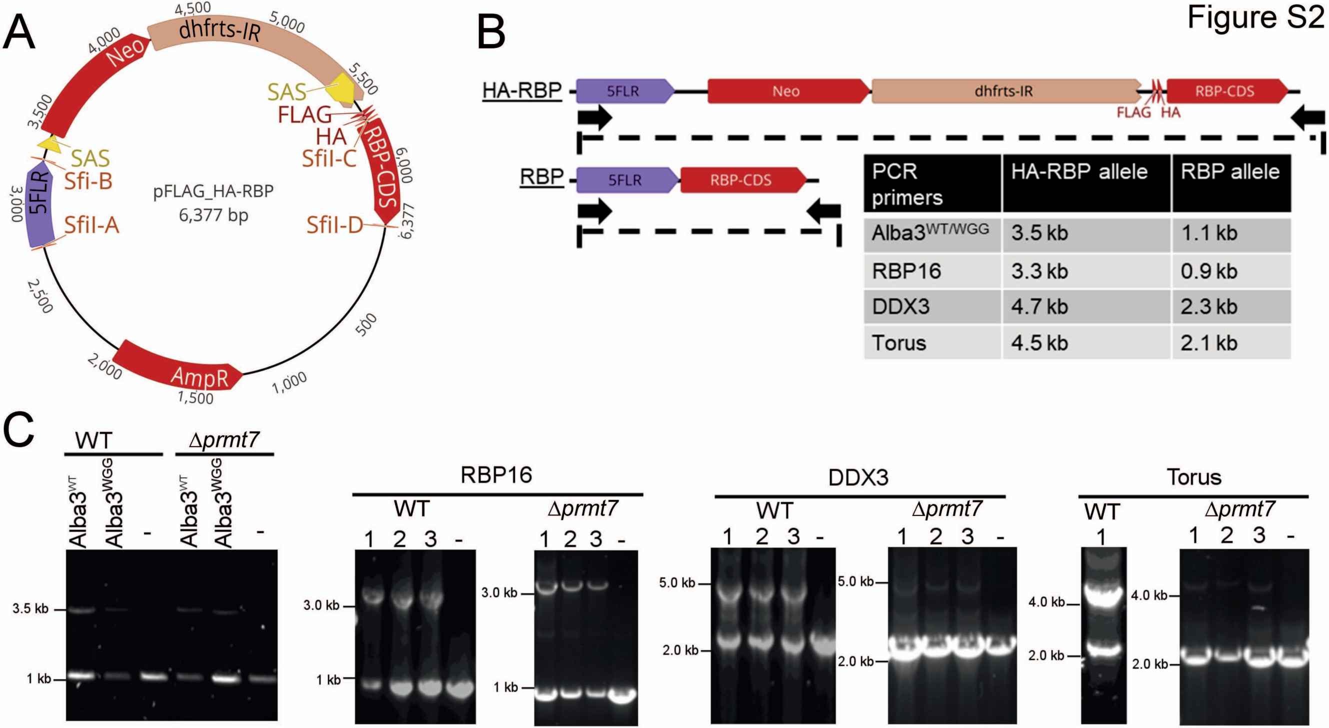

| PCR primers | HA-RBP allele | RBP allele |
|---|---|---|
| Alba3$^{WT/WGG}$ | 3.5 kb | 1.1 kb |
| RBP16 | 3.3 kb | 0.9 kb |
| DDX3 | 4.7 kb | 2.3 kb |
| Torus | 4.5 kb | 2.1 kb |

### Figure S3

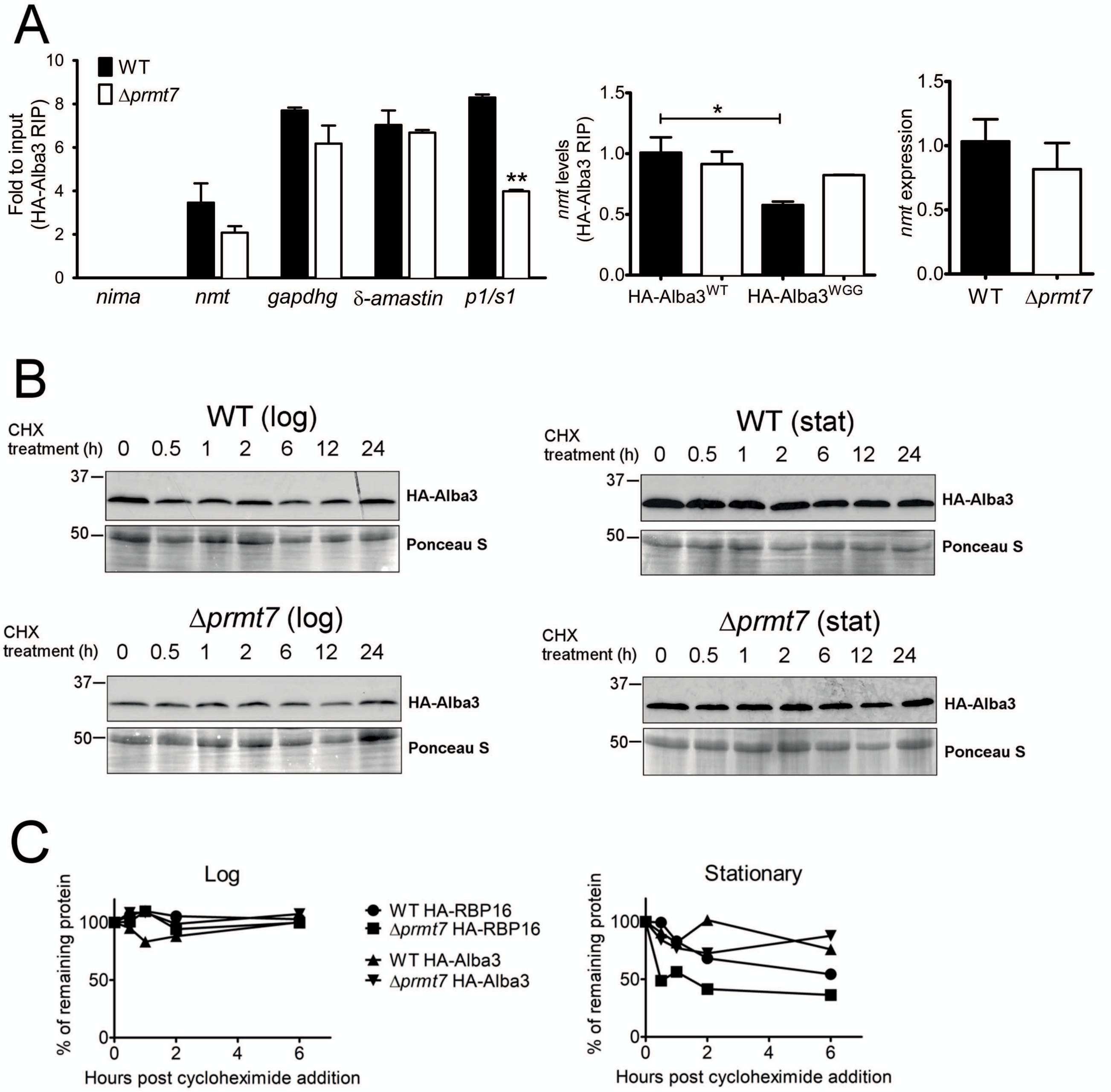

A
WT
Δprmt7
Fold to input (HA-Alba3 RIP)
nima
nmt
gapdhg
δ-amastin
p1/s1
**
nmt levels (HA-Alba3 RIP)
*
HA-Alba3^WT
HA-Alba3^WGG
nmt expression
WT
Δprmt7
B
WT (log)
WT (stat)
Δprmt7 (log)
Δprmt7 (stat)
CHX treatment (h)
0 0.5 1 2 6 12 24
HA-Alba3
Ponceau S
37
50
C
Log
Stationary
% of remaining protein
Hours post cycloheximide addition
WT HA-RBP16
Δprmt7 HA-RBP16
WT HA-Alba3
Δprmt7 HA-Alba3

### Figure S4

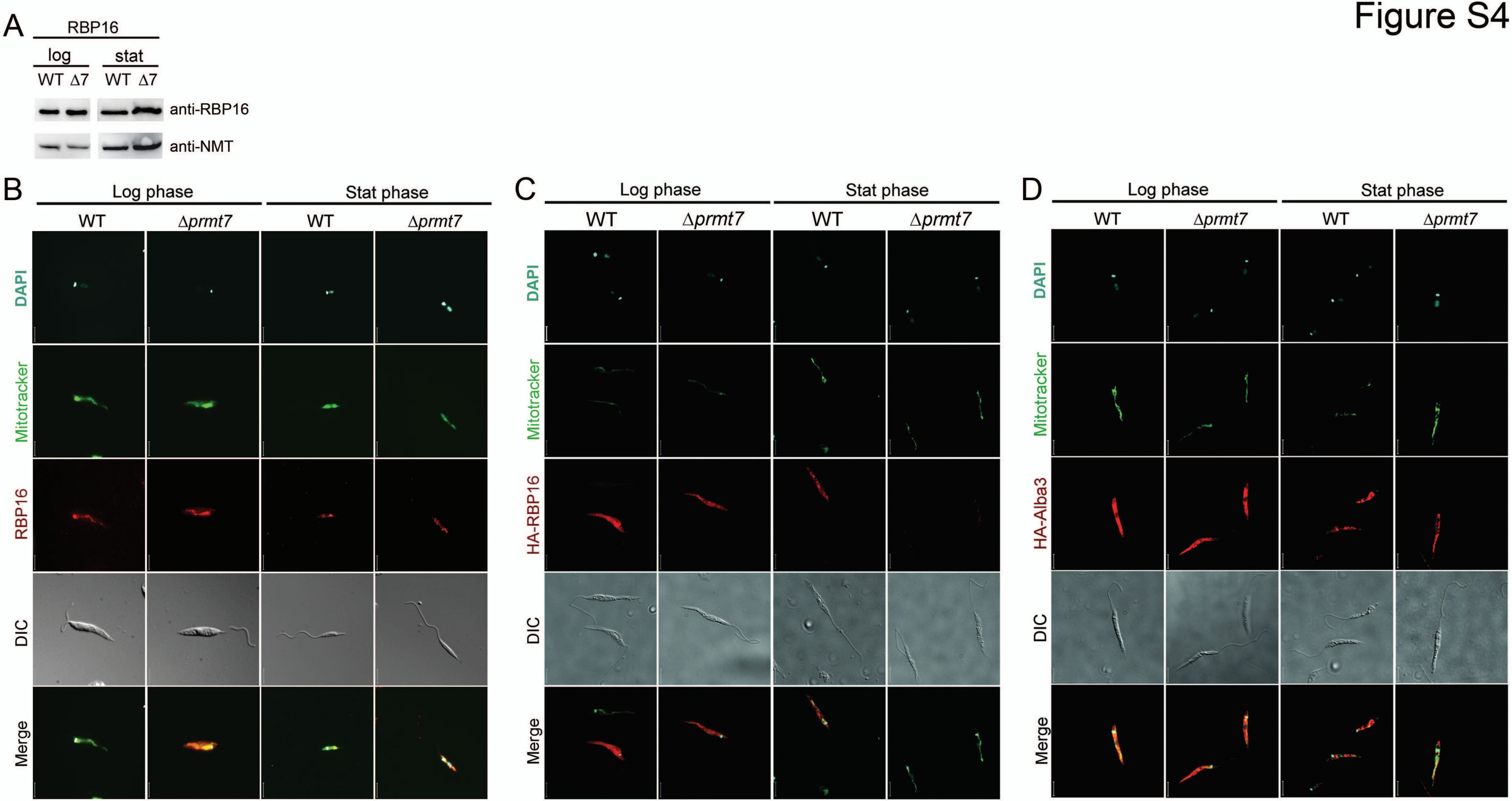

A
RBP16
log
stat
WT Δ7 WT Δ7
anti-RBP16
anti-NMT
B
Log phase
Stat phase
WT
Δprmt7
DAPI
Mitotracker
RBP16
DIC
Merge
C
Log phase
Stat phase
WT
Δprmt7
DAPI
Mitotracker
HA-RBP16
DIC
Merge
D
Log phase
Stat phase
WT
Δprmt7
DAPI
Mitotracker
HA-Alba3
DIC
Merge

**Supplemental Table S1.** Global monomethyl arginine peptides identified and quantified by heavy methyl SILAC analysis in WT and Δ*prmt7 Leishmania major*.

| Gene ID | Product description | log2FC WT/Δ7 (methylpeptides) | q-value (H&B test) | Methylpeptide sequence* | Modification site[$] |
|---|---|---|---|---|---|
| LmjF.27.1680 | hypothetical protein, conserved (NXF1) | 41.28 | 0.00E+00 | SGPGSGGGGVGILGNGNNNGSRGGGR +3 Methyl x 2 | R26 Methyl |
| LmjF.35.2200 | RNA-binding protein, putative | 39.62 | 0.00E+00 | HNMAGSAYGRGAPYQVGAAESTEMNGTEPLPKPR +5 Methyl x | M3 Oxidation, R10 Methyl, M24 Oxidation |
| LmjF.16.0730 | ubiquitin hydrolase, putative | 38.04 | 0.00E+00 | SGGGRSGGRGAGATNAPSsPVAPSPSETVTEASAPPEPQTLEGR + | ethyl:2H(3)13C(1), R9 Methyl:2H(3)13 |
| LmjF.10.0880 | hypothetical protein, conserved | 37.47 | 0.00E+00 | GRGCGAEDAGPSAARGAGIPAVALR +4 Methyl x 2 | ethyl:2H(3)13C(1), R15 Methyl:2H(3)13 |
| LmjF.16.0730 | ubiquitin hydrolase, putative | 35.85 | 0.00E+00 | SGGGRSGGRGAGATNAPSSPVAPSPSETVTEASAPPEPQTLEGR + | ethyl:2H(3)13C(1), R9 Methyl:2H(3)13 |
| LmjF.35.2200 | RNA-binding protein, putative | 35.33 | 0.00E+00 | HNMAGsAYGRGAPYQVGAAESTEMNGTEPLPKPR +4 Methyl x | M3 Oxidation, R10 Methyl, M24 Oxidation |
| LmjF.34.1110 | Shwachman-Bodian-Diamond syndrome (SBDS) protein/SBDS protein C-terminal domain conta | 35.19 | 0.00E+00 | LGLDPSHDLQDDsDDDGGRGRGSR +3 Methyl x 2 Phos x 1 | S13 Phospho, R21 Methyl |
| LmjF.25.0540 | hypothetical protein SCD6.10 (SCD6) | 35.17 | 0.00E+00 | GRGGRAAAAAPAASATAASSAATR +4 Methyl x 2 | R2 Methyl:2H(3)13C(1), R5 Methyl:2H(3)13C(1) |
| LmjF.25.0540 | hypothetical protein SCD6.10 (SCD6) | 34.51 | 0.00E+00 | GRGGRAAAAAPAASATAASSAATR +3 Methyl x 2 | R2 Methyl:2H(3)13C(1), R5 Methyl:2H(3)13C(1) |
| LmjF.19.0380 | hypothetical protein, conserved | 34.32 | 0.00E+00 | RVPDSGASMASSGAAGRGAGRGR +4 Methyl x 2 | R17 Methyl, R21 Methyl, R23 Methyl |
| LmjF.01.0260 | Calmodulin-binding, putative | 34.10 | 0.00E+00 | AQYDHVSSHGYGRAGSGAAAPSGSR +3 Methyl x 1 | R13 Methyl |
| LmjF.14.0570 | WD domain, G-beta repeat, putative | 7.33 | 1.80E-18 | GGSSYGRGEHATSGR +3 Methyl x 1 | R7 Methyl:2H(3)13C(1) |
| LmjF.34.2580 | ALBA-domain protein 3 (Alba3) | 7.15 | 1.15E-02 | GGRGVAAIDRR +3 Methyl x 1 | R3 Methyl:2H(3)13C(1) |
| LmjF.25.0290 | RNA-binding protein, putative | 6.70 | 4.96E-16 | SVARPPPPPPPIGRGR +4 Methyl x 2 | R14 Methyl, R16 Methyl |
| LmjF.27.1680 | hypothetical protein, conserved (NXF1) | 6.70 | 1.94E-16 | SGPGSGGGGVGILGNGNNNGSR +2 Methyl x 1 | R22 Methyl |
| LmjF.27.0130 | WW domain/Zinc finger C-x8-C-x5-C-x3-H type (and similar), putative (ZFP3) | 6.58 | 2.15E-07 | TTHWSIPPTAYEPYYNNGR +3 Methyl x 1 | R19 Methyl |
| LmjF.32.2410 | hypothetical protein, conserved | 6.54 | 3.79E-03 | TSTGRAGSAASGPLLHR +3 Methyl x 1 | R5 Methyl:2H(3)13C(1) |
| LmjF.32.1800 | hypothetical protein, conserved | 6.28 | 2.88E-16 | SGRGSARGGGSSHNR +3 Methyl x 2 | R3 Methyl:2H(3)13C(1), R7 Methyl:2H(3)13C(1) |
| LmjF.10.0280 | hypothetical protein, conserved | 5.80 | 8.74E-14 | GGGRGGSLSR +3 Methyl x 1 | R4 Methyl:2H(3)13C(1) |
| LmjF.27.0130 | WW domain/Zinc finger C-x8-C-x5-C-x3-H type (and similar), putative (ZFP3) | 5.38 | 3.17E-05 | TTHWSIPPTAYEPYYNNGRGR +4 Methyl x 2 | R19 Methyl:2H(3)13C(1), R21 Methyl:2H(3)13C(1) |
| LmjF.18.1420 | pumilio protein 2, putative (PUF2) | 4.89 | 3.54E-13 | DNRPGGGVGSSSGGNNNGSGGRAGR +3 Methyl x 1 | R22 Methyl |
| LmjF.26.0730 | hypothetical protein, conserved | 4.63 | 4.23E-12 | TSSGRGGGsRENDVDQPAEQQR +3 Methyl x 1 Phos x 1 | R5 Methyl |
| LmjF.10.1110 | PAB1-binding protein , putative (PBP1) | 4.53 | 4.97E-09 | KEAAAATSAAPAAPAATPAATTATAGRGAAR +4 Methyl x 1 | R27 Methyl |
| LmjF.34.1110 | Shwachman-Bodian-Diamond syndrome (SBDS) protein/SBDS protein C-terminal domain conta | 4.44 | 4.14E-11 | LGLDPSHDLQDDsDDDGGRGR +3 Methyl x 1 Phos x 1 | S13 Phospho, R19 Methyl:2H(3)13C(1) |
| LmjF.19.0190 | multiprotein-bridging factor 1, putative (MBF1) | 4.39 | 5.24E-11 | PRGAIPTGQDWEEQR +3 Methyl x 1 | R2 Methyl |
| LmjF.19.0190 | multiprotein-bridging factor 1, putative (MBF1) | 4.03 | 1.81E-05 | PRGAIPTGQDWEEQRFNFQQR +4 Methyl x 1 | R2 Methyl |
| LmjF.25.0290 | RNA-binding protein, putative | 3.92 | 1.67E-12 | SVARPPPPPPPIGR +3 Methyl x 1 | R14 Methyl:2H(3)13C(1) |
| LmjF.04.0570 | DENN (AEX-3) domain containing protein, putative | 3.88 | 6.80E-06 | AYGRGLGIESEMFDPVAR +3 Methyl x 1 | R4 Methyl:2H(3)13C(1), M12 Label:13C(1)2H(3) |
| LmjF.27.1300 | KH domain containing protein, putative | 3.69 | 2.41E-13 | GGRGGGRGAGAPNGDDAAR +3 Methyl x 2 | R3 Methyl, R7 Methyl |
| LmjF.15.0620 | hypothetical protein, conserved | 3.64 | 3.24E-12 | ASSHDNGSNGGAVIGRGGGNAR +3 Methyl x 1 | R16 Methyl |
| LmjF.22.0930 | mitochondrial chaperone, putative | 3.45 | 7.96E-12 | SDGAPAVGRGGGGSTGGR +2 Methyl x 1 | R18 Methyl |
| LmjF.24.0820 | inositol polyphosphate phosphatase, putative | 3.22 | 9.58E-08 | GRGGAAGMHTSAAGPSAR +3 Methyl x 1 | R2 Methyl |
| LmjF.10.1110 | PAB1-binding protein , putative (PBP1) | 2.97 | 5.67E-08 | EAAAATSAAPAAPAATPAATTATAGRGAAR +3 Methyl x 1 | R26 Methyl |
| LmjF.24.1180 | Surfeit locus protein 6, putative (SURF6) | 2.77 | 2.44E-07 | GARPFAPQGRGGGASSR +3 Methyl x 1 | #N/A |
| LmjF.26.0730 | hypothetical protein, conserved | 2.61 | 7.62E-07 | TSSGRGGGSRENDVDQPAEQQR +3 Methyl x 1 | R5 Methyl |
| LmjF.24.0710 | Protein of unknown function (DUF523), putative | 2.60 | 1.29E-04 | RPAASASAQRR +3 Methyl x 1 | #N/A |
| LmjF.25.1740 | mitochondrial RNA binding complex 1 subunit, putative | 2.34 | 2.94E-06 | SSGGGRGSGSSFGPPVGSHGTGSFGNPR +3 Methyl x 1 | R6 Methyl:2H(3)13C(1) |
| LmjF.21.1350 | PB1 domain containing protein, putative | 2.15 | 2.28E-03 | QASVSSTLTTSAGGRGGAMAAR +3 Methyl x 1 | R15 Methyl:2H(3)13C(1), M19 Label:13C(1)2H(3) |
| LmjF.22.0930 | mitochondrial chaperone, putative | 2.05 | 3.87E-07 | SDGAPAVGRGGGGSTGGR +3 Methyl x 1 | R18 Methyl |
| LmjF.36.5100 | hypothetical protein, conserved (PUF11) | 1.99 | 1.16E-03 | GGYYQGGGAYGGAGIQQGYPGAAAPISR +3 Methyl x 1 | R28 Methyl |
| LmjF.19.0190 | multiprotein-bridging factor 1, putative (MBF1) | 1.79 | 5.55E-03 | FNFQQRGSGGSAQPR +3 Methyl x 1 | R6 Methyl:2H(3)13C(1) |
| LmjF.25.1740 | mitochondrial RNA binding complex 1 subunit, putative | 1.75 | 1.15E-03 | SSGGGRGSGSSFGPPVGSHGTGSFGNPR +4 Methyl x 1 | R6 Methyl:2H(3)13C(1) |
| LmjF.22.0930 | mitochondrial chaperone, putative | 1.74 | 1.83E-11 | SDGAPAVGRGGGGSTGGRR +3 Methyl x 1 | R9 Methyl |
| LmjF.05.0850 | zinc-finger of a C2HC-type, putative | 1.59 | 1.16E-03 | GSGGSGGGGRGGRPAWNSDVEVR +4 Methyl x 1 | R13 Methyl |
| LmjF.14.0570 | WD domain, G-beta repeat, putative | 1.56 | 6.34E-04 | GGSSYGRGEHATSGR +2 Methyl x 1 | R7 Methyl:2H(3)13C(1) |
| LmjF.33.1150 | pumilio protein 6, putative (PUF6) | 1.49 | 2.34E-04 | GSYEGEMAGGVTYGRGTGPTAK +3 Methyl x 1 | R15 Methyl |
| LmjF.25.0540 | hypothetical protein SCD6.10 (SCD6) | 1.44 | 1.49E-03 | AAAAAPAASATAASSAATRGSR +3 Methyl x 1 | R19 Methyl |
| LmjF.25.0540 | hypothetical protein SCD6.10 (SCD6) | 1.33 | 4.90E-03 | KADTETFGPEMVSSMRGFR +3 Methyl x 1 | M11 Label:13C(1)2H(3), M15 Label:13C(1)2H(3), R16 Methyl:2H(3)13C(1) |
| LmjF.32.1800 | hypothetical protein, conserved | 1.26 | 1.19E-04 | RGGASWAANTDTTPPGGK +3 Methyl x 1 | R1 Methyl |
| LmjF.33.0260 | RGG-containing protein 2, putative (RGG2) | 1.25 | 1.52E-04 | DSSDGWGGGRGGGHSGWGGDGGWKDAPTGDR +5 Methyl x | R10 Methyl:2H(3)13C(1) |
| LmjF.08.0520 | hypothetical protein, conserved | 1.19 | 2.37E-02 | SSRGGAAAQAR +2 Methyl x 1 | R3 Methyl |
| LmjF.30.0090 | hypothetical protein, conserved | 1.12 | 3.00E-04 | IEGFGRGGTQVSR +3 Methyl x 1 | R6 Methyl |
| LmjF.15.1310 | MGT1 magnesium transporter | 0.91 | 1.18E-03 | LVRsPLLSRGGAVVGFGSR +3 Methyl x 1 Phos x 1 | S4 Phospho, R9 Methyl |
| LmjF.13.0580 | hypothetical protein, conserved | 0.82 | 1.33E-03 | SGAARGGGIGVSAPSK +3 Methyl x 1 | R5 Methyl:2H(3)13C(1) |
| LmjF.36.5850 | flagellum targeting protein kharon1, putative | 0.81 | 2.88E-03 | RAsLHNLMYASGDAYGAVASRGQGASR +4 Methyl x 1 Phos x 1 | S3 Phospho, M8 Label:13C(1)2H(3), R27 Methyl:2H(3)13C(1) |
| LmjF.31.2360 | hypothetical protein, conserved | 0.73 | 1.85E-02 | NYPNLSDPPTEEEALGSRGGTGGAYGSR +3 Methyl x 1 | R28 Methyl |
| LmjF.28.0825 | RNA binding protein rbp16, putative (RBP16) | 0.67 | 2.34E-02 | LPSGPRPPEGSPSGR +3 Methyl x 1 | #N/A |
| LmjF.33.1150 | pumilio protein 6, putative (PUF6) | 0.64 | 2.02E-02 | YNNNNSAGGGVGGGRVGR +3 Methyl x 1 | R15 Methyl |
| LmjF.05.0850 | zinc-finger of a C2HC-type, putative | 0.63 | 5.07E-03 | GSGGSGGGGRGGRPAWNSDVEVR +3 Methyl x 1 | R13 Methyl |
| LmjF.24.1430 | kinesin, putative | 0.60 | 1.29E-03 | NGGAAAAGGRGGCNAPALLTDAPR +3 Methyl x 1 | R10 Methyl:2H(3)13C(1) |
| LmjF.35.3100 | ATP-dependent RNA helicase, putative (DED1) | 0.475378065 | 2.33E-02 | NYDDGDGGEGGYSRNYDSGYGGGGYR +3 Methyl x 1 | R14 Methyl |
| LmjF.32.0400 | ATP-dependent RNA helicase HEL67 (DDX3) | 0.398086975 | 3.12E-03 | KPNVQNQPR +2 Methyl x 1 | R9 Methyl |
| LmjF.36.4590 | PHD-like zinc-binding domain containing protein, putative | 0.38904051 | 3.98E-02 | NSGGGTTDSAGALSSSPTARGGAGVGGGHAR +3 Methyl x 1 | R31 Methyl:2H(3)13C(1) |
| LmjF.21.1030 | hypothetical protein, conserved | 0.352663399 | 4.92E-02 | GAGGPSRPPFAVDDDANySDGGGRGGR +3 Methyl x 1 Phos x | R27 Methyl:2H(3)13C(1) |
| LmjF.04.0920 | hypothetical protein | 0.319371052 | 8.31E-03 | SRDAGCSPQALRGGIR +3 Methyl x 1 | R12 Methyl |
| LmjF.17.1030 | hypothetical protein, conserved | 0.232130541 | 5.40E-03 | SSGRGGVPPGQPSAVAGGGSPGQLNR +3 Methyl x 1 | R4 Methyl |
| LmjF.34.1360 | Nucleoporin NUP225 | 0.212786983 | 2.26E-04 | SGSAVNRGGSAAALEAYYGYQQR +3 Methyl x 1 | R7 Methyl |
| LmjF.31.2360 | hypothetical protein, conserved | 0.161499462 | 3.61E-03 | LQRGGYGQALQGR +2 Methyl x 1 | R3 Methyl |
| LmjF.35.1510 | NLI interacting factor-like phosphatase/Zinc finger C-x8-C-x5-C-x3-H type (and similar), putative | 0.126983206 | 2.20E-03 | ANNSNDHQHQRGGGAR +3 Methyl x 1 | R11 Methyl |
| LmjF.36.0490 | zinc-finger of a C2HC-type, putative | -0.60 | 2.30E-03 | FSSRGGGGGMGGGGGGGR +2 Methyl x 1 | R4 Methyl:2H(3)13C(1), M10 Label:13C(1)2H(3)+Oxidation |
| LmjF.32.0850 | polypyrimidine tract-binding protein, putative (DRBD4) | -0.75 | 9.35E-03 | DRGGCGAGSDCAKDSDPAVQR +3 Methyl x 1 | R2 Methyl |
| LmjF.23.0080 | hypothetical protein, conserved | -0.79 | 2.91E-03 | AVSAQVQARGR +2 Methyl x 1 | R9 Methyl |
| LmjF.36.4260 | hypothetical protein, conserved | -0.81 | 9.04E-03 | SSMARGGASPQR +3 Methyl x 1 | M3 Label:13C(1)2H(3), R5 Methyl:2H(3)13C(1) |
| LmjF.07.0340 | ATP-dependent RNA helicase DBP2B, putative | -1.05 | 4.65E-03 | SGGGYGGGRGYGGGYGGGR +2 Methyl x 1 | R9 Methyl |
| LmjF.05.0850 | zinc-finger of a C2HC-type, putative | -1.70 | 1.99E-07 | GAGGGGGRGGGADAAAGAAK +3 Methyl x 1 | R8 Methyl:2H(3)13C(1) |
| LmjF.05.0850 | zinc-finger of a C2HC-type, putative | -1.83 | 2.01E-06 | GAGGGGGRGGGADAAAGAAK +2 Methyl x 1 | R8 Methyl:2H(3)13C(1) |
| LmjF.30.0760 | hypothetical protein, conserved | -2.50 | 9.72E-07 | RGGSGGDVAVAGGNEGR +3 Methyl x 1 | R1 Methyl:2H(3)13C(1) |
| LmjF.15.0800 | hypothetical protein, conserved | -2.50 | 6.17E-03 | RRGGGDSGAVDR +3 Methyl x 1 | R1 Methyl |
| LmjF.32.0840 | hypothetical protein, conserved (DRBD18) | -3.01 | 4.02E-04 | GGRGGSFGGFSQQPTNTISIR +3 Methyl x 1 | R3 Methyl |
| LmjF.07.0340 | ATP-dependent RNA helicase DBP2B, putative | -3.57 | 7.25E-09 | DGGYGGGYGGGRGGDR +3 Methyl x 1 | R12 Methyl:2H(3)13C(1) |
| LmjF.36.6980 | eukaryotic translation initiation factor 3 subunit c | -3.62 | 1.35E-08 | GRGGMAGGR +2 Methyl x 1 | R2 Methyl:2H(3)13C(1), M5 Label:13C(1)2H(3) |
| LmjF.25.0540 | hypothetical protein SCD6.10 (SCD6) | -3.65 | 7.62E-08 | DPAIVEVHAPARGR +3 Methyl x 1 | R14 Methyl |
| LmjF.32.1800 | hypothetical protein, conserved | -3.83 | 1.17E-03 | GGQGIRGGHSAQR +3 Methyl x 1 | R6 Methyl:2H(3)13C(1) |
| LmjF.27.1300 | KH domain containing protein, putative | -3.96 | 5.52E-11 | GGGRGAGAPNGDDAAR +2 Methyl x 1 | R4 Methyl:2H(3)13C(1) |
| LmjF.34.4290 | nucleolar protein family a, putative | -4.11 | 1.44E-10 | LFTEPPKPR +3 Methyl x 1 | R9 Methyl:2H(3)13C(1) |
| LmjF.27.1680 | hypothetical protein, conserved (NXF1) | -4.66 | 4.15E-10 | SGPGSGGGGVGILGNGNNNGSRGGGR +3 Methyl x 1 | R22 Methyl |
| LmjF.36.5100 | hypothetical protein, conserved (PUF11) | -5.37 | 8.50E-14 | GGPGGLQRGGR +2 Methyl x 1 | R8 Methyl:2H(3)13C(1) |
| LmjF.31.0080 | hypothetical protein, conserved (ZC3H34) | -5.82 | 6.18E-07 | GGWYYPSAGSGGYNNMR +2 Methyl x 1 | M16 Label:13C(1)2H(3), R17 Methyl:2H(3)13C(1) |
| LmjF.01.0210 | CUE domain/Domain of unknown function (DUF1771)/Smr domain containing protein, putative | -3.88 | 3.32E-01 | RGGSGQQDNVAR +2 Methyl x 1 | R1 Methyl |
| LmjF.01.0210 | CUE domain/Domain of unknown function (DUF1771)/Smr domain containing protein, putative | 0.00 | 1.00E+00 | RGGSGQQDNVAR +3 Methyl x 1 | R1 Methyl |
| LmjF.01.0260 | Calmodulin-binding, putative | 0.00 | 1.00E+00 | LGSPCAPGASAASGGGGGTSDASGGAASCGVGAPNR +3 Methyl | R36 Methyl |
| LmjF.01.0260 | Calmodulin-binding, putative | 0.00 | 1.00E+00 | LGSPCAPGASAASGGGGGTSDASGGAASCGVGAPNRAR +3 Met | R36 Methyl:2H(3)13C(1) |
| LmjF.01.0540 | hypothetical protein, conserved | -0.42 | 6.83E-01 | RGGVYVPYLHDTILFCEPLPR +4 Methyl x 1 | R1 Methyl:2H(3)13C(1) |
| LmjF.01.0620 | hypothetical protein, conserved | 2.69 | 8.39E-01 | QARGLGNTSGGASAIR +3 Methyl x 1 | R3 Methyl |
| LmjF.01.0680 | hypothetical protein, unknown function | 0.35 | 1.00E+00 | ARGGCTVQR +2 Methyl x 1 | R2 Methyl:2H(3)13C(1) |

| | | | | | |
|---|---|---|---|---|---|
| LmjF.01.0740 | hypothetical protein, conserved | 0.00 | 1.00E+00 | RSRGGAAALR +3 Methyl x 1 | R3 Methyl:2H(3)13C(1) |
| LmjF.01.0740 | hypothetical protein, conserved | 0.06 | 1.07E+00 | SRGGAAALR +2 Methyl x 1 | R2 Methyl:2H(3)13C(1) |
| LmjF.02.0530 | hypothetical protein, conserved | 0.00 | 1.00E+00 | IAANGRGCGNPDR +3 Methyl x 1 | R6 Methyl:2H(3)13C(1) |
| LmjF.02.0560 | hypothetical protein, unknown function | -0.73 | 8.39E-01 | GVDGVSGTRGGGGQPIMIPR +3 Methyl x 1 | R9 Methyl:2H(3)13C(1), M17 Label:13C(1)2H(3) |
| LmjF.02.0660 | Nop14-like family, putative | 0.99 | 1.00E+00 | SGHGRGAGGYTASEMEVR +3 Methyl x 1 | R5 Methyl |
| LmjF.03.0260 | hypothetical protein, conserved | -1.00 | 1.09E+00 | YGVSPKAGVDGR +2 Methyl x 1 | #N/A |
| LmjF.03.0350 | protein kinase, putative | 1.85 | 1.00E+00 | EVVAAASSAGAVAGRGANMSR +3 Methyl x 1 | #N/A |
| LmjF.03.0560 | hypothetical protein | -1.28 | 8.55E-01 | AYGVTGGVAATSPAARGGACGTMPLSTK +3 Methyl x 1 | R16 Methyl:2H(3)13C(1), M23 Label:13C(1)2H(3) |
| LmjF.03.0600 | arginine N-methyltransferase, putative (PRMT3) | -0.11 | 9.43E-01 | TGAPTQRGESR +2 Methyl x 1 | R7 Methyl |
| LmjF.03.0800 | 6-phosphofructo-2-kinase 1 | -1.13 | 5.01E-01 | VGRGGATGGAVASDsLEEEALLR +3 Methyl x 1 Phos x 1 | R3 Methyl |
| LmjF.03.0800 | 6-phosphofructo-2-kinase 1 | -0.63 | 2.08E-01 | VGRGGATGGAVASDSLEEEALLR +3 Methyl x 1 | R3 Methyl |
| LmjF.03.0800 | 6-phosphofructo-2-kinase 1 | -0.34 | 1.06E+00 | SRGTGSLVSAASQLATHAR +4 Methyl x 1 | R2 Methyl |
| LmjF.03.0800 | 6-phosphofructo-2-kinase 1 | 0.00 | 1.00E+00 | RVGRGGATGGAVASDSLEEEALLR +4 Methyl x 1 | R4 Methyl |
| LmjF.03.0800 | 6-phosphofructo-2-kinase 1 | 0.00 | 1.00E+00 | SRGTGSLVSAASQLATHAR +3 Methyl x 1 | R2 Methyl |
| LmjF.03.0820 | hypothetical protein, conserved | -0.15 | 1.00E+00 | ARGGGVAVQQQR +2 Methyl x 1 | R2 Methyl |
| LmjF.04.0830 | hypothetical protein, conserved | -0.61 | 1.05E+00 | GRGGLTQPR +2 Methyl x 1 | R2 Methyl:2H(3)13C(1) |
| LmjF.04.0920 | hypothetical protein | 0.66 | 1.20E-01 | DAGCSPQALRGGIR +3 Methyl x 1 | R10 Methyl |
| LmjF.05.0140 | nucleolar RNA helicase II, putative | -2.30 | 1.68E-01 | GGFNYGGGRGGFGGYGGNR +3 Methyl x 1 | R9 Methyl:2H(3)13C(1) |
| LmjF.05.0140 | nucleolar RNA helicase II, putative | -0.62 | 8.17E-01 | GGRGGFNYGGGR +2 Methyl x 1 | #N/A |
| LmjF.05.0300 | hypothetical protein, conserved | -1.01 | 1.03E+00 | GSAPDNSAVGGRGR +3 Methyl x 1 | #N/A |
| LmjF.05.0370 | hypothetical protein, conserved | 0.25 | 8.21E-01 | RGGAAATWCAQGIFEDPAVTMAR +3 Methyl x 1 | R1 Methyl |
| LmjF.05.0640 | Ankyrin repeats (many copies)/Ankyrin repeats (3 copies), putative | 0.00 | 1.00E+00 | HQQAPSYAADVMDGDAPGVNYPLGGSTGGSSSARGR +4 Methy | R34 Methyl |
| LmjF.05.0720 | phophatase-like protein | -0.48 | 1.00E+00 | QQLNALIPVMSPSYTAFGPAtPTRGGAK +3 Methyl x 1 Phos x 1 | #N/A |
| LmjF.05.0850 | zinc-finger of a C2HC-type, putative | 2.27 | 8.68E-02 | GSGGSGGGGRGGRPAWNSDVEVR +2 Methyl x 1 | R13 Methyl |
| LmjF.05.0850 | zinc-finger of a C2HC-type, putative | 0.01 | 3.46E-01 | GPAASSSPAAGASGVLGTSAQGRGYGsDNEEAGNNAYMPQQPS | S5 Phospho, R23 Methyl |
| LmjF.05.0850 | zinc-finger of a C2HC-type, putative | 8.42 | 4.62E-01 | SGGGGGGRGAGGGGGRGGGADAAAGAAK +4 Methyl x 2 | R8 Methyl, R16 Methyl |
| LmjF.05.0850 | zinc-finger of a C2HC-type, putative | 8.04 | 1.06E+00 | SGGGGGGRGAGGGGGRGGGADAAAGAAK +3 Methyl x 2 | R8 Methyl, R16 Methyl |
| LmjF.05.1190 | hypothetical protein, conserved | -0.05 | 4.19E-01 | RGGGGNSAAAPPR +3 Methyl x 1 | R1 Methyl:2H(3)13C(1) |
| LmjF.05.1190 | hypothetical protein, conserved | -0.04 | 2.19E-01 | RGGGGNSAAAPPR +2 Methyl x 1 | R1 Methyl:2H(3)13C(1) |
| LmjF.06.0460 | ATP-NAD kinase-like protein | 0.07 | 1.07E+00 | SPLLRGVDTTTR +2 Methyl x 1 | R5 Methyl:2H(3)13C(1) |
| LmjF.06.0550 | hypothetical protein, conserved | -0.35 | 9.78E-01 | QLRGGSGPR +2 Methyl x 1 | R3 Methyl |
| LmjF.06.0640 | STE/STE11 serine/threonine-protein kinase, putative | 0.62 | 1.17E-01 | RGGAGASTPAPsPPLVPLDAAVDVTK +3 Methyl x 1 Phos x 1 | R1 Methyl, S12 Phospho |
| LmjF.06.0810 | hypothetical protein, unknown function | -0.39 | 9.25E-01 | GARGGGGGPSSSSASSPTSR +2 Methyl x 1 | R3 Methyl:2H(3)13C(1) |
| LmjF.06.0810 | hypothetical protein, unknown function | -0.18 | 8.59E-01 | GARGGGGGPSSSSASSPTSR +3 Methyl x 1 | R3 Methyl:2H(3)13C(1) |
| LmjF.06.0940 | STE/STE11 serine/threonine-protein kinase, putative | -0.74 | 1.04E+00 | AAVVSSSNAAAMSSVHGAPDIVVPSPRGR +4 Methyl x 1 | M12 Label:13C(1)2H(3), R27 Methyl:2H(3)13C(1) |
| LmjF.06.0970 | Domain of unknown function (DUF3883), putative | -0.13 | 1.09E+00 | SGTASSALRGSGGGAESR +3 Methyl x 1 | #N/A |
| LmjF.07.0340 | ATP-dependent RNA helicase DBP2B, putative | -0.67 | 2.61E-01 | SGGGYGGGRGYGGGYGGGR +3 Methyl x 1 | R9 Methyl |
| LmjF.07.0340 | ATP-dependent RNA helicase DBP2B, putative | -0.90 | 8.65E-01 | SGGGYGGGRGYGGGYGGGR +3 Methyl x 2 | R9 Methyl |
| LmjF.07.0490 | hypothetical protein, conserved | -0.42 | 2.15E-01 | VVSSSSIHGGRGGTN +2 Methyl x 1 | R11 Methyl:2H(3)13C(1) |
| LmjF.07.0490 | hypothetical protein, conserved | -0.21 | 1.09E+00 | VVSSSSIHGGRGGTN +3 Methyl x 1 | R11 Methyl:2H(3)13C(1) |
| LmjF.07.0740 | hypothetical protein, conserved | 2.49 | 7.84E-01 | AVTAAGRGTVDDHEGAFTSMR +3 Methyl x 1 | #N/A |
| LmjF.07.0870 | splicing factor ptsr1-like protein | -0.19 | 8.54E-01 | RGGYHGSDR +2 Methyl x 1 | R1 Methyl |
| LmjF.07.0940 | Domain of unknown function (DUF3437), putative | 1.46 | 1.00E+00 | LLPATTTAAVESPGYGNGRAGQPLVLSTPAVIGVAPSATLR +4 Me | R19 Methyl:2H(3)13C(1) |
| LmjF.08.0110 | hypothetical protein, unknown function | -1.72 | 1.00E+00 | MGSGLRGGLSPSVPAAQPYYNDIASIR +3 Methyl x 1 | M1 Label:13C(1)2H(3), R6 Methyl:2H(3)13C(1) |
| LmjF.08.0620 | hypothetical protein, conserved | -0.67 | 4.36E-01 | SGAAAVGRGGPSLAGVSLAQSPPLSVVLGGAAAEAASSSSNPR +3 | R8 Methyl, R43 Methyl:2H(3)13C(1) |
| LmjF.08.0620 | hypothetical protein, conserved | 0.98 | 1.06E+00 | SGAAAVGRGGPSLAGVSLAQSPPLSVVLGGAAAEAASSSSNPR +5 | R8 Methyl, R43 Methyl:2H(3)13C(1) |
| LmjF.08.0620 | hypothetical protein, conserved | 1.43 | 3.78E-01 | SPGAVGGGGGRGTTSSSSAAASANTAGANR +3 Methyl x 1 | R11 Methyl |
| LmjF.08.0620 | hypothetical protein, conserved | -0.58 | 8.47E-01 | SGAAAVGRGGPSLAGVSLAQSPPLsVVLGGAAAEAASSSSNPR +4 | R8 Methyl, R43 Methyl:2H(3)13C(1) |
| LmjF.08.0620 | hypothetical protein, conserved | 0.32 | 1.21E-01 | SGAAAVGRGGPSLAGVSLAQSPPLSVVLGGAAAEAASSSSNPR +4 | R8 Methyl, R43 Methyl:2H(3)13C(1) |
| LmjF.08.0880 | SET domain containing protein, putative | 0.66 | 1.04E+00 | HMGIDPLSNQAGTHASGGIHRGGNSDGVR +5 Methyl x 1 | #N/A |
| LmjF.08.1000 | AAA domain (dynein-related subfamily)/von Willebrand factor type A domain containing prote | -0.40 | 4.70E-01 | NSASAAPACGRGR +2 Methyl x 1 | R11 Methyl |
| LmjF.08.1250 | hypothetical protein, conserved | 0.00 | 1.00E+00 | DADGSCWGIVGPSPATQPQQPFAPRGGAGTCDTR +3 Methyl x | R25 Methyl:2H(3)13C(1) |
| LmjF.09.0390 | hypothetical protein, conserved | 1.26 | 1.06E+00 | RGGGGGGGGNSNSGGTHVDPDDFSNTQTPTSSMNGGGGSR +4 | R1 Methyl:2H(3)13C(1), M33 Label:13C(1)2H(3) |
| LmjF.09.0520 | Flagellum attachment zone protein 3 | -1.02 | 7.70E-01 | ASRGGAATPRPSTAPPSPENNAADILADENMR +4 Methyl x 1 | R10 Methyl |
| LmjF.09.0520 | Flagellum attachment zone protein 3 | 0.05 | 1.00E+00 | ASRGGAAtPRPSTAPPSPENNAADILADENMR +4 Methyl x 1 Ph | R10 Methyl |
| LmjF.09.0530 | leucine-rich repeat protein, putative | -0.48 | 1.00E+00 | QRGGGGGDGALASMR +3 Methyl x 1 | #N/A |
| LmjF.09.0540 | hypothetical protein, unknown function | -0.18 | 9.25E-01 | VRGSSGGFGGR +2 Methyl x 1 | R2 Methyl |
| LmjF.09.0540 | hypothetical protein, unknown function | 0.26 | 1.09E+00 | VRGSSGGFGGR +3 Methyl x 1 | R2 Methyl |
| LmjF.09.0620 | hypothetical protein, conserved | -0.12 | 3.44E-01 | AQGTSNPGGRGGAVEPNPDPLLQR +3 Methyl x 1 | R10 Methyl |
| LmjF.09.0750 | acyl-CoA binding protein, putative | 1.23 | 1.00E+00 | LPTPVRGGSPHAAPVSSSPGGIESALAGGCHCSTGIAVGEDALVQN | R6 Methyl:2H(3)13C(1) |
| LmjF.09.1180 | hypothetical protein, conserved | -2.29 | 1.06E+00 | NLGDEGLDSVEVSSSSSESMSPFGCPANTTTTVPNGRGAR +4 Me | #N/A |
| LmjF.09.1440 | DnaJ domain containing protein, putative | -0.08 | 1.07E+00 | NHRGGPPNK +2 Methyl x 1 | R3 Methyl:2H(3)13C(1) |
| LmjF.10.1110 | PAB1-binding protein , putative (PBP1) | 0.00 | 1.00E+00 | RGGMATAAR +2 Methyl x 1 | R1 Methyl |
| LmjF.10.1110 | PAB1-binding protein , putative (PBP1) | -0.35 | 1.00E+00 | KEAAAATSAAPAAPAATPAATTATAGRGAAR +3 Methyl x 1 | R27 Methyl |
| LmjF.10.1130 | hypothetical protein, conserved | -1.23 | 1.05E+00 | MAAAALSFTSR +3 Methyl x 1 | #N/A |
| LmjF.11.0600 | hypothetical protein, conserved | 0.00 | 1.00E+00 | YSEGEHGGAGSRGHGQR +3 Methyl x 1 | R12 Methyl |
| LmjF.11.0820 | hypothetical protein, conserved | -1.84 | 2.71E-01 | QQQMDMNFGGR +2 Methyl x 1 | R11 Methyl |
| LmjF.11.0820 | hypothetical protein, conserved | -1.08 | 4.74E-01 | MQPMMPPPQQYSR +2 Methyl x 1 | R13 Methyl |
| LmjF.11.0820 | hypothetical protein, conserved | -0.54 | 4.80E-01 | QQQMDMNFGGRSQQSYNPAPNTGYPQR +3 Methyl x 1 | M4 Label:13C(1)2H(3), M6 Label:13C(1)2H(3), R11 Methyl:2H(3)13C(1) |
| LmjF.11.0830 | hypothetical protein, conserved | 0.97 | 1.03E+00 | SIGSYARGGGGGAGDK +2 Methyl x 1 | #N/A |
| LmjF.11.1330 | Flagellar-associated PapD-like, putative | -0.24 | 1.21E-01 | GRGSGAGAAASSSSSASK +3 Methyl x 1 | R2 Methyl:2H(3)13C(1) |
| LmjF.11.1330 | Flagellar-associated PapD-like, putative | 0.40 | 1.04E+00 | GRGSGAGAAASSSSSASK +2 Methyl x 1 | R2 Methyl:2H(3)13C(1) |
| LmjF.12.0320 | Myotubularin-related protein, putative | 0.00 | 1.00E+00 | GCLGRGGAPSVGSEAER +3 Methyl x 1 | #N/A |
| LmjF.12.1110 | DnaJ domain containing protein, putative | 0.65 | 1.01E+00 | GARGGGVPPFSSMASPSWDPAAPAAGAAYR +3 Methyl x 1 | #N/A |
| LmjF.12.1180 | hypothetical protein, conserved | 0.82 | 1.00E+00 | SAQRGGTNAANGSGGGGGPTSYR +3 Methyl x 1 | R4 Methyl:2H(3)13C(1) |
| LmjF.13.0580 | hypothetical protein, conserved | -2.64 | 8.73E-01 | WGPSALSATATPTPPSNRAGGAVNNR +3 Methyl x 1 | R18 Methyl |
| LmjF.13.0580 | hypothetical protein, conserved | 0.28 | 3.33E-01 | SGAARGGGIGVSAPSK +2 Methyl x 1 | R5 Methyl:2H(3)13C(1) |
| LmjF.13.0700 | kinesin, putative | 0.00 | 1.00E+00 | QRGGGGAPPLAAAALVQR +3 Methyl x 1 | #N/A |
| LmjF.13.0810 | hypothetical protein, conserved | 0.02 | 1.69E-01 | GAAAAFVGGSASRGGAVNR +3 Methyl x 1 | #N/A |
| LmjF.13.0880 | hypothetical protein, conserved | -0.10 | 2.77E-01 | RGMGSHSQSQQQSR +2 Methyl x 1 | R1 Methyl |
| LmjF.13.0880 | hypothetical protein, conserved | -0.14 | 1.09E+00 | RGMGSHSQSQQQSR +3 Methyl x 1 | R1 Methyl |
| LmjF.13.1130 | Uncharacterised ACR, YagE family COG1723, putative | -0.93 | 1.00E+00 | HSYRGGGGNAAASGYRsDSGLATANCSVALLSR +4 Methyl x 1 P | R4 Methyl, S13 Phospho |
| LmjF.13.1130 | Uncharacterised ACR, YagE family COG1723, putative | -0.46 | 1.00E+00 | DNLEDDDEAALLALSDVDVVGGGVRGGR +3 Methyl x 1 | S15 Phospho, R28 Methyl |
| LmjF.13.1130 | Uncharacterised ACR, YagE family COG1723, putative | -0.45 | 7.60E-01 | HSYRGGGGNAAASGYR +3 Methyl x 1 | R4 Methyl |
| LmjF.13.1130 | Uncharacterised ACR, YagE family COG1723, putative | -0.39 | 8.18E-01 | SLLAPKDNLEDDDEAALLALSDVDVVGGGVRGGR +4 Methyl x 1 | S21 Phospho, R34 Methyl:2H(3)13C(1) |
| LmjF.13.1130 | Uncharacterised ACR, YagE family COG1723, putative | -0.25 | 9.31E-01 | HSYRGGGGNAAASGYR +2 Methyl x 1 | R4 Methyl |
| LmjF.13.1130 | Uncharacterised ACR, YagE family COG1723, putative | 0.00 | 1.00E+00 | SLLAPKDNLEDDDEAALLALSDVDVVGGGVRGGR +3 Methyl x 1 | S21 Phospho, R34 Methyl:2H(3)13C(1) |
| LmjF.14.0570 | WD domain, G-beta repeat, putative | 0.00 | 1.00E+00 | GSRGGSSYGR +2 Methyl x 1 | R3 Methyl |
| LmjF.15.0290 | hypothetical protein, conserved | -0.50 | 8.73E-01 | LLADTLSCFNRGGGGGGAR +3 Methyl x 1 | R11 Methyl |
| LmjF.15.0360 | RNA pseudouridylate synthase, putative | 0.53 | 1.00E+00 | QLTRGGGASSLGAWAAALQPVSGVGAAQPAR +3 Methyl x 1 | R4 Methyl |
| LmjF.15.0860 | hypothetical protein, conserved | -0.82 | 4.20E-01 | SDGPGRGGGVAGPPSGADLFHGSPAEQLLQLR +4 Methyl x 1 | R6 Methyl |
| LmjF.15.0920 | protein phosphatase 2A regulatory subunit, putative | 0.00 | 1.00E+00 | QAAVQHVALGSAGGGSGAAHWSSNSSRGSASR +4 Methyl x 1 | R27 Methyl:2H(3)13C(1) |
| LmjF.15.0920 | protein phosphatase 2A regulatory subunit, putative | 0.00 | 1.00E+00 | QQASTSASAHSRGSCYK +3 Methyl x 1 | R12 Methyl |
| LmjF.15.1200 | STE group serine/threonine-protein kinase, putative | 0.00 | 1.00E+00 | HVAGGGRGGVGGGNASNIHLTAPPSNEGSVYLMMADEHQGDG | #N/A |
| LmjF.15.1310 | MGT1 magnesium transporter | -0.08 | 8.50E-02 | sPLLSRGGAVVGFGSR +3 Methyl x 1 Phos x 1 | S1 Phospho, R6 Methyl |
| LmjF.15.1380 | nucleolar RNA binding protein, putative | 0.38 | 3.32E-01 | MNTSGFNNDRGGR +3 Methyl x 1 | M1 Label:13C(1)2H(3)+Oxidation, R10 Methyl:2H(3)13C(1) |
| LmjF.15.1380 | nucleolar RNA binding protein, putative | 1.16 | 7.76E-01 | GGRGGGGGFK +2 Methyl x 1 | R3 Methyl:2H(3)13C(1) |
| LmjF.15.1380 | nucleolar RNA binding protein, putative | 2.07 | 1.05E+00 | MNTSGFNNDRGGRGGGGGFK +3 Methyl x 2 | M1 Label:13C(1)2H(3), R10 Methyl:2H(3)13C(1), R13 Methyl:2H(3)13C(1) |
| LmjF.16.0660 | hypothetical protein, conserved | 0.52 | 3.95E-01 | CDMTNPSGAGRGTGASSASTSSLR +3 Methyl x 1 | #N/A |
| LmjF.16.0730 | ubiquitin hydrolase, putative | 1.16 | 1.00E+00 | RSGGGRSGGRGAGATNAPsSPVAPSPSETVTEASAPPEPQTLEGR | R6 Methyl:2H(3)13C(1), R10 Methyl:2H(3)13C(1), T15 Phospho |
| LmjF.16.0730 | ubiquitin hydrolase, putative | 0.00 | 1.00E+00 | RSGGGRSGGRGAGATNAPSSPVAPSPSETVTEASAPPEPQTLEGR | R6 Methyl:2H(3)13C(1), R10 Methyl:2H(3)13C(1), T15 Phospho |
| LmjF.16.1230 | hypothetical protein, conserved | 0.10 | 3.95E-01 | GSRPASPSGVASGSRGGR +4 Methyl x 1 | R15 Methyl |
| LmjF.16.1230 | hypothetical protein, conserved | -0.40 | 9.47E-01 | GKGSRPASPSGVASGSRGGR +3 Methyl x 1 | R17 Methyl:2H(3)13C(1) |
| LmjF.16.1230 | hypothetical protein, conserved | 0.12 | 1.09E+00 | GSRPASPSGVASGSRGGR +3 Methyl x 1 | R15 Methyl |

| | | | | | |
|---|---|---|---|---|---|
| LmjF.16.1270 | hypothetical protein, conserved | -0.15 | 8.42E-01 | VALSQPMPCSPRGGGLGGR +3 Methyl x 1 | M7 Label:13C(1)2H(3), R12 Methyl:2H(3)13C(1) |
| LmjF.16.1340 | hypothetical protein, conserved | -1.57 | 1.03E+00 | ESQGTSARGGANGVGSSAR +3 Methyl x 1 | R8 Methyl:2H(3)13C(1) |
| LmjF.17.0380 | Qa-SNARE protein, putative | -0.31 | 1.00E+00 | APHQPATAGVGSSSSSGTGRGGGGNVNSSSAR +4 Methyl x 1 | R20 Methyl |
| LmjF.17.0380 | Qa-SNARE protein, putative | -0.29 | 8.53E-01 | APHQPATAGVGSSSSSGTGRGGGGNVNSSSAR +3 Methyl x 1 | R20 Methyl |
| LmjF.17.0510 | hypothetical protein, unknown function | -0.35 | 8.14E-01 | SAALEGFPGASCEISAGPRGGR +3 Methyl x 1 | R22 Methyl |
| LmjF.17.0550 | RNA-binding protein, putative | 0.36 | 8.57E-02 | GMSRGNNNNSRGAGGNHR +3 Methyl x 2 | R4 Methyl, R11 Methyl |
| LmjF.17.0550 | RNA-binding protein, putative | 4.31 | 1.20E-01 | EGRGMSRGNNNNSR +3 Methyl x 2 | R3 Methyl:2H(3)13C(1), M5 Label:13C(1)2H(3), R7 Methyl:2H(3)13C(1) |
| LmjF.17.0550 | RNA-binding protein, putative | 0.00 | 1.00E+00 | GRGGAHLQPQQNFQQPQQYQQHQLHPPLPPPPPR +5 Methyl | R2 Methyl |
| LmjF.18.0390 | hypothetical protein, conserved | 0.00 | 1.00E+00 | AIAGRGGVVVGAPPR +3 Methyl x 1 | #N/A |
| LmjF.18.0650 | PA26 p53-induced protein (sestrin), putative | 0.00 | 1.00E+00 | GVHRGGGVSAGGR +3 Methyl x 1 | R4 Methyl |
| LmjF.18.0800 | Ribosomal protein S8, putative | 0.01 | 4.20E-01 | VYGGGATTAFVPGSSSSSSSSSTASVGADYLRGR +3 Methyl x 1 | R33 Methyl |
| LmjF.18.0840 | hypothetical protein, conserved | -0.71 | 1.03E+00 | GSFGTGRGGASTK +2 Methyl x 1 | R7 Methyl |
| LmjF.18.1240 | pre-RNA processing PIH1/Nop17, putative | 1.12 | 4.82E-01 | DRLNSAAAASSGRGGGSGEAAAR +4 Methyl x 1 | R13 Methyl:2H(3)13C(1) |
| LmjF.18.1240 | pre-RNA processing PIH1/Nop17, putative | 1.03 | 8.32E-01 | LNSAAAASSGRGGGSGEAAAR +3 Methyl x 1 | R11 Methyl:2H(3)13C(1) |
| LmjF.18.1420 | pumilio protein 2, putative (PUF2) | 0.51 | 8.63E-01 | DNRPGGGVGSSSGGNNNGSGGRAGRGGR +4 Methyl x 2 | R22 Methyl:2H(3)13C(1), R25 Methyl:2H(3)13C(1) |
| LmjF.18.1420 | pumilio protein 2, putative (PUF2) | 10.97 | 1.00E+00 | AGRGGRGGNNNNNSSNNSNNQHSDGK +3 Methyl x 2 | R3 Methyl:2H(3)13C(1), R6 Methyl:2H(3)13C(1) |
| LmjF.18.1420 | pumilio protein 2, putative (PUF2) | 10.97 | 1.00E+00 | DNRPGGGVGSSSGGNNNGSGGRAGRGGR +3 Methyl x 2 | R22 Methyl:2H(3)13C(1), R25 Methyl:2H(3)13C(1) |
| LmjF.18.1420 | pumilio protein 2, putative (PUF2) | 10.97 | 1.00E+00 | DNRPGGGVGSSSGGNNNGSGGRAGRGGR +4 Methyl x 3 | R22 Methyl:2H(3)13C(1), R25 Methyl:2H(3)13C(1) |
| LmjF.18.1420 | pumilio protein 2, putative (PUF2) | 0.00 | 1.00E+00 | DNRPGGGVGSSSGGNNNGSGGR +3 Methyl x 1 | R22 Methyl |
| LmjF.18.1420 | pumilio protein 2, putative (PUF2) | 0.00 | 1.00E+00 | DNRPGGGVGSSSGGNNNGSGGRAGR +3 Methyl x 2 | R22 Methyl |
| LmjF.18.1420 | pumilio protein 2, putative (PUF2) | 0.00 | 1.00E+00 | GGRGGNNNNNSSNNSNNQHSDGKNNAMR +4 Methyl x 1 | R3 Methyl |
| LmjF.18.1420 | pumilio protein 2, putative (PUF2) | 0.00 | 1.03E+00 | DNRPGGGVGSSSGGNNNGSGGRAGR +4 Methyl x 1 | R22 Methyl |
| LmjF.19.0060 | 40S ribosomal protein S2 | 0.15 | 1.00E+00 | GRGGPGEEKEWVPCTK +3 Methyl x 1 | R2 Methyl:2H(3)13C(1) |
| LmjF.19.0270 | hypothetical protein, conserved | -0.32 | 1.00E+00 | YGHAPDAVIAAIGRGSGSGNGGGAAAADNSPSAPFHTNGGCGSN | R14 Methyl:2H(3)13C(1) |
| LmjF.19.0270 | hypothetical protein, conserved | 0.00 | 1.00E+00 | KYGHAPDAVIAAIGRGSGSGNGGGAAAADNSPSAPFHTNGGCGS | R15 Methyl:2H(3)13C(1) |
| LmjF.19.0380 | hypothetical protein, conserved | 10.97 | 1.00E+00 | RVPDSGASMASSGAAGRGAGRGR +4 Methyl x 3 | R17 Methyl, R21 Methyl, R23 Methyl |
| LmjF.19.0430 | hypothetical protein, conserved | -0.92 | 5.59E-02 | DGRGGAAPLFCSSSSSSGGK +3 Methyl x 1 | R3 Methyl:2H(3)13C(1) |
| LmjF.19.0800 | ABC transport system ATP-binding protein, putative | -1.34 | 5.82E-02 | MTRGGLTDAVVKDSR +3 Methyl x 1 | R3 Methyl |
| LmjF.19.1020 | tRNA pseudouridine synthase A-like protein | -2.13 | 4.55E-01 | AAASSTGSGRGGPVVSSAYSAEHEPLTSHGLR +5 Methyl x 1 | R10 Methyl |
| LmjF.19.1020 | tRNA pseudouridine synthase A-like protein | 0.32 | 8.22E-01 | AAASSTGSGRGGPVVSSAYSAEHEPLTSHGLR +4 Methyl x 1 | R10 Methyl |
| LmjF.19.1090 | hypothetical protein, unknown function | -0.60 | 1.08E+00 | TARGRGAAGGAAR +3 Methyl x 2 | R3 Methyl:2H(3)13C(1), R5 Methyl:2H(3)13C(1) |
| LmjF.19.1130 | hypothetical protein, conserved | -0.64 | 4.81E-01 | ALLFEQVAAQsQVLVAsR +2 Methyl x 1 Phos x 1 | #N/A |
| LmjF.20.0360 | Protein of unknown function (DUF2946), putative | -0.50 | 9.44E-01 | TLRGGP +2 Methyl x 1 | #N/A |
| LmjF.20.1080 | WD40 repeat-containing protein | -1.18 | 9.25E-01 | LTGGSDMVGNLGRGGWLR +3 Methyl x 1 | M7 Label:13C(1)2H(3), R13 Methyl:2H(3)13C(1) |
| LmjF.20.1240 | Raptor N-terminal CASPase like domain containing protein, putative | -0.20 | 8.61E-01 | TGARGGTSSTATTR +2 Methyl x 1 | R4 Methyl |
| LmjF.21.0270 | STE group serine/threonine-protein kinase, putative | 0.02 | 4.09E-01 | EVGRGGSALVYR +3 Methyl x 1 | #N/A |
| LmjF.21.0270 | STE group serine/threonine-protein kinase, putative | 0.45 | 7.73E-02 | SLTVVVNPCGGRGGGELNSTWDK +3 Methyl x 1 | R12 Methyl |
| LmjF.21.0490 | DnaJ protein, putative | 0.96 | 1.00E+00 | GRQAHAEEDEFEDVtDDDDDEQQQYFR +4 Methyl x 1 Phos x 1 | R2 Methyl, T15 Phospho |
| LmjF.21.0490 | DnaJ protein, putative | 0.32 | 1.00E+00 | GRQAHAEEDEFEDVtDDDDDEQQQYFR +3 Methyl x 1 Phos x 1 | R2 Methyl, T15 Phospho |
| LmjF.21.1030 | hypothetical protein, conserved | -0.70 | 8.60E-01 | GAGGPSRPPFAVDDDANYSDGGGRGGR +4 Methyl x 1 | R27 Methyl:2H(3)13C(1) |
| LmjF.21.1030 | hypothetical protein, conserved | 0.00 | 1.00E+00 | GAGGPSRPPFAVDDDANYSDGGGRGGR +3 Methyl x 1 | R27 Methyl:2H(3)13C(1) |
| LmjF.21.1850 | hypothetical protein, conserved | 0.46 | 1.18E-01 | GRGGAEPPPEER +2 Methyl x 1 | R2 Methyl:2H(3)13C(1) |
| LmjF.22.1320 | hypothetical protein, unknown function | -1.09 | 1.00E+00 | GGNANLSSGEPSVGGRGSVLWGGGGGDGSSGALAPLR +3 Meth | R16 Methyl:2H(3)13C(1) |
| LmjF.23.0080 | hypothetical protein, conserved | -0.56 | 1.09E+00 | AVSAQVQAR +2 Methyl x 1 | R9 Methyl |
| LmjF.23.0090 | Domain of unknown function (DUF1767), putative (TDRD3) | 5.72 | 4.57E-01 | NGGGGGRAQQAGDNYEGR +3 Methyl x 1 | R7 Methyl:2H(3)13C(1) |
| LmjF.23.0090 | Domain of unknown function (DUF1767), putative (TDRD3) | -0.02 | 8.95E-01 | NGGGGGRAQQAGDNYEGR +2 Methyl x 1 | R7 Methyl:2H(3)13C(1) |
| LmjF.23.1010 | hypothetical protein, conserved | 0.60 | 8.13E-01 | QHQQRGGGSGKEPVSSAPNSGGGGSNSNNR +4 Methyl x 1 | R5 Methyl:2H(3)13C(1) |
| LmjF.23.1290 | hypothetical protein, unknown function | -0.23 | 4.74E-01 | RGGGGMASEASFR +2 Methyl x 1 | R1 Methyl:2H(3)13C(1), M6 Label:13C(1)2H(3) |
| LmjF.23.1290 | hypothetical protein, unknown function | 0.35 | 8.75E-01 | RGGGGMASEASFR +3 Methyl x 1 | R1 Methyl:2H(3)13C(1), M6 Label:13C(1)2H(3) |
| LmjF.23.1620 | hypothetical protein | -1.95 | 1.00E+00 | DQGRPRGGGGGSSSAAATFGHGSVVTDCYDQTVSAAELR +5 Me | R4 Methyl |
| LmjF.23.1620 | hypothetical protein | 0.00 | 1.00E+00 | DQGRPRGGGGGSSSAAATFGHGSVVTDCYDQTVSAAELR +4 Me | R4 Methyl |
| LmjF.23.1730 | RING-H2 zinc finger, putative | -0.78 | 1.07E+00 | SRGGDGLHHQGR +3 Methyl x 1 | R2 Methyl |
| LmjF.23.1730 | RING-H2 zinc finger, putative | 10.97 | 1.00E+00 | GGDGLHHQGRGSGQR +3 Methyl x 1 | R10 Methyl:2H(3)13C(1) |
| LmjF.23.1730 | RING-H2 zinc finger, putative | 0.00 | 1.00E+00 | SRGGDGLHHQGR +4 Methyl x 1 | R2 Methyl |
| LmjF.24.1070 | hypothetical protein, conserved | -0.85 | 3.46E-01 | SAAGSPTAAGAATSRGGGPGKGPPR +3 Methyl x 1 | #N/A |
| LmjF.24.1070 | hypothetical protein, conserved | -0.52 | 1.02E-01 | SAAGSPTAAGAATSRGGGPGK +3 Methyl x 1 | R15 Methyl |
| LmjF.24.1470 | hypothetical protein, conserved | -3.13 | 8.25E-01 | GGGGGRGGEALLMNEsSSLQR +3 Methyl x 1 Phos x 1 | R6 Methyl:2H(3)13C(1), M13 Label:13C(1)2H(3) |
| LmjF.24.1470 | hypothetical protein, conserved | -1.19 | 1.00E+00 | TEFVDSTYRGGGGGRGGEALLMNESSSLQR +4 Methyl x 1 | R15 Methyl |
| LmjF.24.1470 | hypothetical protein, conserved | -0.34 | 8.04E-01 | GGGGGRGGEALLMNESSSLQR +3 Methyl x 1 | R6 Methyl:2H(3)13C(1), M13 Label:13C(1)2H(3) |
| LmjF.24.1580 | hypothetical protein, conserved | -0.03 | 2.78E-01 | HGAQPQQQGSRGGAVAAGAAAQAR +3 Methyl x 1 | R11 Methyl |
| LmjF.24.1580 | hypothetical protein, conserved | 0.12 | 3.48E-01 | HGAQPQQQGSRGGAVAAGAAAQAR +4 Methyl x 1 | R11 Methyl |
| LmjF.24.1880 | cyclin 11, putative | 0.34 | 1.10E+00 | SGVRGGGSGGGVR +2 Methyl x 1 | R4 Methyl:2H(3)13C(1) |
| LmjF.25.0290 | RNA-binding protein, putative | -0.12 | 4.32E-01 | GRGVGCMTNPPAPISDSGLAIPVPASR +3 Methyl x 1 | R2 Methyl:2H(3)13C(1), M7 Label:13C(1)2H(3) |
| LmjF.25.0290 | RNA-binding protein, putative | 10.97 | 1.00E+00 | SVARPPPPPPPIGRGR +3 Methyl x 2 | R14 Methyl, R16 Methyl |
| LmjF.25.0290 | RNA-binding protein, putative | 10.97 | 1.00E+00 | SVARPPPPPPPIGRGRGVGCMTNPPAPISDSGLAIPVPASR +5 Me | #N/A |
| LmjF.25.0290 | RNA-binding protein, putative | 1.25 | 1.00E+00 | SVARPPPPPPPIGRGRGVGCMTNPPAPISDSGLAIPVPASR +5 Me | #N/A |
| LmjF.25.0290 | RNA-binding protein, putative | 0.00 | 1.00E+00 | AEQYFVPSPtGPSIPLPR +2 Methyl x 1 Phos x 1 | R18 Methyl:2H(3)13C(1) |
| LmjF.25.0290 | RNA-binding protein, putative | -4.36 | 1.00E+00 | AEQYFVPSPTGPSIPLPR +2 Methyl x 1 | R18 Methyl:2H(3)13C(1) |
| LmjF.25.0290 | RNA-binding protein, putative | 0.13 | 1.03E+00 | AEQYFVPSPTGPSIPLPRGR +3 Methyl x 1 | R20 Methyl |
| LmjF.25.0540 | hypothetical protein SCD6.10 (SCD6) | 0.83 | 1.19E-01 | ADTETFGPEMVSSMRGFR +3 Methyl x 1 | R15 Methyl |
| LmjF.25.0540 | hypothetical protein SCD6.10 (SCD6) | -0.70 | 1.36E-01 | DPAIVEVHAPARGRGGR +3 Methyl x 3 | R14 Methyl:2H(3)13C(1), R17 Methyl:2H(3)13C(1) |
| LmjF.25.0540 | hypothetical protein SCD6.10 (SCD6) | 0.32 | 2.71E-01 | KADTETFGPEMVSSMR +3 Methyl x 1 | R16 Methyl |
| LmjF.25.0540 | hypothetical protein SCD6.10 (SCD6) | -0.09 | 3.88E-01 | DSsSPQARDPAIVEVHAPARGRGGR +4 Methyl x 2 Phos x 1 | R20 Methyl, R22 Methyl, R25 Methyl |
| LmjF.25.0540 | hypothetical protein SCD6.10 (SCD6) | -0.98 | 4.39E-01 | DSSSPQARDPAIVEVHAPARGRGGR +4 Methyl x 3 | R20 Methyl, R22 Methyl, R25 Methyl |
| LmjF.25.0540 | hypothetical protein SCD6.10 (SCD6) | 0.15 | 4.63E-01 | DSSSPQARDPAIVEVHAPARGRGGR +4 Methyl x 2 | R20 Methyl, R22 Methyl, R25 Methyl |
| LmjF.25.0540 | hypothetical protein SCD6.10 (SCD6) | 1.36 | 8.22E-01 | DPAIVEVHAPARGRGGR +3 Methyl x 2 | R14 Methyl:2H(3)13C(1), R17 Methyl:2H(3)13C(1) |
| LmjF.25.0540 | hypothetical protein SCD6.10 (SCD6) | -1.32 | 8.32E-01 | GGRAAAAAPAASATAASSAATR +3 Methyl x 1 | #N/A |
| LmjF.25.0540 | hypothetical protein SCD6.10 (SCD6) | -4.77 | 8.35E-01 | NHRGGYGGR +2 Methyl x 1 | R3 Methyl:2H(3)13C(1) |
| LmjF.25.0540 | hypothetical protein SCD6.10 (SCD6) | 1.13 | 8.43E-01 | KADTETFGPEMVSSMRGFR +4 Methyl x 1 | M11 Label:13C(1)2H(3), M15 Label:13C(1)2H(3), R16 Methyl:2H(3)13C(1) |
| LmjF.25.0540 | hypothetical protein SCD6.10 (SCD6) | 10.97 | 1.00E+00 | DPAIVEVHAPARGRGGRAAAAAPAASATAASSAATR +5 Methyl | R12 Methyl:2H(3)13C(1), R14 Methyl:2H(3)13C(1), R17 Methyl:2H(3)13C(1) |
| LmjF.25.0540 | hypothetical protein SCD6.10 (SCD6) | -0.49 | 1.03E+00 | SFYDEAQPYSRGGGGGR +3 Methyl x 1 | R11 Methyl |
| LmjF.25.0540 | hypothetical protein SCD6.10 (SCD6) | -1.18 | 1.07E+00 | GGYGGRGGSGNYR +2 Methyl x 1 | R6 Methyl |
| LmjF.25.0540 | hypothetical protein SCD6.10 (SCD6) | -1.04 | 1.09E+00 | GGYGGRGGSGNYR +3 Methyl x 1 | R6 Methyl |
| LmjF.25.1080 | hypothetical protein, conserved | -1.63 | 8.23E-01 | GGSRGGRGGAGSCER +3 Methyl x 2 | R4 Methyl, R7 Methyl |
| LmjF.25.1430 | Putative intraflagellar transport protein A1 | 0.00 | 1.00E+00 | RGGGGGSLtPAMsDDEDNGYENAPR +3 Methyl x 1 Phos x 1 | #N/A |
| LmjF.25.1540 | mitochondrial RNA binding complex 1 subunit, putative | 1.06 | 4.56E-01 | AIAAQSVGRGGVGLVESAAPPATSAFR +3 Methyl x 1 | R9 Methyl |
| LmjF.25.1540 | mitochondrial RNA binding complex 1 subunit, putative | 4.23 | 1.00E+00 | RGRGSYADAVSDGNMNR +3 Methyl x 2 | R1 Methyl, R3 Methyl |
| LmjF.25.1540 | mitochondrial RNA binding complex 1 subunit, putative | -0.38 | 1.00E+00 | AIAAQSVGRGGVGLVESAAPPATSAFR +4 Methyl x 1 | R9 Methyl |
| LmjF.25.1740 | mitochondrial RNA binding complex 1 subunit, putative | -0.11 | 3.17E-01 | HGYGGNNGSRGGGDGAPR +2 Methyl x 1 | R10 Methyl |
| LmjF.25.1740 | mitochondrial RNA binding complex 1 subunit, putative | -0.04 | 3.43E-01 | HGYGGNNGSRGGGDGAPR +3 Methyl x 1 | R10 Methyl |
| LmjF.25.1740 | mitochondrial RNA binding complex 1 subunit, putative | 1.10 | 4.03E-01 | RGGPMGSR +2 Methyl x 1 | R1 Methyl |
| LmjF.25.1740 | mitochondrial RNA binding complex 1 subunit, putative | -0.06 | 1.03E+00 | HGYGGNNGSRGGGDGAPR +4 Methyl x 1 | R10 Methyl |
| LmjF.25.1830 | Histone RNA hairpin-binding protein RNA-binding domain containing protein, putative | 0.53 | 1.00E+00 | MRGSNVSGSAR +2 Methyl x 1 | M1 Label:13C(1)2H(3), R2 Methyl:2H(3)13C(1) |
| LmjF.25.2000 | hypothetical protein, conserved | -0.50 | 8.26E-01 | GRGGCAGGAGTLHWQR +3 Methyl x 1 | R2 Methyl:2H(3)13C(1) |
| LmjF.25.2070 | Rieske [2Fe-2S] domain containing protein, putative | -0.47 | 6.28E-01 | ERPMSLPVDGGGRGLR +3 Methyl x 1 | M4 Label:13C(1)2H(3), R13 Methyl:2H(3)13C(1) |
| LmjF.26.0920 | hypothetical protein, conserved | -1.56 | 1.00E+00 | TGAALDCAPQPLNTLPSPYSDTELFRGR +3 Methyl x 1 | #N/A |
| LmjF.26.1110 | TROVE domain containing protein, putative | 0.47 | 8.54E-01 | RGGASVAVSATPAPQAATFVEAPR +3 Methyl x 1 | R1 Methyl |

| | | | | | |
|---|---|---|---|---|---|
| LmjF.26.1530 | RNA recognition motif. (a.k.a. RRM, RBD, or RNP domain), putative | -0.16 | 5.14E-01 | SHSNQYSQDSGVSSGQGAGMQRGMR +4 Methyl x 1 | M20 Label:13C(1)2H(3), M24 Label:13C(1)2H(3), R25 Methyl:2H(3)13C(1) |
| LmjF.26.1820 | hypothetical protein, conserved | -0.54 | 1.03E+00 | SSRGGTFLVPQSEDAR +2 Methyl x 1 | R3 Methyl |
| LmjF.26.1820 | hypothetical protein, conserved | 0.12 | 1.36E-01 | SSRGGTFLVPQSEDAR +3 Methyl x 1 | R3 Methyl |
| LmjF.26.2120 | hypothetical protein, conserved | 2.12 | 5.86E-01 | GASSPLLLPPPEFVGRGVGR +3 Methyl x 1 | R20 Methyl |
| LmjF.26.2170 | Cornifin (SPRR) family, putative | 0.52 | 8.74E-01 | SCRGGGSCFSAAHPTTSR +3 Methyl x 1 | R3 Methyl |
| LmjF.26.2380 | Fibronectin type III domain containing protein, putative | -1.52 | 4.11E-01 | RGGASTSSPAAATLTEMAANR +3 Methyl x 1 | R1 Methyl |
| LmjF.26.2420 | hypothetical protein, conserved | -0.12 | 5.30E-01 | RGGGTPLGR +2 Methyl x 1 | R1 Methyl, T5 Phospho |
| LmjF.27.0130 | WW domain/Zinc finger C-x8-C-x5-C-x3-H type (and similar), putative (ZFP3) | 0.87 | 7.48E-02 | TTHWSIPPTAYEPYYNNGRGRGAGYR +5 Methyl x 2 | #N/A |
| LmjF.27.0130 | WW domain/Zinc finger C-x8-C-x5-C-x3-H type (and similar), putative (ZFP3) | -7.63 | 1.00E+00 | TTHWSIPPTAYEPYYNNGRGR +4 Methyl x 1 | R19 Methyl:2H(3)13C(1), R21 Methyl:2H(3)13C(1) |
| LmjF.27.0270 | hypothetical protein, conserved | -1.16 | 1.03E+00 | TKVPGNNAQFDQPSNEVAsDLDDNHTALRGK +4 Methyl x 1 Ph | S19 Phospho, R29 Methyl |
| LmjF.27.0700 | hypothetical protein, conserved | -1.35 | 8.55E-01 | RGGGCGASAQGSGVDDMSQTASGDVR +3 Methyl x 1 | R1 Methyl:2H(3)13C(1), M17 Label:13C(1)2H(3) |
| LmjF.27.1300 | KH domain containing protein, putative | -2.35 | 1.17E-01 | GGGRGAGAPNGDDAAR +3 Methyl x 1 | R4 Methyl:2H(3)13C(1) |
| LmjF.27.1300 | KH domain containing protein, putative | -1.53 | 3.34E-01 | GGNDEGRGGYAPR +3 Methyl x 1 | R7 Methyl |
| LmjF.27.1300 | KH domain containing protein, putative | 0.01 | 6.82E-01 | RAGPPAGQPHPPR +3 Methyl x 1 | R1 Methyl |
| LmjF.27.1300 | KH domain containing protein, putative | 0.07 | 1.01E+00 | GGNDEGRGGYAPR +2 Methyl x 1 | R7 Methyl |
| LmjF.27.2170 | hypothetical protein, conserved | -0.62 | 1.21E-01 | SVQLSSSEVDSTRGGTGTR +3 Methyl x 1 | R13 Methyl:2H(3)13C(1) |
| LmjF.27.2170 | hypothetical protein, conserved | 0.38 | 8.93E-01 | RGGTGGSAGQRPVTPQSEIEINSR +4 Methyl x 1 | R11 Methyl:2H(3)13C(1) |
| LmjF.28.0825 | RNA binding protein rbp16, putative (RBP16) | 1.24 | 9.75E-01 | AENVTAPGGGKLPSGPRPPEGSPSGR +4 Methyl x 1 | R26 Methyl |
| LmjF.28.0825 | RNA binding protein rbp16, putative (RBP16) | 10.97 | 1.00E+00 | VVSWMSGRGFGFIEDNADKK +3 Methyl x 1 | R8 Methyl |
| LmjF.28.0825 | RNA binding protein rbp16, putative (RBP16) | 10.81 | 1.00E+00 | VVSWMSGRGFGFIEDNADKK +4 Methyl x 1 | R8 Methyl |
| LmjF.28.0825 | RNA binding protein rbp16, putative (RBP16) | 0.00 | 1.00E+00 | VVSWMSGRGFGFIEDNADK +3 Methyl x 1 | #N/A |
| LmjF.28.0825 | RNA binding protein rbp16, putative (RBP16) | -0.41 | 1.07E+00 | VVSWMSGR +2 Methyl x 1 | #N/A |
| LmjF.28.1060 | hypothetical protein, conserved | 0.00 | 1.00E+00 | RGQAATPTGNR +2 Methyl x 1 | R1 Methyl:2H(3)13C(1) |
| LmjF.28.1080 | hypothetical protein, conserved | -1.50 | 1.36E-01 | EVADTQDVEVPVAASRGGR +3 Methyl x 1 | R19 Methyl |
| LmjF.28.2780 | heat-shock protein hsp70, putative | -0.44 | 1.07E+00 | TTPSYVAFTDSER +2 Methyl x 1 | R13 Methyl |
| LmjF.29.0290 | zinc-finger of a C2HC-type, putative | -0.51 | 1.04E+00 | QGRGGISASVAPVSTSAAGR +2 Methyl x 1 | R3 Methyl |
| LmjF.29.0290 | zinc-finger of a C2HC-type, putative | -0.40 | 7.74E-01 | QGRGGISASVAPVSTSAAGR +3 Methyl x 1 | R3 Methyl |
| LmjF.29.0370 | protein kinase-like protein | -0.63 | 1.00E+00 | SSPLSNNLEANGTGRGGFSSPCPSPLK +3 Methyl x 1 | R15 Methyl:2H(3)13C(1) |
| LmjF.29.0370 | protein kinase-like protein | 0.00 | 1.00E+00 | sSPLSNNLEANGTGRGGFSSPCPSPLK +3 Methyl x 1 Phos x 1 | R15 Methyl:2H(3)13C(1) |
| LmjF.29.0680 | Triple RNA binding domain protein 3 (TRRM3) | -0.53 | 1.55E-01 | HAGGGNHGRGGAAAATR +4 Methyl x 1 | R9 Methyl |
| LmjF.29.0680 | Triple RNA binding domain protein 3 (TRRM3) | -0.43 | 3.16E-01 | HAGGGNHGRGGAAAATR +3 Methyl x 1 | R9 Methyl |
| LmjF.29.0680 | Triple RNA binding domain protein 3 (TRRM3) | 0.00 | 1.00E+00 | GGASASSASAFSSR +2 Methyl x 1 | #N/A |
| LmjF.29.0680 | Triple RNA binding domain protein 3 (TRRM3) | -0.15 | 1.00E+00 | RGGHDSSASHHDEEEHDsDEEAGLDSNAAK +3 Methyl x 1 Phos | R1 Methyl, S18 Phospho |
| LmjF.29.0680 | Triple RNA binding domain protein 3 (TRRM3) | 0.14 | 1.04E+00 | RGGHDSSASHHDEEEHDsDEEAGLDSNAAK +4 Methyl x 1 Phos | R1 Methyl, S18 Phospho |
| LmjF.29.0770 | hypothetical protein, conserved | -0.97 | 8.43E-01 | ATTDSVQGLLR +2 Methyl x 1 | R11 Methyl:2H(3)13C(1) |
| LmjF.29.1090 | ribosomal protein L1a, putative | -0.13 | 8.52E-01 | ISGGGTSTSGAGAFGNMCRGGR +3 Methyl x 1 | M17 Oxidation, R19 Methyl |
| LmjF.29.1090 | ribosomal protein L1a, putative | -1.94 | 1.04E+00 | ISGGGTSTSGAGAFGNMCR +2 Methyl x 1 | M17 Label:13C(1)2H(3), R19 Methyl:2H(3)13C(1) |
| LmjF.29.1110 | hypothetical protein, conserved | 10.97 | 1.00E+00 | AAGRGDAAGLLSQHASFGDVLR +4 Methyl x 1 | R4 Methyl:2H(3)13C(1) |
| LmjF.29.1230 | COPII coat assembly protein sec16, putative | -0.35 | 1.16E-01 | VDLTTTASRGR +3 Methyl x 1 | #N/A |
| LmjF.29.2330 | hypothetical protein, conserved | 0.03 | 4.99E-01 | VSsPTEGPGRGSVNLDDLFH +3 Methyl x 1 Phos x 1 | S3 Phospho, R10 Methyl |
| LmjF.30.0090 | hypothetical protein, conserved | 1.19 | 9.28E-01 | TRIEGFGRGGTQVSR +3 Methyl x 1 | R8 Methyl |
| LmjF.30.0260 | mitochondrial RNA binding protein 1, putative | -3.64 | 6.16E-01 | QQRGGAGCVGAASNSHAAEGFDVAYPVR +4 Methyl x 1 | #N/A |
| LmjF.30.0320 | Sad1 / UNC-like C-terminal, putative | 10.97 | 1.00E+00 | YYNIGRGSGGVALGGGNGPPLWPR +3 Methyl x 1 | R6 Methyl:2H(3)13C(1) |
| LmjF.30.0760 | hypothetical protein, conserved | -2.97 | 3.18E-01 | RGGSGGDVAVAGGNEGR +2 Methyl x 1 | R1 Methyl:2H(3)13C(1) |
| LmjF.30.0760 | hypothetical protein, conserved | 4.25 | 4.60E-01 | RGGSGGDVAVAGGNEGRGR +3 Methyl x 2 | R1 Methyl:2H(3)13C(1), R17 Methyl:2H(3)13C(1) |
| LmjF.30.0760 | hypothetical protein, conserved | -0.27 | 1.02E+00 | GRGGGGHHEDAMEK +3 Methyl x 1 | R2 Methyl |
| LmjF.30.0760 | hypothetical protein, conserved | 0.00 | 1.00E+00 | GRGGGGHHEDAMEK +2 Methyl x 1 | R2 Methyl |
| LmjF.30.0760 | hypothetical protein, conserved | 0.00 | 1.00E+00 | RGGPPTR +2 Methyl x 1 | #N/A |
| LmjF.30.0780 | mitochondrial oligo_U binding protein TBRGG1, putative | -0.39 | 1.02E+00 | GGYEGRGGGPR +2 Methyl x 1 | R6 Methyl:2H(3)13C(1) |
| LmjF.30.0900 | hypothetical protein, conserved | -1.05 | 1.23E-01 | RGNGIIRPGHSSAPSPAQASAVSVDQPR +4 Methyl x 1 | #N/A |
| LmjF.30.1810 | Zeta toxin, putative | -0.96 | 8.59E-01 | AGTGSSSSTAGSGRGGNAGGGNYAHTSK +4 Methyl x 1 | R14 Methyl:2H(3)13C(1) |
| LmjF.30.1810 | Zeta toxin, putative | 0.00 | 1.00E+00 | AGTGSSSSTAGSGRGGNAGGGNYAHTSK +3 Methyl x 1 | R14 Methyl:2H(3)13C(1) |
| LmjF.30.2710 | hypothetical protein, conserved | -0.32 | 8.55E-01 | SSSGAGGRGASGGPGGGDSGQR +3 Methyl x 1 | S2 Phospho, R8 Methyl |
| LmjF.30.2710 | hypothetical protein, conserved | 0.00 | 1.00E+00 | SsSGAGGRGASGGPGGGDSGQR +3 Methyl x 1 Phos x 1 | S2 Phospho, R8 Methyl |
| LmjF.31.0080 | hypothetical protein, conserved (ZC3H34) | -1.99 | 9.73E-01 | GRGGWYYPSAGSGGYNNMR +3 Methyl x 1 | #N/A |
| LmjF.31.0080 | hypothetical protein, conserved (ZC3H34) | 0.00 | 1.00E+00 | HMDGIPPHAYNQPPVNYYQPPQVYPMGNPMYNVPQGYPPAM | R45 Methyl |
| LmjF.31.0560 | mevalonate kinase, putative | -0.68 | 8.51E-01 | LSGTGRGGIAVALAASSDQR +3 Methyl x 1 | R6 Methyl:2H(3)13C(1) |
| LmjF.31.1050 | hypothetical protein, unknown function | -2.19 | 9.29E-01 | SHAVPYEGTVVHPSSRGR +4 Methyl x 1 | #N/A |
| LmjF.31.1260 | Microtubule-binding stalk of dynein motor, putative | -0.69 | 4.76E-01 | VSPAPSPRGGSAK +2 Methyl x 1 | R8 Methyl |
| LmjF.31.1380 | hypothetical protein, unknown function | -0.38 | 6.28E-01 | TARGGAAACVLQR +3 Methyl x 1 | R3 Methyl:2H(3)13C(1) |
| LmjF.31.1750 | nucleosome assembly protein-like protein | -2.78 | 1.02E+00 | GGRGGAASQQPPQQECK +3 Methyl x 1 | R3 Methyl:2H(3)13C(1) |
| LmjF.31.1750 | nucleosome assembly protein-like protein | -1.92 | 4.37E-01 | QRGGGATPGGR +2 Methyl x 1 | R2 Methyl |
| LmjF.31.2360 | hypothetical protein, conserved | 0.95 | 6.36E-01 | LQRGGYGQALQGR +3 Methyl x 1 | R3 Methyl |
| LmjF.31.2590 | TerD domain containing protein, putative | 0.00 | 1.00E+00 | RGGRGEVTVQTAR +3 Methyl x 1 | R1 Methyl |
| LmjF.31.3030 | hypothetical protein, unknown function | 0.34 | 1.07E+00 | LATPPYEGVQLPGAsAAEER +4 Methyl x 1 Phos x 1 | #N/A |
| LmjF.32.0400 | ATP-dependent RNA helicase HEL67 (DDX3) | -0.16 | 5.30E-01 | NGGGGGFRGGFGGFGGGAR +3 Methyl x 1 | R8 Methyl:2H(3)13C(1) |
| LmjF.32.0400 | ATP-dependent RNA helicase HEL67 (DDX3) | 0.02 | 8.53E-01 | QYGGGNRGFNDAQGTYGGGFNR +3 Methyl x 1 | R7 Methyl |
| LmjF.32.0400 | ATP-dependent RNA helicase HEL67 (DDX3) | 0.32 | 1.00E+00 | QYGGGNRGFNDAQGTYGGGFNR +2 Methyl x 1 | R7 Methyl |
| LmjF.32.0400 | ATP-dependent RNA helicase HEL67 (DDX3) | -0.74 | 1.03E+00 | SGGFGGGFGGGYGGGYGGRGGYR +3 Methyl x 1 | R19 Methyl |
| LmjF.32.0400 | ATP-dependent RNA helicase HEL67 (DDX3) | 0.07 | 1.05E+00 | NGGGGGFRGGFGGFGGGAR +2 Methyl x 1 | R8 Methyl:2H(3)13C(1) |
| LmjF.32.0620 | hypothetical protein, conserved | -0.31 | 2.77E-01 | GPSFVERGAGAR +3 Methyl x 1 | #N/A |
| LmjF.32.0620 | hypothetical protein, conserved | 6.24 | 1.00E+00 | GGMGRAGSGR +2 Methyl x 1 | R5 Methyl |
| LmjF.32.0840 | hypothetical protein, conserved (DRBD18) | 0.51 | 1.00E+00 | GRGGRGGCGLDTMGMQAGYMPFPQVGGMAR +4 Methyl x 2 | R2 Methyl:2H(3)13C(1), M13 Label:13C(1)2H(3), M15 Label:13C(1)2H(3), M20 Label:13C(1)2H(3), M28 Label:13C(1)2H(3) |
| LmjF.32.0840 | hypothetical protein, conserved (DRBD18) | -7.25 | 1.00E+00 | GGRGGCGLDTMGMQAGYMPFPQVGGMAR +3 Methyl x 1 | R3 Methyl |
| LmjF.32.0850 | polypyrimidine tract-binding protein, putative (DRBD4) | 0.66 | 1.18E-01 | GGIGGRGSGTGLGAATGFSPFEPSGTTPARGGGASVGR +4 Meth | R6 Methyl:2H(3)13C(1), R30 Methyl:2H(3)13C(1), R38 Methyl:2H(3)13C(1) |
| LmjF.32.0850 | polypyrimidine tract-binding protein, putative (DRBD4) | -0.42 | 8.29E-01 | GGIGGRGSGTGLGAATGFSPFEPSGTTPAR +3 Methyl x 1 | R6 Methyl:2H(3)13C(1) |
| LmjF.32.0880 | 60S ribosomal protein L18a, putative | 0.74 | 7.75E-01 | HYQVVGR +2 Methyl x 1 | #N/A |
| LmjF.32.1130 | hypothetical protein, conserved | -0.66 | 9.70E-02 | QSASPRGGGSAVAR +3 Methyl x 1 | R6 Methyl |
| LmjF.32.1650 | hypothetical protein, conserved | -0.89 | 2.88E-01 | SIRAGGPISGAGK +3 Methyl x 1 | R3 Methyl:2H(3)13C(1) |
| LmjF.32.1800 | hypothetical protein, conserved | -0.71 | 7.77E-01 | GGGAGRGGQGIR +2 Methyl x 1 | R6 Methyl:2H(3)13C(1) |
| LmjF.32.1970 | ribosome biogenesis protein TSR3, putative | -0.18 | 8.36E-01 | GRGGGSGGVCSTSGFHNNEPPAHPCSVPLAMWDFEQCDPNACS | #N/A |
| LmjF.32.2130 | RNA recognition motif. (a.k.a. RRM, RBD, or RNP domain), putative | 0.01 | 2.93E-01 | EPYPSLNSSRPPYGPGMGRGGGMR +4 Methyl x 1 | #N/A |
| LmjF.32.2490 | hypothetical protein, conserved | -0.14 | 8.61E-01 | GYQYGAGRGAK +2 Methyl x 1 | R8 Methyl |
| LmjF.32.2520 | hypothetical protein, unknown function | 0.74 | 8.24E-01 | GHPSPVGERGGR +3 Methyl x 1 | R9 Methyl |
| LmjF.32.2520 | hypothetical protein, unknown function | 1.31 | 1.00E+00 | SAPPRGGYGAAPMSLSPAPDPALMSK +3 Methyl x 1 | #N/A |
| LmjF.32.2580 | hypothetical protein, conserved | -0.82 | 1.05E+00 | MHEEQDSGGGRGGGR +3 Methyl x 1 | M1 Label:13C(1)2H(3), R11 Methyl:2H(3)13C(1) |
| LmjF.32.2830 | hypothetical protein, conserved | 0.71 | 8.45E-01 | QNVTGSGRGsSVAAQGDGADVGEGSTAPR +3 Methyl x 1 Phos | R8 Methyl:2H(3)13C(1) |
| LmjF.32.2830 | hypothetical protein, conserved | 0.00 | 1.00E+00 | QNVTGSGRGSSVAAQGDGADVGEGSTAPR +3 Methyl x 1 | R8 Methyl:2H(3)13C(1) |
| LmjF.32.3090 | hypothetical protein, conserved | -0.31 | 1.06E+00 | SPALVTSPSSSASRGGR +3 Methyl x 1 | R14 Methyl |
| LmjF.32.3090 | hypothetical protein, conserved | 0.00 | 1.00E+00 | AELPASGGTSAAPSLGAARGGK +3 Methyl x 1 | R19 Methyl:2H(3)13C(1) |
| LmjF.32.3690 | hypothetical protein, conserved | 6.81 | 8.57E-01 | RGGEVSGAGAAEGR +2 Methyl x 1 | R1 Methyl:2H(3)13C(1) |
| LmjF.33.0260 | RGG-containing protein 2, putative (RGG2) | -0.18 | 4.53E-01 | DSSDGWGGGRGGGHSGWGGDGGWKDAPTGDR +4 Methyl x | R10 Methyl:2H(3)13C(1) |
| LmjF.33.0260 | RGG-containing protein 2, putative (RGG2) | -1.16 | 8.61E-01 | RGRGGSGGGVWGQPAAADEDAWNAAPPSFQPPVR +4 Methyl | #N/A |
| LmjF.33.0260 | RGG-containing protein 2, putative (RGG2) | 0.21 | 9.74E-01 | DSSDGWGGGRGGGHSGWGGDGGWKDAPTGDR +3 Methyl x | R10 Methyl:2H(3)13C(1) |
| LmjF.33.0260 | RGG-containing protein 2, putative (RGG2) | -2.14 | 1.00E+00 | GRGGSGGGVWGQPAAADEDAWNAAPPSFQPPVR +3 Methyl x | R2 Methyl:2H(3)13C(1) |
| LmjF.33.0940 | hypothetical protein, conserved | -1.17 | 1.00E+00 | QASGRGGGNAVAMSLLTGASATNLR +3 Methyl x 1 | R5 Methyl:2H(3)13C(1), M13 Label:13C(1)2H(3) |
| LmjF.33.1080 | YEATS family, putative | 0.14 | 8.08E-02 | VGAQEVRGAGPSPSPSR +3 Methyl x 1 | #N/A |
| LmjF.33.1150 | pumilio protein 6, putative (PUF6) | -0.58 | 6.84E-01 | FARGGGIGNINPSGR +2 Methyl x 1 | R3 Methyl:2H(3)13C(1) |
| LmjF.33.1150 | pumilio protein 6, putative (PUF6) | 0.07 | 8.55E-01 | GGRGAGGMGSMESGSGPNSHR +4 Methyl x 1 | R3 Methyl |
| LmjF.33.1150 | pumilio protein 6, putative (PUF6) | -0.59 | 8.71E-01 | FARGGGIGNINPSGR +3 Methyl x 1 | R3 Methyl:2H(3)13C(1) |
| LmjF.33.1150 | pumilio protein 6, putative (PUF6) | -0.25 | 8.77E-01 | GGRGAGGMGSMESGSGPNSHR +3 Methyl x 1 | R3 Methyl |
| LmjF.33.1150 | pumilio protein 6, putative (PUF6) | -0.29 | 9.75E-01 | RYNNNNSAGGGVGGGRVGR +3 Methyl x 1 | R16 Methyl |
| LmjF.33.1150 | pumilio protein 6, putative (PUF6) | 0.29 | 1.00E+00 | NNYRGGRGAGGMGSMESGSGPNSHR +4 Methyl x 2 | R4 Methyl, R7 Methyl |

| | | | | | |
|---|---|---|---|---|---|
| LmjF.33.1220 | hypothetical protein, conserved | -1.34 | 8.07E-01 | EVVTTDNPDWASPLIARGGR +3 Methyl x 1 | R20 Methyl |
| LmjF.33.2130 | hypothetical protein, conserved | 0.76 | 1.00E+00 | ESSGSGTRGHGGAGGGAAASGGGGHR +3 Methyl x 1 | R8 Methyl:2H(3)13C(1) |
| LmjF.33.2130 | hypothetical protein, conserved | -0.07 | 1.04E+00 | ESSGSGTRGHGGAGGGAAASGGGGHR +4 Methyl x 1 | R8 Methyl:2H(3)13C(1) |
| LmjF.33.2210 | hypothetical protein, conserved | -1.36 | 1.04E+00 | HLRGGAAAAR +3 Methyl x 1 | R3 Methyl:2H(3)13C(1) |
| LmjF.33.2360 | ATP synthase regulation protein NCA2, putative | 0.00 | 1.00E+00 | GGGVWGCDFNANQNSMFWSPIAAGVGRGR +3 Methyl x 1 | R29 Methyl |
| LmjF.33.2450 | hypothetical protein, conserved | -0.51 | 8.50E-01 | AGSAVLGGGGPNAHAPPARGGGGSGAGGR +3 Methyl x 1 | R29 Methyl |
| LmjF.33.2640 | hypothetical protein, conserved | -0.44 | 6.45E-01 | QAVCSECAGRGYEHWVPR +4 Methyl x 1 | R10 Methyl |
| LmjF.34.1110 | Shwachman-Bodian-Diamond syndrome (SBDS) protein/SBDS protein C-terminal domain conta | 0.82 | 1.00E+00 | LGLDPSHDLQDDsDDDGGRGRGSR +4 Methyl x 1 Phos x 1 | S13 Phospho, R21 Methyl |
| LmjF.34.1260 | mitochondrial DNA polymerase I protein A, putative | 0.33 | 8.34E-01 | KRGGGDAILPGALGPSVCATPAPR +4 Methyl x 1 | R2 Methyl |
| LmjF.34.1470 | EAP30/Vps36 family, putative | -1.95 | 8.58E-01 | VTTAGARGGSPAK +2 Methyl x 1 | R7 Methyl:2H(3)13C(1) |
| LmjF.34.2430 | DNAj-like protein | -0.38 | 4.75E-01 | AGGFNFSSTYSGGNANDYYRGR +3 Methyl x 1 | R20 Methyl |
| LmjF.34.2473 | hypothetical protein, conserved | -0.15 | 1.80E-01 | QSVAGRGGSGGGGYGGAFY +2 Methyl x 1 | R6 Methyl:2H(3)13C(1) |
| LmjF.34.3815 | ribosomal protein L14, putative | 0.31 | 1.01E+00 | FKGRGGGGEVSR +3 Methyl x 1 | R4 Methyl:2H(3)13C(1) |
| LmjF.34.4290 | nucleolar protein family a, putative | 2.10 | 7.26E-01 | GGRGGGGFGGGR +2 Methyl x 1 | R3 Methyl |
| LmjF.34.4290 | nucleolar protein family a, putative | -5.12 | 8.20E-01 | GGRGGGGFGGGR +3 Methyl x 1 | R3 Methyl |
| LmjF.34.4290 | nucleolar protein family a, putative | -0.65 | 1.00E+00 | GGGRGGHMSEPDPPENVEEVGTFMNAAEGELVYK +3 Methyl x | R4 Methyl |
| LmjF.34.4290 | nucleolar protein family a, putative | -4.18 | 1.00E+00 | GGGRGGHMSEPDPPENVEEVGTFMNAAEGELVYK +4 Methyl x | R4 Methyl |
| LmjF.35.0610 | hypothetical protein, conserved | 0.00 | 1.00E+00 | ANGPAPSGSRGSK +2 Methyl x 1 | R10 Methyl:2H(3)13C(1) |
| LmjF.35.0630 | Nucleoporin NUP65, putative | -0.90 | 8.30E-01 | RGGLILESPQANVPDSADATESILR +3 Methyl x 1 | R1 Methyl:2H(3)13C(1) |
| LmjF.35.0630 | Nucleoporin NUP65, putative | 0.00 | 1.00E+00 | RRGGLILESPQANVPDSADATESILR +4 Methyl x 1 | #N/A |
| LmjF.35.0740 | hypothetical protein, conserved | 0.01 | 9.26E-01 | ARGGGASGSTSGLGTAK +2 Methyl x 1 | R2 Methyl:2H(3)13C(1) |
| LmjF.35.0740 | hypothetical protein, conserved | 0.43 | 1.07E+00 | ARGGGASGSTSGLGTAK +3 Methyl x 1 | R2 Methyl:2H(3)13C(1) |
| LmjF.35.1510 | NLI interacting factor-like phosphatase/Zinc finger C-x8-C-x5-C-x3-H type (and similar), putative | 0.26 | 7.31E-02 | RANNSNDHQHQRGGGAR +3 Methyl x 1 | #N/A |
| LmjF.35.1510 | NLI interacting factor-like phosphatase/Zinc finger C-x8-C-x5-C-x3-H type (and similar), putativ | 2.24 | 4.74E-01 | RANNSNDHQHQRGGGAR +4 Methyl x 1 | #N/A |
| LmjF.35.1620 | Concanavalin A-like lectin/glucanases superfamily, putative | -0.37 | 8.46E-01 | NNGSSSNRANQHGVASStTIGASSAAR +3 Methyl x 2 Phos x 1 | #N/A |
| LmjF.35.2090 | kinesin, putative | 0.44 | 5.68E-02 | RGGGPASAAQDAQLSEWEMPFQGTVFR +3 Methyl x 1 | R1 Methyl |
| LmjF.35.2200 | RNA-binding protein, putative | 2.45 | 8.29E-01 | HNMAGSAYGR +2 Methyl x 1 | M3 Oxidation, R10 Methyl |
| LmjF.35.2200 | RNA-binding protein, putative | 10.97 | 1.00E+00 | HNMAGSAYGRGAPYQVGAAESTEMNGTEPLPKPR +4 Methyl x | M3 Oxidation, R10 Methyl, M24 Oxidation |
| LmjF.35.2200 | RNA-binding protein, putative | 0.00 | 1.00E+00 | HNMAGSAYGRGAPYQVGAAEsTEMNGTEPLPKPR +3 Methyl x | M3 Oxidation, R10 Methyl, M24 Oxidation |
| LmjF.35.3100 | ATP-dependent RNA helicase, putative (DED1) | 0.03 | 8.06E-02 | SGGPGGRGGGGGGGSWGDSRPSAPSGGGR +4 Methyl x 1 | R7 Methyl |
| LmjF.35.3100 | ATP-dependent RNA helicase, putative (DED1) | 0.14 | 6.16E-01 | SGGPGGRGGGGGGGSWGDSRPSAPSGGGR +3 Methyl x 1 | R7 Methyl |
| LmjF.35.3100 | ATP-dependent RNA helicase, putative (DED1) | 0.22 | 8.48E-01 | RRGGGGGGGGGHR +3 Methyl x 3 | #N/A |
| LmjF.35.3100 | ATP-dependent RNA helicase, putative (DED1) | 0.00 | 1.00E+00 | RYYDDEDDYGEGGDRAGGTGNEDDDYEDGGYDAYR +4 Methyl | R1 Methyl |
| LmjF.35.3100 | ATP-dependent RNA helicase, putative (DED1) | -0.36 | 1.03E+00 | RGGGVDDGGF +2 Methyl x 1 | R1 Methyl:2H(3)13C(1) |
| LmjF.35.3310 | Leucine Rich repeats (2 copies), putative | -1.13 | 1.00E+00 | KATSSVSGFGGGNVYNGRGGSGASGNPTIHK +4 Methyl x 1 | #N/A |
| LmjF.35.3310 | Leucine Rich repeats (2 copies), putative | -0.97 | 8.21E-01 | ATSSVSGFGGGNVYNGRGGSGASGNPTIHK +3 Methyl x 1 | R17 Methyl:2H(3)13C(1) |
| LmjF.35.3310 | Leucine Rich repeats (2 copies), putative | 0.63 | 2.82E-01 | ATSSVSGFGGGNVYNGRGGSGASGNPTIHK +4 Methyl x 1 | R17 Methyl:2H(3)13C(1) |
| LmjF.35.3310 | Leucine Rich repeats (2 copies), putative | -0.13 | 2.14E-01 | KATsSVSGFGGGNVYNGRGGSGASGNPTIHK +4 Methyl x 1 Pho | #N/A |
| LmjF.35.3980 | 4E-interacting protein, putative | 0.48 | 2.09E-01 | RGGGGGGGNGGRDDSSNSNSVR +4 Methyl x 1 | R1 Methyl, S15 Phospho |
| LmjF.35.3980 | 4E-interacting protein, putative | 1.47 | 2.15E-01 | GGHNSGMNGGGSSSNHPGSSSTPVYGSGGGRGGDNR +4 Meth | M7 Oxidation, R31 Methyl |
| LmjF.35.3980 | 4E-interacting protein, putative | -0.23 | 8.05E-01 | RGGGGGGGNGGRDDSSNSNSVR +3 Methyl x 1 | R1 Methyl, S15 Phospho |
| LmjF.35.3980 | 4E-interacting protein, putative | -0.11 | 8.42E-01 | QGRGVTGNGMSTVPPSPPQFQQPPPPR +3 Methyl x 1 | R3 Methyl |
| LmjF.35.3980 | 4E-interacting protein, putative | 10.97 | 1.00E+00 | GGHNSGMNGGGSSSNHPGSSSTPVYGSGGGRGGDNRR +5 Met | M7 Label:13C(1)2H(3), R37 Methyl:2H(3)13C(1) |
| LmjF.35.4125 | hypothetical protein, conserved | -0.26 | 3.75E-01 | SARGGASNGNHSFSASCTATR +3 Methyl x 1 | R3 Methyl |
| LMJF.35.4140 | hypothetical protein, unknown function | 2.02 | 1.05E+00 | ERGGFSAR +2 Methyl x 1 | R2 Methyl:2H(3)13C(1) |
| LmjF.35.4380 | hypothetical protein, conserved | 0.00 | 1.00E+00 | GGRGGAPQIGLSTPSLNTR +3 Methyl x 1 | #N/A |
| LmjF.35.4950 | zinc finger protein family member, putative | -0.42 | 3.85E-01 | SPPATSASTTSAPPPALRGLGR +3 Methyl x 1 | R22 Methyl:2H(3)13C(1) |
| LmjF.35.4950 | zinc finger protein family member, putative | 0.27 | 4.72E-01 | LSSMERGGMTGVPQHNLLPPR +4 Methyl x 1 | R6 Methyl |
| LmjF.35.4950 | zinc finger protein family member, putative | 0.65 | 8.14E-01 | GAQQHHGYRGVGENGSSR +3 Methyl x 1 | #N/A |
| LmjF.35.4950 | zinc finger protein family member, putative | -4.78 | 8.52E-01 | SPPATSASTTSAPPPALR +2 Methyl x 1 | R18 Methyl:2H(3)13C(1) |
| LmjF.35.4950 | zinc finger protein family member, putative | -1.07 | 8.56E-01 | LSSMERGGMTGVPQHNLLPPR +4 Methyl x 2 | R6 Methyl |
| LmjF.35.4950 | zinc finger protein family member, putative | 0.70 | 1.00E+00 | GTEGVGNSYYRGAQQHHGYR +4 Methyl x 1 | #N/A |
| LmjF.35.4950 | zinc finger protein family member, putative | -0.65 | 1.00E+00 | GATVTGGVHIDSSVGPRGGAAPMSGMGAGAASR +4 Methyl x 1 | R17 Methyl |
| LmjF.35.4950 | zinc finger protein family member, putative | -0.25 | 1.03E+00 | LSSMERGGMTGVPQHNLLPPRAsPASSQPR +4 Methyl x 1 Phos | #N/A |
| LmjF.35.4950 | zinc finger protein family member, putative | -0.05 | 1.03E+00 | LSSMERGGMTGVPQHNLLPPR +3 Methyl x 1 | R6 Methyl |
| LmjF.35.5040 | polyadenylate-binding protein 1 (PABP1) | 10.97 | 1.00E+00 | QQLGGRAQGHPMPMPSPQQPQAPAQPQGFATPSAVGFVQATP | R6 Methyl |
| LmjF.36.0450 | hypothetical protein, conserved | 0.53 | 9.34E-01 | STFSAGIGEGGIAGGRGGAR +3 Methyl x 1 | R16 Methyl |
| LmjF.36.0490 | zinc-finger of a C2HC-type, putative | -0.41 | 4.17E-01 | GPARGGLFGATPSGYGDGAGMSAR +3 Methyl x 1 | R4 Methyl:2H(3)13C(1), M21 Label:13C(1)2H(3)+Oxidation |
| LmjF.36.0490 | zinc-finger of a C2HC-type, putative | -0.56 | 6.04E-01 | GPARGGLFGAtPSGYGDGAGMSAR +3 Methyl x 1 Phos x 1 | R4 Methyl:2H(3)13C(1), M21 Label:13C(1)2H(3)+Oxidation |
| LmjF.36.0490 | zinc-finger of a C2HC-type, putative | -0.56 | 8.08E-01 | FSSRGGGGGMGGGGGGGR +3 Methyl x 1 | R4 Methyl:2H(3)13C(1), M10 Label:13C(1)2H(3)+Oxidation |
| LmjF.36.1240 | hypothetical protein, conserved | -1.38 | 1.07E+00 | VSPPVSPTPPILPRGR +3 Methyl x 1 | #N/A |
| LmjF.36.1640 | universal minicircle sequence binding protein, putative | 0.73 | 1.22E-01 | CGQEGHLSRDCPSSQGGSRGGYGQK +3 Methyl x 1 | R19 Methyl |
| LmjF.36.1640 | universal minicircle sequence binding protein, putative | -0.15 | 8.55E-01 | DCPSSQGGSRGGYGQKR +4 Methyl x 1 | R10 Methyl:2H(3)13C(1) |
| LmjF.36.1640 | universal minicircle sequence binding protein, putative | -0.25 | 8.81E-01 | DCPSSQGGSRGGYGQKR +3 Methyl x 1 | R10 Methyl:2H(3)13C(1) |
| LmjF.36.1640 | universal minicircle sequence binding protein, putative | -0.28 | 9.26E-01 | DCPSSQGGSRGGYGQK +2 Methyl x 1 | R10 Methyl:2H(3)13C(1) |
| LmjF.36.1640 | universal minicircle sequence binding protein, putative | -0.26 | 1.01E+00 | DCPSSQGGSRGGYGQK +3 Methyl x 1 | R10 Methyl:2H(3)13C(1) |
| LmjF.36.1925 | 60S ribosomal protein L37a | -0.19 | 1.08E+00 | TVAGGAYTLSTPNNSTVR +2 Methyl x 1 | #N/A |
| LmjF.36.2130 | ATP-dependent RNA helicase DBP2A, putative | -0.68 | 2.91E-01 | NYSPFSGFSTTSRGGGGSGAHR +3 Methyl x 1 | R13 Methyl:2H(3)13C(1) |
| LmjF.36.2130 | ATP-dependent RNA helicase DBP2A, putative | -0.08 | 8.50E-01 | NYSPFSGFSTTSR +2 Methyl x 1 | #N/A |
| LmjF.36.2130 | ATP-dependent RNA helicase DBP2A, putative | 0.52 | 1.06E+00 | NYSPFSGFSTTSRGGGGSGAHR +4 Methyl x 1 | R13 Methyl:2H(3)13C(1) |
| LmjF.36.2580 | related to multifunctional cyclin-dependent kinase pho85-like protein | 0.67 | 8.90E-01 | VMRGSSSSSQNDMCGGSSAR +3 Methyl x 1 | R3 Methyl |
| LmjF.36.2670 | hypothetical protein, conserved | -2.47 | 1.04E+00 | IATRGGGGQPEGVVR +3 Methyl x 1 | R4 Methyl |
| LmjF.36.3200 | DNA topoisomerase III, putative | 0.00 | 1.00E+00 | GGRGGDAAAANTPSGGVDPVCGCGTPAK +3 Methyl x 1 | #N/A |
| LmjF.36.3280 | Enoyl-CoA hydratase/isomerase family/2-enoyl-CoA Hydratase C-terminal region, putative | 0.19 | 1.00E+00 | WFIQPGHNPNPSGLLR +3 Methyl x 1 | #N/A |
| LmjF.36.3280 | Enoyl-CoA hydratase/isomerase family/2-enoyl-CoA Hydratase C-terminal region, putative | 0.27 | 1.00E+00 | WFIQPGHNPNPSGLLRGR +4 Methyl x 1 | #N/A |
| LmjF.36.4330 | predicted C2 domain protein | 10.97 | 1.00E+00 | YECDHGVGHTEAPPAGRGGSHGR +3 Methyl x 1 | #N/A |
| LmjF.36.4590 | PHD-like zinc-binding domain containing protein, putative | -0.91 | 1.05E+00 | NSGGGTTDSAGALSSSPTARGGAGVGGGHAR +4 Methyl x 1 | R31 Methyl:2H(3)13C(1) |
| LmjF.36.5100 | hypothetical protein, conserved (PUF11) | -0.86 | 2.61E-01 | GGYQPTPPQQQAQLQQGYASRGGYYQGGGAYGGAGIQQGYPG. | R21 Methyl |
| LmjF.36.5100 | hypothetical protein, conserved (PUF11) | -0.85 | 4.72E-01 | GYPPQDGMQGYPQQAGR +2 Methyl x 1 | M8 Label:13C(1)2H(3), R17 Methyl:2H(3)13C(1) |
| LmjF.36.5100 | hypothetical protein, conserved (PUF11) | -0.26 | 5.02E-01 | GYSQAYNQSYAAR +2 Methyl x 1 | R13 Methyl |
| LmjF.36.5100 | hypothetical protein, conserved (PUF11) | -0.83 | 8.03E-01 | GYSQAYNQSYAARGGYQPTPPQQQAQLQQGYASR +4 Methyl x | R13 Methyl:2H(3)13C(1) |
| LmjF.36.5100 | hypothetical protein, conserved (PUF11) | 0.46 | 8.50E-01 | GYSQAYNQSYAARGGYQPTPPQQQAQLQQGYASR +3 Methyl x | R13 Methyl:2H(3)13C(1) |
| LmjF.36.5100 | hypothetical protein, conserved (PUF11) | -0.39 | 1.00E+00 | GGYQPTPPQQQAQLQQGYASRGGYYQGGGAYGGAGIQQGYPG. | R21 Methyl |
| LmjF.36.5100 | hypothetical protein, conserved (PUF11) | -1.32 | 1.02E+00 | GGYGPGYGPGYYPVQQQQGYYNQGYNQNPNYTPAAR +3 Meth | R36 Methyl:2H(3)13C(1) |
| LmjF.36.5850 | flagellum targeting protein kharon1, putative | -3.12 | 1.00E+00 | TRGGsVNGATAPFR +2 Methyl x 2 Phos x 1 | R2 Methyl |
| LmjF.36.5850 | flagellum targeting protein kharon1, putative | -2.21 | 8.03E-01 | TRGGSVNGATAPFR +3 Methyl x 2 | R2 Methyl |
| LmjF.36.5850 | flagellum targeting protein kharon1, putative | -1.72 | 8.15E-01 | TRGGSVNGATAPFR +2 Methyl x 1 | R2 Methyl |
| LmjF.36.5850 | flagellum targeting protein kharon1, putative | -0.84 | 8.47E-01 | tRGGsVNGATAPFR +2 Methyl x 1 Phos x 1 | R2 Methyl |
| LmjF.36.5850 | flagellum targeting protein kharon1, putative | -0.72 | 8.61E-01 | TRGGsVNGATAPFR +3 Methyl x 1 Phos x 1 | R2 Methyl |
| LmjF.36.5850 | flagellum targeting protein kharon1, putative | -0.43 | 2.80E-01 | TRGGSVNGATAPFR +3 Methyl x 1 | R2 Methyl |
| LmjF.36.5850 | flagellum targeting protein kharon1, putative | -0.35 | 5.67E-01 | ASLHNLMYASGDAYGAVASRGQGASR +4 Methyl x 1 | R20 Methyl |
| LmjF.36.6030 | hypothetical protein, conserved | -3.27 | 8.49E-01 | FRGMR +2 Methyl x 1 | #N/A |
| LmjF.36.6060 | eukaryotic translation initiation factor 4 gamma 4 | -0.09 | 1.18E-01 | GPIIGSSNAGGNRGGTRPPSMADR +4 Methyl x 1 | #N/A |
| LmjF.36.6060 | eukaryotic translation initiation factor 4 gamma 4 | 0.23 | 1.09E+00 | YGRGGGPGGLR +2 Methyl x 1 | R3 Methyl:2H(3)13C(1) |
| LmjF.36.6525 | hypothetical protein, conserved | 0.13 | 1.00E+00 | SRGGAPHSVSFASHSDR +4 Methyl x 1 | R2 Methyl |
| LmjF.36.6980 | eukaryotic translation initiation factor 3 subunit c | -2.11 | 4.70E-01 | GRGGMAGGRGGAGVR +3 Methyl x 2 | R2 Methyl, R9 Methyl |
| LmjF.36.6980 | eukaryotic translation initiation factor 3 subunit c | 10.97 | 1.00E+00 | GRGGMAGGR +2 Methyl x 2 | R2 Methyl:2H(3)13C(1), M5 Label:13C(1)2H(3) |

*some peptides have the same sequence but different charges or additional post-translational modifications (e.g. Phosphorylation)
$localization score of PTMs was not possible to calculate for some peptides identified by the Proteome Discoverer software
red values: hypomethylated in Δ*prmt7*, log2WT/Δ7> 0.6; q-value<0.05
blue values: hypermethylated in Δprmt7, log2WT/Δ7< -0.6; q-value<0.05

**Supplemental Table S2.** Methylpeptides from RNA-binding proteins (RBPs) that are differentially methylated between WT and Δ*prmt7 Leishmania major*. Proteins were considered RBPs if they present an RNA-binding domain or if they are orthologs of a

| Gene ID | Product description | log2FC WT/Δ7 (methylpeptides) | q-value (H&B test) | Methylpeptide sequence* |
|---|---|---|---|---|
| LmjF.27.1680 | hypothetical protein, conserved (NXF1) | 41.28 | 0.00E+00 | SGPGSGGGGVGILGNGNNNGSRGGGR +3 Methyl x 2 |
| LmjF.35.2200 | RNA-binding protein, putative (DRBD2) | 39.62 | 0.00E+00 | HNMAGSAYGRGAPYQVGAAESTEMNGTEPLPKPR +5 Methyl x 1 |
| LmjF.35.2200 | RNA-binding protein, putative (DRBD2) | 35.33 | 0.00E+00 | HNMAGsAYGRGAPYQVGAAESTEMNGTEPLPKPR +4 Methyl x 1 Phos x 1 |
| LmjF.34.1110 | Shwachman-Bodian-Diamond syndrome (SBDS) protein/SBDS protein C-terminal domain contai | 35.19 | 0.00E+00 | LGLDPSHDLQDDsDDDGGRGRGSR +3 Methyl x 2 Phos x 1 |
| LmjF.25.0540 | hypothetical protein SCD6.10 (SCD6) | 35.17 | 0.00E+00 | GRGGRAAAAAPAASATAASSAATR +4 Methyl x 2 |
| LmjF.25.0540 | hypothetical protein SCD6.10 (SCD6) | 34.51 | 0.00E+00 | GRGGRAAAAAPAASATAASSAATR +3 Methyl x 2 |
| LmjF.34.2580 | ALBA-domain protein 3 (Alba3) | 7.15 | 1.15E-02 | GGRGVAAIDRR +3 Methyl x 1 |
| LmjF.25.0290 | RNA-binding protein, putative (eIF4F-like component) | 6.70 | 4.96E-16 | SVARPPPPPPPIGRGR +4 Methyl x 2 |
| LmjF.27.1680 | hypothetical protein, conserved (NXF1) | 6.70 | 1.94E-16 | SGPGSGGGGVGILGNGNNNGSR +2 Methyl x 1 |
| LmjF.27.0130 | WW domain/Zinc finger C-x8-C-x5-C-x3-H type (and similar), putative (ZFP3) | 6.58 | 2.15E-07 | TTHWSIPPTAYEPYYNNGR +3 Methyl x 1 |
| LmjF.27.0130 | WW domain/Zinc finger C-x8-C-x5-C-x3-H type (and similar), putative (ZFP3) | 5.38 | 3.17E-05 | TTHWSIPPTAYEPYYNNGRGR +4 Methyl x 2 |
| LmjF.18.1420 | pumilio protein 2, putative (PUF2) | 4.89 | 3.54E-13 | DNRPGGGVGSSSGGNNNGSGGRAGR +3 Methyl x 1 |
| LmjF.34.1110 | Shwachman-Bodian-Diamond syndrome (SBDS) protein/SBDS protein C-terminal domain contai | 4.44 | 4.14E-11 | LGLDPSHDLQDDsDDDGGRGR +3 Methyl x 1 Phos x 1 |
| LmjF.25.0290 | RNA-binding protein, putative (eIF4F-like component) | 3.92 | 1.67E-12 | SVARPPPPPPPIGR +3 Methyl x 1 |
| LmjF.27.1300 | KH domain containing protein, putative | 3.69 | 2.41E-13 | GGRGGGRGAGAPNGDDAAR +3 Methyl x 2 |
| LmjF.15.0620 | hypothetical protein, conserved | 3.64 | 3.24E-12 | ASSHDNGSNGGAVIGRGGGNAR +3 Methyl x 1 |
| LmjF.24.1180 | Surfeit locus protein 6, putative (SURF6) | 2.77 | 2.44E-07 | GARPFAPQGRGGGASSR +3 Methyl x 1 |
| LmjF.25.1740 | mitochondrial RNA binding complex 1 subunit, putative (RGG3) | 2.34 | 2.94E-06 | SSGGGRGSGSSFGPPVGSHGTGSFGNPR +3 Methyl x 1 |
| LmjF.36.5100 | hypothetical protein, conserved (PUF11) | 1.99 | 1.16E-03 | GGYYQGGGAYGGAGIQQGYPGAAAPISR +3 Methyl x 1 |
| LmjF.25.1740 | mitochondrial RNA binding complex 1 subunit, putative (RGG3) | 1.75 | 1.15E-03 | SSGGGRGSGSSFGPPVGSHGTGSFGNPR +4 Methyl x 1 |
| LmjF.05.0850 | zinc-finger of a C2HC-type, putative | 1.59 | 1.16E-03 | GSGGSGGGGRGGRPAWNSDVEVR +4 Methyl x 1 |
| LmjF.33.1150 | pumilio protein 6, putative (PUF6) | 1.49 | 2.34E-04 | GSYEGEMAGGVTYGRGTGPTAK +3 Methyl x 1 |
| LmjF.25.0540 | hypothetical protein SCD6.10 (SCD6) | 1.44 | 1.49E-03 | AAAAAPAASATAASSAATRGSR +3 Methyl x 1 |
| LmjF.25.0540 | hypothetical protein SCD6.10 (SCD6) | 1.33 | 4.90E-03 | KADTETFGPEMVSSMRGFR +3 Methyl x 1 |
| LmjF.33.0260 | RGG-containing protein 2, putative (RGG2) | 1.25 | 1.52E-04 | DSSDGWGGGRGGGHSGWGGDGGWKDAPTGDR +5 Methyl x 1 |
| LmjF.28.0825 | RNA binding protein rbp16, putative (RBP16) | 0.67 | 2.34E-02 | LPSGPRPPEGSPSGR +3 Methyl x 1 |
| LmjF.33.1150 | pumilio protein 6, putative (PUF6) | 0.64 | 2.02E-02 | YNNNNSAGGGVGGGRVGR +3 Methyl x 1 |
| LmjF.05.0850 | zinc-finger of a C2HC-type, putative | 0.63 | 5.07E-03 | GSGGSGGGGRGGRPAWNSDVEVR +3 Methyl x 1 |
| LmjF.35.3100 | ATP-dependent RNA helicase, putative (DED1) | 0.48 | 2.33E-02 | NYDDGDGGEGGYSRNYDSGYGGGGYR +3 Methyl x 1 |
| LmjF.32.0400 | ATP-dependent RNA helicase HEL67 (DDX3) | 0.40 | 3.12E-03 | KPNVQNQPR +2 Methyl x 1 |
| LmjF.35.1510 | NLI interacting factor-like phosphatase/Zinc finger C-x8-C-x5-C-x3-H type (and similar), putative | 0.13 | 2.20E-03 | ANNSNDHQHQRGGGAR +3 Methyl x 1 |
| LmjF.36.0490 | zinc-finger of a C2HC-type, putative | -0.60 | 2.30E-03 | FSSRGGGGGMGGGGGGGR +2 Methyl x 1 |
| LmjF.32.0850 | polypyrimidine tract-binding protein, putative (DRBD4) | -0.75 | 9.35E-03 | DRGGCGAGSDCAKDSDPAVQR +3 Methyl x 1 |
| LmjF.07.0340 | ATP-dependent RNA helicase DBP2B, putative | -1.05 | 4.65E-03 | SGGGYGGGRGYGGGYGGGR +2 Methyl x 1 |
| LmjF.05.0850 | zinc-finger of a C2HC-type, putative | -1.70 | 1.99E-07 | GAGGGGGRGGGADAAAGAAK +3 Methyl x 1 |
| LmjF.05.0850 | zinc-finger of a C2HC-type, putative | -1.83 | 2.01E-06 | GAGGGGGRGGGADAAAGAAK +2 Methyl x 1 |
| LmjF.32.0840 | hypothetical protein, conserved (DRBD18) | -3.01 | 4.02E-04 | GGRGGSFGGFSQQPTNTISIR +3 Methyl x 1 |
| LmjF.07.0340 | ATP-dependent RNA helicase DBP2B, putative | -3.57 | 7.25E-09 | DGGYGGGYGGGRGGDR +3 Methyl x 1 |
| LmjF.36.6980 | eukaryotic translation initiation factor 3 subunit c | -3.62 | 1.35E-08 | GRGGMAGGR +2 Methyl x 1 |
| LmjF.25.0540 | hypothetical protein SCD6.10 (SCD6) | -3.65 | 7.62E-08 | DPAIVEVHAPARGR +3 Methyl x 1 |
| LmjF.27.1300 | KH domain containing protein, putative | -3.96 | 5.52E-11 | GGGRGAGAPNGDDAAR +2 Methyl x 1 |
| LmjF.34.4290 | nucleolar protein family a, putative | -4.11 | 1.44E-10 | LFTEPPKPR +3 Methyl x 1 |
| LmjF.27.1680 | hypothetical protein, conserved (NXF1) | -4.66 | 4.15E-10 | SGPGSGGGGVGILGNGNNNGSRGGGR +3 Methyl x 1 |
| LmjF.36.5100 | hypothetical protein, conserved (PUF11) | -5.37 | 8.50E-14 | GGPGGLQRGGR +2 Methyl x 1 |
| LmjF.31.0080 | hypothetical protein, conserved (ZC3H34) | -5.82 | 6.18E-07 | GGWYYPSAGSGGYNNMR +2 Methyl x 1 |
| LmjF.01.0210 | CUE domain/Domain of unknown function (DUF1771)/Smr domain containing protein, putative | -3.88 | 3.32E-01 | RGGSGQQDNVAR +3 Methyl x 1 |
| LmjF.01.0210 | CUE domain/Domain of unknown function (DUF1771)/Smr domain containing protein, putative | 0.00 | 1.00E+00 | RGGSGQQDNVAR +2 Methyl x 1 |
| LmjF.02.0660 | Nop14-like family, putative | 0.99 | 1.00E+00 | SGHGRGAGGYTASEMEVR +3 Methyl x 1 |
| LmjF.05.0140 | nucleolar RNA helicase II, putative | -2.30 | 1.68E-01 | GGRGGFNYGGGR +2 Methyl x 1 |
| LmjF.05.0140 | nucleolar RNA helicase II, putative | -0.62 | 8.17E-01 | GGFNYGGGRGGFGGYGGNR +3 Methyl x 1 |
| LmjF.05.0850 | zinc-finger of a C2HC-type, putative | 2.27 | 8.68E-02 | SGGGGGGRGAGGGGGRGGGADAAAGAAK +4 Methyl x 2 |
| LmjF.05.0850 | zinc-finger of a C2HC-type, putative | 0.01 | 3.46E-01 | SGGGGGGRGAGGGGGRGGGADAAAGAAK +3 Methyl x 2 |
| LmjF.05.0850 | zinc-finger of a C2HC-type, putative | 8.42 | 4.62E-01 | GSGGSGGGGRGGRPAWNSDVEVR +2 Methyl x 1 |
| LmjF.05.0850 | zinc-finger of a C2HC-type, putative | 8.04 | 1.06E+00 | GPAASSSPAAGASGVLGTSAQGRGYGsDNEEAGNNAYMPQQPSSAVPR +4 Methyl x 1 Phos x 1 |
| LmjF.07.0340 | ATP-dependent RNA helicase DBP2B, putative | -0.67 | 2.61E-01 | SGGGYGGGRGYGGGYGGGR +3 Methyl x 1 |
| LmjF.07.0340 | ATP-dependent RNA helicase DBP2B, putative | -0.90 | 8.65E-01 | SGGGYGGGRGYGGGYGGGR +3 Methyl x 2 |
| LmjF.07.0870 | splicing factor ptsr1-like protein | -0.19 | 8.54E-01 | RGGYHGSDR +2 Methyl x 1 |
| LmjF.11.0600 | hypothetical protein, conserved | 0.00 | 1.00E+00 | YSEGEHGGAGSRGHGQR +3 Methyl x 1 |
| LmjF.13.0880 | hypothetical protein, conserved | -0.10 | 2.77E-01 | RGMGSHSQSQQQSR +2 Methyl x 1 |
| LmjF.13.0880 | hypothetical protein, conserved | -0.14 | 1.09E+00 | RGMGSHSQSQQQSR +3 Methyl x 1 |
| LmjF.15.0360 | RNA pseudouridylate synthase, putative | 0.53 | 1.00E+00 | QLTRGGGASSLGAWAAALQPVSGVGAAQPAR +3 Methyl x 1 |
| LmjF.15.1380 | nucleolar RNA binding protein, putative | 0.38 | 3.32E-01 | MNTSGFNNDRGGRGGGGGFK +3 Methyl x 2 |
| LmjF.15.1380 | nucleolar RNA binding protein, putative | 1.16 | 7.76E-01 | GGRGGGGGFK +2 Methyl x 1 |
| LmjF.15.1380 | nucleolar RNA binding protein, putative | 2.07 | 1.05E+00 | MNTSGFNNDRGGR +3 Methyl x 1 |
| LmjF.16.1230 | hypothetical protein, conserved | 0.10 | 3.95E-01 | GSRPASPSGVASGSRGGR +3 Methyl x 1 |
| LmjF.16.1230 | hypothetical protein, conserved | -0.40 | 9.47E-01 | GSRPASPSGVASGSRGGR +4 Methyl x 1 |
| LmjF.16.1230 | hypothetical protein, conserved | 0.12 | 1.09E+00 | GKGSRPASPSGVASGSRGGR +3 Methyl x 1 |
| LmjF.17.0550 | RNA-binding protein, putative | 0.36 | 8.57E-02 | EGRGMSRGNNNNSR +3 Methyl x 2 |
| LmjF.17.0550 | RNA-binding protein, putative | 4.31 | 1.20E-01 | GMSRGNNNNSRGAGGNHR +3 Methyl x 2 |
| LmjF.17.0550 | RNA-binding protein, putative | 0.00 | 1.00E+00 | GRGGAHLQPQQNFQQPQQYQQHQLHPPLPPPPPR +5 Methyl x 1 |
| LmjF.18.0800 | Ribosomal protein S8, putative | 0.01 | 4.20E-01 | VYGGGATTAFVPGSSSSSSSSSSTASVGADYLRGR +3 Methyl x 1 |
| LmjF.18.1240 | pre-RNA processing PIH1/Nop17, putative | 1.12 | 4.82E-01 | DRLNSAAAASSGRGGGSGEAAAR +4 Methyl x 1 |
| LmjF.18.1240 | pre-RNA processing PIH1/Nop17, putative | 1.03 | 8.32E-01 | LNSAAAASSGRGGGSGEAAAR +3 Methyl x 1 |
| LmjF.18.1420 | pumilio protein 2, putative (PUF2) | 0.51 | 8.63E-01 | AGRGGRGGNNNNNSSNNSNNQHSDGK +3 Methyl x 2 |
| LmjF.18.1420 | pumilio protein 2, putative (PUF2) | 10.97 | 1.00E+00 | DNRPGGGVGSSSGGNNNGSGGRAGRGGR +3 Methyl x 2 |
| LmjF.18.1420 | pumilio protein 2, putative (PUF2) | 10.97 | 1.00E+00 | DNRPGGGVGSSSGGNNNGSGGRAGRGGR +4 Methyl x 3 |
| LmjF.18.1420 | pumilio protein 2, putative (PUF2) | 10.97 | 1.00E+00 | DNRPGGGVGSSSGGNNNGSGGRAGRGGR +4 Methyl x 2 |
| LmjF.18.1420 | pumilio protein 2, putative (PUF2) | 0.00 | 1.00E+00 | DNRPGGGVGSSSGGNNNGSGGR +3 Methyl x 1 |
| LmjF.18.1420 | pumilio protein 2, putative (PUF2) | 0.00 | 1.00E+00 | DNRPGGGVGSSSGGNNNGSGGRAGR +3 Methyl x 2 |
| LmjF.18.1420 | pumilio protein 2, putative (PUF2) | 0.00 | 1.00E+00 | GGRGGNNNNNSSNNSNNQHSDGKNNAMR +4 Methyl x 1 |
| LmjF.18.1420 | pumilio protein 2, putative (PUF2) | 0.00 | 1.03E+00 | DNRPGGGVGSSSGGNNNGSGGRAGR +4 Methyl x 1 |
| LmjF.19.0060 | 40S ribosomal protein S2 | 0.15 | 1.00E+00 | GRGGPGEEKEWVPCTK +3 Methyl x 1 |
| LmjF.19.1020 | tRNA pseudouridine synthase A-like protein | -2.13 | 4.55E-01 | AAASSTGSGRGGPVVSSAYSAEHEPLTSHGLR +4 Methyl x 1 |
| LmjF.19.1020 | tRNA pseudouridine synthase A-like protein | 0.32 | 8.22E-01 | AAASSTGSGRGGPVVSSAYSAEHEPLTSHGLR +5 Methyl x 1 |
| LmjF.25.0290 | RNA-binding protein, putative (eIF4F-like component) | -0.12 | 4.32E-01 | SVARPPPPPPPIGRGR +3 Methyl x 2 |
| LmjF.25.0290 | RNA-binding protein, putative (eIF4F-like component) | 10.97 | 1.00E+00 | SVARPPPPPPPIGRGRGVGCMTNPPAPISDSGLAIPVPASR +5 Methyl x 2 |
| LmjF.25.0290 | RNA-binding protein, putative (eIF4F-like component) | 10.97 | 1.00E+00 | SVARPPPPPPPIGRGRGVGCMTNPPAPISDSGLAIPVPASR +5 Methyl x 3 |
| LmjF.25.0290 | RNA-binding protein, putative (eIF4F-like component) | 1.25 | 1.00E+00 | AEQYFVPSPTGPSIPLPRGR +3 Methyl x 1 |
| LmjF.25.0290 | RNA-binding protein, putative (eIF4F-like component) | 0.00 | 1.00E+00 | AEQYFVPSPtGPSIPLPR +2 Methyl x 1 Phos x 1 |
| LmjF.25.0290 | RNA-binding protein, putative (eIF4F-like component) | -4.36 | 1.00E+00 | GRGVGCMTNPPAPISDSGLAIPVPASR +3 Methyl x 1 |
| LmjF.25.0290 | RNA-binding protein, putative (eIF4F-like component) | 0.13 | 1.03E+00 | AEQYFVPSPTGPSIPLPR +2 Methyl x 1 |
| LmjF.25.0540 | hypothetical protein SCD6.10 (SCD6) | 0.83 | 1.19E-01 | DPAIVEVHAPARGRGGRAAAAAPAASATAASSAATR +5 Methyl x 3 |
| LmjF.25.0540 | hypothetical protein SCD6.10 (SCD6) | -0.70 | 1.36E-01 | DPAIVEVHAPARGRGGR +3 Methyl x 2 |
| LmjF.25.0540 | hypothetical protein SCD6.10 (SCD6) | 0.32 | 2.71E-01 | KADTETFGPEMVSSMRGFR +4 Methyl x 1 |
| LmjF.25.0540 | hypothetical protein SCD6.10 (SCD6) | -0.09 | 3.88E-01 | ADTETFGPEMVSSMRGFR +3 Methyl x 1 |
| LmjF.25.0540 | hypothetical protein SCD6.10 (SCD6) | -0.98 | 4.39E-01 | KADTETFGPEMVSSMR +3 Methyl x 1 |
| LmjF.25.0540 | hypothetical protein SCD6.10 (SCD6) | 0.15 | 4.63E-01 | DSSSPQARDPAIVEVHAPARGRGGR +4 Methyl x 2 |
| LmjF.25.0540 | hypothetical protein SCD6.10 (SCD6) | 1.36 | 8.22E-01 | DSsSPQARDPAIVEVHAPARGRGGR +4 Methyl x 2 Phos x 1 |
| LmjF.25.0540 | hypothetical protein SCD6.10 (SCD6) | -1.32 | 8.32E-01 | SFYDEAQPYSRGGGGGR +3 Methyl x 1 |
| LmjF.25.0540 | hypothetical protein SCD6.10 (SCD6) | -4.77 | 8.35E-01 | DPAIVEVHAPARGRGGR +3 Methyl x 3 |
| LmjF.25.0540 | hypothetical protein SCD6.10 (SCD6) | 1.13 | 8.43E-01 | DSSSPQARDPAIVEVHAPARGRGGR +4 Methyl x 3 |
| LmjF.25.0540 | hypothetical protein SCD6.10 (SCD6) | 10.97 | 1.00E+00 | GGYGGRGGSGNYR +3 Methyl x 1 |
| LmjF.25.0540 | hypothetical protein SCD6.10 (SCD6) | -0.49 | 1.03E+00 | GGYGGRGGSGNYR +2 Methyl x 1 |
| LmjF.25.0540 | hypothetical protein SCD6.10 (SCD6) | -1.18 | 1.07E+00 | GGRAAAAAPAASATAASSAATR +3 Methyl x 1 |
| LmjF.25.0540 | hypothetical protein SCD6.10 (SCD6) | -1.04 | 1.09E+00 | NHRGGYGGR +2 Methyl x 1 |
| LmjF.25.1080 | hypothetical protein, conserved | -1.63 | 8.23E-01 | GGSRGGRGGAGSCER +3 Methyl x 2 |
| LmjF.25.1540 | mitochondrial RNA binding complex 1 subunit, putative | 1.06 | 4.56E-01 | RGRGSYADAVSDGNMNR +3 Methyl x 2 |
| LmjF.25.1540 | mitochondrial RNA binding complex 1 subunit, putative | 4.23 | 1.00E+00 | AIAAQSVGRGGVGLVESAAPPATSAFR +3 Methyl x 1 |
| LmjF.25.1540 | mitochondrial RNA binding complex 1 subunit, putative | -0.38 | 1.00E+00 | AIAAQSVGRGGVGLVESAAPPATSAFR +4 Methyl x 1 |
| LmjF.25.1740 | mitochondrial RNA binding complex 1 subunit, putative (RGG3) | -0.11 | 3.17E-01 | RGGPMGSR +2 Methyl x 1 |
| LmjF.25.1740 | mitochondrial RNA binding complex 1 subunit, putative (RGG3) | -0.04 | 3.43E-01 | HGYGGNNGSRGGGDGAPR +3 Methyl x 1 |
| LmjF.25.1740 | mitochondrial RNA binding complex 1 subunit, putative (RGG3) | 1.10 | 4.03E-01 | HGYGGNNGSRGGGDGAPR +4 Methyl x 1 |
| LmjF.25.1740 | mitochondrial RNA binding complex 1 subunit, putative (RGG3) | -0.06 | 1.03E+00 | HGYGGNNGSRGGGDGAPR +2 Methyl x 1 |
| LmjF.25.1830 | Histone RNA hairpin-binding protein RNA-binding domain containing protein, putative | 0.53 | 1.00E+00 | MRGSNVSGSAR +2 Methyl x 1 |
| LmjF.26.1110 | TROVE domain containing protein, putative | 0.47 | 8.54E-01 | RGGASVAVSATPAPQAATFVEAPR +3 Methyl x 1 |
| LmjF.26.1530 | RNA recognition motif. (a.k.a. RRMRBD, or RNP domain), putative | -0.16 | 5.14E-01 | SHSNQYSQDSGVSSGQGAGMQRGMR +4 Methyl x 1 |
| LmjF.27.0130 | WW domain/Zinc finger C-x8-C-x5-C-x3-H type (and similar), putative (ZFP3) | 0.87 | 7.48E-02 | TTHWSIPPTAYEPYYNNGRGRGAGYR +5 Methyl x 2 |
| LmjF.27.0130 | WW domain/Zinc finger C-x8-C-x5-C-x3-H type (and similar), putative (ZFP3) | -7.63 | 1.00E+00 | TTHWSIPPTAYEPYYNNGRGR +4 Methyl x 1 |
| LmjF.27.1300 | KH domain containing protein, putative | -2.35 | 1.17E-01 | GGNDEGRGGYAPR +2 Methyl x 1 |
| LmjF.27.1300 | KH domain containing protein, putative | -1.53 | 3.34E-01 | RAGPPAGQPHPPR +3 Methyl x 1 |
| LmjF.27.1300 | KH domain containing protein, putative | 0.01 | 6.82E-01 | GGNDEGRGGYAPR +3 Methyl x 1 |
| LmjF.27.1300 | KH domain containing protein, putative | 0.07 | 1.01E+00 | GGGRGAGAPNGDDAAR +3 Methyl x 1 |
| LmjF.28.0825 | RNA binding protein rbp16, putative (RBP16) | 1.24 | 9.75E-01 | VVSWMSGRGFGFIEDNADKK +3 Methyl x 1 |
| LmjF.28.0825 | RNA binding protein rbp16, putative (RBP16) | 10.97 | 1.00E+00 | VVSWMSGRGFGFIEDNADKK +4 Methyl x 1 |
| LmjF.28.0825 | RNA binding protein rbp16, putative (RBP16) | 10.81 | 1.00E+00 | AENVTAPGGGKLPSGPRPPEGSPSGR +4 Methyl x 1 |
| LmjF.28.0825 | RNA binding protein rbp16, putative (RBP16) | 0.00 | 1.00E+00 | VVSWMSGRGFGFIEDNADK +3 Methyl x 1 |
| LmjF.28.0825 | RNA binding protein rbp16, putative (RBP16) | -0.41 | 1.07E+00 | VVSWMSGR +2 Methyl x 1 |
| LmjF.29.0680 | Triple RNA binding domain protein 3 (TRRM3) | -0.53 | 1.55E-01 | RGGHDSSASHHDEEEHDsDEEAGLDSNAAK +4 Methyl x 1 Phos x 1 |
| LmjF.29.0680 | Triple RNA binding domain protein 3 (TRRM3) | -0.43 | 3.16E-01 | GGASASSASAFSSR +2 Methyl x 1 |
| LmjF.29.0680 | Triple RNA binding domain protein 3 (TRRM3) | 0.00 | 1.00E+00 | RGGHDSSASHHDEEEHDsDEEAGLDSNAAK +3 Methyl x 1 Phos x 1 |
| LmjF.29.0680 | Triple RNA binding domain protein 3 (TRRM3) | -0.15 | 1.00E+00 | HAGGGNHGRGGAAAATR +3 Methyl x 1 |
| LmjF.29.0680 | Triple RNA binding domain protein 3 (TRRM3) | 0.14 | 1.04E+00 | HAGGGNHGRGGAAAATR +4 Methyl x 1 |
| LmjF.29.1090 | ribosomal protein L1a, putative | -0.13 | 8.52E-01 | ISGGGTSTSGAGAFGNMCRGGR +3 Methyl x 1 |
| LmjF.29.1090 | ribosomal protein L1a, putative | -1.94 | 1.04E+00 | ISGGGTSTSGAGAFGNMCR +2 Methyl x 1 |
| LmjF.30.0260 | mitochondrial RNA binding protein 1, putative | -3.64 | 6.16E-01 | QQRGGAGCVGAASNSHAAEGFDVAYPVR +4 Methyl x 1 |
| LmjF.30.0780 | mitochondrial oligo_U binding protein TBRGG1, putative | -0.39 | 1.02E+00 | GGYEGRGGGPR +2 Methyl x 1 |
| LmjF.31.0080 | hypothetical protein, conserved (ZC3H34) | -1.99 | 9.73E-01 | HMDGIPPHAYNQPPVNYYQPPQVYPMGNPMYNVPQGYPPAMPGYRGGR +5 Methyl x 1 |
| LmjF.31.0080 | hypothetical protein, conserved (ZC3H34) | 0.00 | 1.00E+00 | GRGGWYYPSAGSGGYNNMR +3 Methyl x 1 |
| LmjF.32.0400 | ATP-dependent RNA helicase HEL67 (DDX3) | -0.16 | 5.30E-01 | QYGGGNRGFNDAQGTYGGGFNR +2 Methyl x 1 |
| LmjF.32.0400 | ATP-dependent RNA helicase HEL67 (DDX3) | 0.02 | 8.53E-01 | NGGGGGFRGGFGGFGGGAR +2 Methyl x 1 |
| LmjF.32.0400 | ATP-dependent RNA helicase HEL67 (DDX3) | 0.32 | 1.00E+00 | QYGGGNRGFNDAQGTYGGGFNR +3 Methyl x 1 |
| LmjF.32.0400 | ATP-dependent RNA helicase HEL67 (DDX3) | -0.74 | 1.03E+00 | NGGGGGFRGGFGGFGGGAR +3 Methyl x 1 |
| LmjF.32.0400 | ATP-dependent RNA helicase HEL67 (DDX3) | 0.07 | 1.05E+00 | SGGFGGGFGGGYGGGYGGRGGYR +3 Methyl x 1 |
| LmjF.32.0620 | hypothetical protein, conserved | -0.31 | 2.77E-01 | GGMGRAGSGR +2 Methyl x 1 |
| LmjF.32.0620 | hypothetical protein, conserved | 6.24 | 1.00E+00 | GPSFVERGAGAR +3 Methyl x 1 |
| LmjF.32.0840 | hypothetical protein, conserved (DRBD18) | 0.51 | 1.00E+00 | GRGGRGGCGLDTMGMQAGYMPFPQVGGMAR +4 Methyl x 2 |
| LmjF.32.0840 | hypothetical protein, conserved (DRBD18) | -7.25 | 1.00E+00 | GGRGGCGLDTMGMQAGYMPFPQVGGMAR +3 Methyl x 1 |
| LmjF.32.0850 | polypyrimidine tract-binding protein, putative (DRBD4) | 0.66 | 1.18E-01 | GGIGGRGSGTGLGAATGFSPFEPSGTTPARGGGASVGR +4 Methyl x 3 |
| LmjF.32.0850 | polypyrimidine tract-binding protein, putative (DRBD4) | -0.42 | 8.29E-01 | GGIGGRGSGTGLGAATGFSPFEPSGTTPAR +3 Methyl x 1 |
| LmjF.32.0880 | 60S ribosomal protein L18a, putative | 0.74 | 7.75E-01 | HYQVVGR +2 Methyl x 1 |
| LmjF.32.2130 | RNA recognition motif. (a.k.a. RRMRBD, or RNP domain), putative | 0.01 | 2.93E-01 | EPYPSLNSSRPPYGPGMGRGGGMR +4 Methyl x 1 |
| LmjF.33.0260 | RGG-containing protein 2, putative (RGG2) | -0.18 | 4.53E-01 | DSSDGWGGGRGGGHSGWGGDGGWKDAPTGDR +3 Methyl x 1 |
| LmjF.33.0260 | RGG-containing protein 2, putative (RGG2) | -1.16 | 8.61E-01 | DSSDGWGGGRGGGHSGWGGDGGWKDAPTGDR +4 Methyl x 1 |

| | | | | |
|---|---|---|---|---|
| LmjF.33.0260 | RGG-containing protein 2, putative (RGG2) | 0.21 | 9.74E-01 | RGRGGSGGGVWGQPAAADEDAWNAAPPSFQPPVR +4 Methyl x 1 |
| LmjF.33.0260 | RGG-containing protein 2, putative (RGG2) | -2.14 | 1.00E+00 | GRGGSGGGVWGQPAAADEDAWNAAPPSFQPPVR +3 Methyl x 1 |
| LmjF.33.1150 | pumilio protein 6, putative (PUF6) | -0.58 | 6.84E-01 | NNYRGGRGAGGMGSMESGSGPNSHR +4 Methyl x 2 |
| LmjF.33.1150 | pumilio protein 6, putative (PUF6) | 0.07 | 8.55E-01 | GGRGAGGMGSMESGSGPNSHR +4 Methyl x 1 |
| LmjF.33.1150 | pumilio protein 6, putative (PUF6) | -0.59 | 8.71E-01 | GGRGAGGMGSMESGSGPNSHR +3 Methyl x 1 |
| LmjF.33.1150 | pumilio protein 6, putative (PUF6) | -0.25 | 8.77E-01 | RYNNNNSAGGGVGGGRVGR +3 Methyl x 1 |
| LmjF.33.1150 | pumilio protein 6, putative (PUF6) | -0.29 | 9.75E-01 | FARGGGIGNINPSGR +2 Methyl x 1 |
| LmjF.33.1150 | pumilio protein 6, putative (PUF6) | 0.29 | 1.00E+00 | FARGGGIGNINPSGR +3 Methyl x 1 |
| LmjF.34.1110 | Shwachman-Bodian-Diamond syndrome (SBDS) protein/SBDS protein C-terminal domain contai | 0.82 | 1.00E+00 | LGLDPSHDLQDDsDDDGGRGRGSR +4 Methyl x 1 Phos x 1 |
| LmjF.34.3815 | ribosomal protein L14, putative | 0.31 | 1.01E+00 | FKGRGGGGEVSR +3 Methyl x 1 |
| LmjF.34.4290 | nucleolar protein family a, putative | 2.10 | 7.26E-01 | GGRGGGGFGGGR +2 Methyl x 1 |
| LmjF.34.4290 | nucleolar protein family a, putative | -5.12 | 8.20E-01 | GGGRGGHMSEPDPPENVEEVGTFMNAAEGELVYK +3 Methyl x 1 |
| LmjF.34.4290 | nucleolar protein family a, putative | -0.65 | 1.00E+00 | GGGRGGHMSEPDPPENVEEVGTFMNAAEGELVYK +4 Methyl x 1 |
| LmjF.34.4290 | nucleolar protein family a, putative | -4.18 | 1.00E+00 | GGRGGGGFGGGR +3 Methyl x 1 |
| LmjF.35.0630 | Nucleoporin NUP65, putative | -0.90 | 8.30E-01 | RRGGLILESPQANVPDSADATESILR +4 Methyl x 1 |
| LmjF.35.0630 | Nucleoporin NUP65, putative | 0.00 | 1.00E+00 | RGGLILESPQANVPDSADATESILR +3 Methyl x 1 |
| LmjF.35.1510 | NLI interacting factor-like phosphatase/Zinc finger C-x8-C-x5-C-x3-H type (and similar), putative | 0.26 | 7.31E-02 | RANNSNDHQHQRGGGAR +4 Methyl x 1 |
| LmjF.35.1510 | NLI interacting factor-like phosphatase/Zinc finger C-x8-C-x5-C-x3-H type (and similar), putative | 2.24 | 4.74E-01 | RANNSNDHQHQRGGGAR +3 Methyl x 1 |
| LmjF.35.2200 | RNA-binding protein, putative (DRBD2) | 2.45 | 8.29E-01 | HNMAGSAYGRGAPYQVGAAESTEMNGTEPLPKPR +4 Methyl x 1 |
| LmjF.35.2200 | RNA-binding protein, putative (DRBD2) | 10.97 | 1.00E+00 | HNMAGSAYGR +2 Methyl x 1 |
| LmjF.35.2200 | RNA-binding protein, putative (DRBD2) | 0.00 | 1.00E+00 | HNMAGSAYGRGAPYQVGAAEsTEMNGTEPLPKPR +3 Methyl x 1 Phos x 1 |
| LmjF.35.3100 | ATP-dependent RNA helicase, putative (DED1) | 0.03 | 8.06E-02 | RRGGGGGGGGGHR +3 Methyl x 3 |
| LmjF.35.3100 | ATP-dependent RNA helicase, putative (DED1) | 0.14 | 6.16E-01 | SGGPGGRGGGGGGGSWGDSRPSAPSGGGR +3 Methyl x 1 |
| LmjF.35.3100 | ATP-dependent RNA helicase, putative (DED1) | 0.22 | 8.48E-01 | SGGPGGRGGGGGGGSWGDSRPSAPSGGGR +4 Methyl x 1 |
| LmjF.35.3100 | ATP-dependent RNA helicase, putative (DED1) | 0.00 | 1.00E+00 | RYYDDEDDYGEGGDRAGGTGNEDDDYEDGGYDAYR +4 Methyl x 1 |
| LmjF.35.3100 | ATP-dependent RNA helicase, putative (DED1) | -0.36 | 1.03E+00 | RGGGVDDGGF +2 Methyl x 1 |
| LmjF.35.3980 | 4E-interacting protein, putative | 0.48 | 2.09E-01 | GGHNSGMNGGGSSSNHPGSSSTPVYGSGGGRGGDNRR +5 Methyl x 1 |
| LmjF.35.3980 | 4E-interacting protein, putative | 1.47 | 2.15E-01 | GGHNSGMNGGGSSSNHPGSSSTPVYGSGGGRGGDNR +4 Methyl x 1 |
| LmjF.35.3980 | 4E-interacting protein, putative | -0.23 | 8.05E-01 | RGGGGGGGNGGRDDSSNSNSVR +4 Methyl x 1 |
| LmjF.35.3980 | 4E-interacting protein, putative | -0.11 | 8.42E-01 | QGRGVTGNGMSTVPPSPPQFQQPPPPR +3 Methyl x 1 |
| LmjF.35.3980 | 4E-interacting protein, putative | 10.97 | 1.00E+00 | RGGGGGGGNGGRDDSSNSNSVR +3 Methyl x 1 |
| LmjF.35.4950 | zinc finger protein family member, putative | -0.42 | 3.85E-01 | GTEGVGNSYYRGAQQHHGYR +4 Methyl x 1 |
| LmjF.35.4950 | zinc finger protein family member, putative | 0.27 | 4.72E-01 | GAQQHHGYRGVGENGSSR +3 Methyl x 1 |
| LmjF.35.4950 | zinc finger protein family member, putative | 0.65 | 8.14E-01 | LSSMERGGMTGVPQHNLLPPR +4 Methyl x 1 |
| LmjF.35.4950 | zinc finger protein family member, putative | -4.78 | 8.52E-01 | LSSMERGGMTGVPQHNLLPPR +3 Methyl x 1 |
| LmjF.35.4950 | zinc finger protein family member, putative | -1.07 | 8.56E-01 | LSSMERGGMTGVPQHNLLPPRAsPASSQPR +4 Methyl x 1 Phos x 1 |
| LmjF.35.4950 | zinc finger protein family member, putative | 0.70 | 1.00E+00 | SPPATSASTTSAPPPALRGLGR +3 Methyl x 1 |
| LmjF.35.4950 | zinc finger protein family member, putative | -0.65 | 1.00E+00 | GATVTGGVHIDSSVGPRGGAAPMSGMGAGAASR +4 Methyl x 1 |
| LmjF.35.4950 | zinc finger protein family member, putative | -0.25 | 1.03E+00 | LSSMERGGMTGVPQHNLLPPR +4 Methyl x 2 |
| LmjF.35.4950 | zinc finger protein family member, putative | -0.05 | 1.03E+00 | SPPATSASTTSAPPPALR +2 Methyl x 1 |
| LmjF.35.5040 | polyadenylate-binding protein 1 (PABP1) | 10.97 | 1.00E+00 | QQLGGRAQGHPMPMPSPQQPQAPAQPQGFATPSAVGFVQATPK +4 Methyl x 1 |
| LmjF.36.0490 | zinc-finger of a C2HC-type, putative | -0.41 | 4.17E-01 | GPARGGLFGATPSGYGDGAGMSAR +3 Methyl x 1 |
| LmjF.36.0490 | zinc-finger of a C2HC-type, putative | -0.56 | 6.04E-01 | GPARGGLFGAtPSGYGDGAGMSAR +3 Methyl x 1 Phos x 1 |
| LmjF.36.0490 | zinc-finger of a C2HC-type, putative | -0.56 | 8.08E-01 | FSSRGGGGGMGGGGGGGR +3 Methyl x 1 |
| LmjF.36.1640 | universal minicircle sequence binding protein, putative | 0.73 | 1.22E-01 | CGQEGHLSRDCPSSQGGSRGGYGQK +3 Methyl x 1 |
| LmjF.36.1640 | universal minicircle sequence binding protein, putative | -0.15 | 8.55E-01 | DCPSSQGGSRGGYGQKR +4 Methyl x 1 |
| LmjF.36.1640 | universal minicircle sequence binding protein, putative | -0.25 | 8.81E-01 | DCPSSQGGSRGGYGQKR +3 Methyl x 1 |
| LmjF.36.1640 | universal minicircle sequence binding protein, putative | -0.28 | 9.26E-01 | DCPSSQGGSRGGYGQK +3 Methyl x 1 |
| LmjF.36.1640 | universal minicircle sequence binding protein, putative | -0.26 | 1.01E+00 | DCPSSQGGSRGGYGQK +2 Methyl x 1 |
| LmjF.36.1925 | 60S ribosomal protein L37a | -0.19 | 1.08E+00 | TVAGGAYTLSTPNNSTVR +2 Methyl x 1 |
| LmjF.36.2130 | ATP-dependent RNA helicase DBP2A, putative | -0.68 | 2.91E-01 | NYSPFSGFSTTSRGGGGSGAHR +4 Methyl x 1 |
| LmjF.36.2130 | ATP-dependent RNA helicase DBP2A, putative | -0.08 | 8.50E-01 | NYSPFSGFSTTSR +2 Methyl x 1 |
| LmjF.36.2130 | ATP-dependent RNA helicase DBP2A, putative | 0.52 | 1.06E+00 | NYSPFSGFSTTSRGGGGSGAHR +3 Methyl x 1 |
| LmjF.36.5100 | hypothetical protein, conserved (PUF11) | -0.86 | 2.61E-01 | GYSQAYNQSYAARGGYQPTPPQQQAQLQQGYASR +3 Methyl x 1 |
| LmjF.36.5100 | hypothetical protein, conserved (PUF11) | -0.85 | 4.72E-01 | GYSQAYNQSYAAR +2 Methyl x 1 |
| LmjF.36.5100 | hypothetical protein, conserved (PUF11) | -0.26 | 5.02E-01 | GGYQPTPPQQQAQLQQGYASRGGYYQGGGAYGGAGIQQGYPGAAAPISR +5 Methyl x 1 |
| LmjF.36.5100 | hypothetical protein, conserved (PUF11) | -0.83 | 8.03E-01 | GYSQAYNQSYAARGGYQPTPPQQQAQLQQGYASR +4 Methyl x 1 |
| LmjF.36.5100 | hypothetical protein, conserved (PUF11) | 0.46 | 8.50E-01 | GYPPQDGMQGYPQQAGR +2 Methyl x 1 |
| LmjF.36.5100 | hypothetical protein, conserved (PUF11) | -0.39 | 1.00E+00 | GGYQPTPPQQQAQLQQGYASRGGYYQGGGAYGGAGIQQGYPGAAAPISR +4 Methyl x 1 |
| LmjF.36.5100 | hypothetical protein, conserved (PUF11) | -1.32 | 1.02E+00 | GGYGPGYGPGYYPVQQQQGYYNQGYNQNPNYTPAAR +3 Methyl x 1 |
| LmjF.36.6060 | eukaryotic translation initiation factor 4 gamma 4 | -0.09 | 1.18E-01 | YGRGGGPGGLR +2 Methyl x 1 |
| LmjF.36.6060 | eukaryotic translation initiation factor 4 gamma 4 | 0.23 | 1.09E+00 | GPIIGSSNAGGNRGGTRPPSMADR +4 Methyl x 1 |
| LmjF.36.6980 | eukaryotic translation initiation factor 3 subunit c | -2.11 | 4.70E-01 | GRGGMAGGR +2 Methyl x 2 |
| LmjF.36.6980 | eukaryotic translation initiation factor 3 subunit c | 10.97 | 1.00E+00 | GRGGMAGGRGGAGVR +3 Methyl x 2 |

*some peptides have the same sequence but different charges or additional post-translational modifications (e.g. Phosphorylation)

$localization score of PTMs was not possible to calculate for some peptides identified by the Proteome Discoverer software

red values: hypomethylated in Δprmt7, log2WT/Δ7> 0.6; q-value<0.05

blue values: hypermethylated in Δprmt7, log2WT/Δ7< -0.6; q-value<0.05

**Supplemental Table S3**. List of oligonucleotides used in this study for PCR or qRT-PCR.

| Oligo | Oligo sequence | Strategy |
|---|---|---|
| 914.RT-NIMA-R | GCA CAA TCT GGT AGA AGA GA | RIP-qRT-PCR |
| 913.RT-NIMA-F | GAC AAG CTG TTG CTT ATT AT | RIP-qRT-PCR |
| 910.RT-p1s1-301530-R | CCA TCG CGT CAA GCT TTT CC | RIP-qRT-PCR |
| 909.RT-p1s1-301530-F | TGT GCG TGA CTG AGG TTC TC | RIP-qRT-PCR |
| 908.RT-GAPDHcyto-R | TGT CAT GAG GCC CTC GAC TA | RIP-qRT-PCR |
| 907.RT-GAPDHcyto-F | ACG GGC AGC CCA TTA TAT CG | RIP-qRT-PCR |
| 906.RT-amastin340500-R | ACC TTC TCT CCA TGC TGT GC | RIP-qRT-PCR |
| 905.RT-amastin340500-F | TGC CAG AAC AGA GGG CTT TC | RIP-qRT-PCR |
| B3-18S-2 | TCC TTG AAG AAT GCC TTC GC | RIP-qRT-PCR |
| F3-18S-2 | AAC CTC GGT TCG GTG TGT | RIP-qRT-PCR |
| Lmex NMT qPCR For | GCCAAAGACGGTGGCCGATA | RIP-qRT-PCR |
| Lmex NMT qPCR Rev | GGCGTCCACCACTCAAATGT | RIP-qRT-PCR |
| 800.5FLR-Alba20-F | ACA TCC TTC CTG TTC CCC CA | Confirm HA-Alba3 integration |
| 787.3FLR-CCCH0740-R | GCC GGT AGA CCC TCT TCC AT | Confirm HA-Torus integration |
| 786.3FLR-Alba20-R | AAA TAG CCA GCC CCT CCC C | Confirm HA-Alba3 integration |
| 784.3FLR-HEL67-R | ACG TGT TCC CTC CTG CCT TA | Confirm HA-DDX3 integration |
| 783.3FLR-RBP16-R | GCC CTC TCA CTC GTC CTC TT | Confirm HA-RBP16 integration |
| 799.Alba20-HindIII-R | TGA AAG CTT CTA GTT CTC GCG GTC ATC | HA-tagging - Alba3 |
| 798.Alba20-noATG-NheI-BamHI-F | TGA GCT AGC GGA TCC CCT CCA CGT CC[illegible] | HA-tagging - Alba3 |
| 5'FLR-CCCH0740-SfiIA-F | TGA GGCCACCTAGGCC GTTCGACGTAGC[illegible] | HA-tagging - Torus |
| 5'FLR-CCCH0740-SfiIB-R | TGA GGCCACGCAGGCC CGTCGTCTTCGTT | HA-tagging - Torus |
| CCCH0740-noATG-SfiIC-F | TGA GGCCGCTGGGGCC TTCCCCATGGAT[illegible] | HA-tagging - Torus |
| 821.CCCH0740-SfiID-OK-R | TGA GGC CTG ACT GGC CCT ACT CGA TC[illegible] | HA-tagging - Torus |
| 5'FLR-Hel67-SfiIA-F | TGA GGCCACCTAGGCC GGGGAGGAAAA[illegible] | HA-tagging - DDX3 |
| 5'FLR-Hel67-SfiIB-R | TGA GGCCACGCAGGCC GATTCGTGCTTA[illegible] | HA-tagging - DDX3 |
| Hel67-noATG-SfiIC-F | TGA GGCCGCTGGGGCC TATAAGAATCAG[illegible] | HA-tagging - DDX3 |
| Hel67-SfiID-R | TGA GGCCTCAGTGGCC CTACTGACCAAA[illegible] | HA-tagging - DDX3 |
| 5'FLR-RBP16-SfiIA-F | TGA GGCCACCTAGGCC CATTCCTCTCGGT[illegible] | HA-tagging - RBP16 |
| 5'FLR-RBP16-SfiIB-R | TGA GGCCACGCAGGCC GACTGCTTGAAA[illegible] | HA-tagging - RBP16 |
| RBP16-noATG-SfiIC-F | TGA GGCCGCTGGGGCC TTCCGTGTTTCCT | HA-tagging - RBP16 |
| RBP16-SfiID-R | TGA GGCCTCAGTGGCC CTAGAACTCATCG | HA-tagging - RBP16 |
